# Three-dimensional spatio-angular fluorescence microscopy with a polarized dual-view inverted selective-plane illumination microscope (pol-diSPIM)

**DOI:** 10.1101/2024.03.09.584243

**Authors:** Talon Chandler, Min Guo, Yijun Su, Jiji Chen, Yicong Wu, Junyu Liu, Atharva Agashe, Robert S. Fischer, Shalin B. Mehta, Abhishek Kumar, Tobias I. Baskin, Valentin Jamouilĺe, Huafeng Liu, Vinay Swaminathan, Amrinder Nain, Rudolf Oldenbourg, Patrick La Rivère, Hari Shroff

**Author notes:** These authors contributed equally to this work (Talon Chandler, Min Guo).

## Abstract

Polarized fluorescence microscopy is a valuable tool for measuring molecular orientations, but techniques for recovering three-dimensional orientations and positions of fluorescent ensembles are limited. We report a polarized dual-view light-sheet system for determining the *three-dimensional orientations* and diffraction-limited positions of ensembles of fluorescent dipoles that label biological structures, and we share a set of visualization, histogram, and profiling tools for interpreting these positions and orientations. We model our samples, their excitation, and their detection using coarse-grained representations we call *orientation distribution functions* (ODFs). We apply ODFs to create physics-informed models of image formation with spatio-angular point-spread and transfer functions. We use theory and experiment to conclude that light-sheet tilting is a necessary part of our design for recovering all three-dimensional orientations. We use our system to extend known two-dimensional results to three dimensions in FM1-43-labelled giant unilamellar vesicles, fast-scarlet-labelled cellulose in xylem cells, and phalloidin-labelled actin in U2OS cells. Additionally, we observe phalloidin-labelled actin in mouse fibroblasts grown on grids of labelled nanowires and identify correlations between local actin alignment and global cell-scale orientation, indicating cellular coordination across length scales.

## 1 Introduction

Measuring the orientation of fluorescent molecules can provide valuable insights into architecture, order, and dynamics in biology and material science [1]. By tagging a biological structure with a fluorescent reporter that rotates along with the structure of interest, biologists can deduce biophysical dynamics by measuring the orientation of the fluorescent molecule. A large fraction of fluorescent reporters absorb and emit light via an electronic dipole moment, i.e. in a polarized anisotropic pattern, so biologists can use optical microscopy to examine a fluorophore’s excitation and emission patterns to draw conclusions about the fluorescent reporter’s orientation.

Many techniques make *ensemble measurements* of diffraction-limited regions. By making multiple measurements of the same region under variably polarized illumination and/or detection, then calculating each region’s *fluorescence anisotropy* [2, 3], researchers can draw conclusions about membrane labelling [4, 5], septin dynamics [6–8], nuclear pore proteins [9], force orientations [10, 11], and liquid crystals [12, 13]. Recent engineering efforts have improved the spatial resolution [14], signal-to-noise ratio (SNR) [15], and out-of-plane resolution [16] of these ensemble measurements. While some of these studies make assumptions about their samples to estimate three-dimensional orientations, to our knowledge none simultaneously measure orientation and position in three dimensions.

More recently, a collection of *single-molecule measurement* techniques has enabled researchers to measure more parameters, including three-dimensional position, orientation, and rotational dynamics, from a sparse set of emitters. These techniques have been used to distinguish ordered and unordered biomolecular condensates [17], follow DNA conformation changes under tension [18], and capture dynamics of amyloid fibrils [19], myosin [20], membrane [21], and actin [22, 23]. Despite considerable success, these single-molecule efforts face challenges beyond those faced by all fluorescence polarization techniques, with tighter constraints on throughput, SNR, and choice of emitters, constraining their wider adoption.

After surveying the field, we identified an unmet need for measurements that can be used to recover the three-dimensional orientation and position of fluorescent ensembles. We reasoned that a dual-view light-sheet system [24] should provide a excellent platform for measuring the orientation of fluorescent ensembles because of its two excitation and detection arms, enabling diverse illumination and detection polarizations alongside improved axial spatial resolution.

In our initial iteration, we added liquid crystal polarizers to both excitation arms of an existing dual-view light sheet system and attempted to recover the predominant orientation of fluorophores from within each diffraction-limited volume [25]. We found it challenging to merge a single-molecule description of the image formation process with the pixel-wise fluorescence anisotropy methods that are common in ensemble measurements, and we were unable to resolve orientational ambiguities. Inspired by the success of diffusion-tensor magnetic resonance imaging (dMRI) [26] and its high-angular resolution extensions [27], we developed a coarse-grained formalism for spatio-angular fluorescence microscopy [28–30] that we applied to our instrument. The formalism led us to a set of critical engineering insights:

- unlike dMRI, fluorescence microscopes are *spatio-angularly coupled*, i.e. the orientation of molecules affects the spatial point-response function,
- in addition to the widely known spatial diffraction limit, fluorescence microscopes face *angular diffraction limits* set by the physics of dipolar excitation, dipolar emission, and the microscope’s geometry, and
- our initial design had a *hole in its spatio-angular transfer function*, a null function, which caused the observed orientational ambiguities.

In this article we describe the key element of our formalism, the orientation distribution function (ODF), and how we have used it to model our three-dimensional spatio-angular fluorescence microscope. We use ODFs to formulate a forward model that lets us identify spatio-angular holes in our designs. We introduce a solution, *light-sheet tilting*, and we demonstrate that it resolves the ambiguity with the same number of measurements. Subsequently, we describe our observations of membranes, cell walls, and actin in cells grown on a coverslip and on grids of nanowires. We close by inspecting spatio-angular correlations across length scales, and we discuss future directions for this field.

## 2 Results

### 2.1 Orientation distribution functions (ODFs) are coarse-grained models of fluorescent dipoles that label biological structures

**Figure 1(a)** depicts the class of fluorescent objects that we are trying to recover— extended objects containing ensembles of molecules that move and rotate in three dimensions and whose spatial and orientational properties we wish to characterize within diffraction-limited regions. Assuming that each molecule’s excitation and emission dipole moments are aligned—a reasonable approximation for many fluorophores [31]—we can summarize each fluorophore with a single axis. We can further summarize all of the molecules within a diffraction-limited region with an *object ODF*, a spherical function that we depict as a surface with a radius proportional to the number of dipoles oriented along each direction. Fluorescent dipoles that label structures are caught in angular potentials where they rotate during a measurement, contributing to the width and angular-diffusive smoothness of the corresponding ODF. Note that fluorescent dipoles are excited and emit symmetrically about their dipole axes which means that (1) we can depict their dipole moments as axes instead of vectors, and (2) their corresponding ODFs are always antipodally symmetric.

**Fig. 1.**
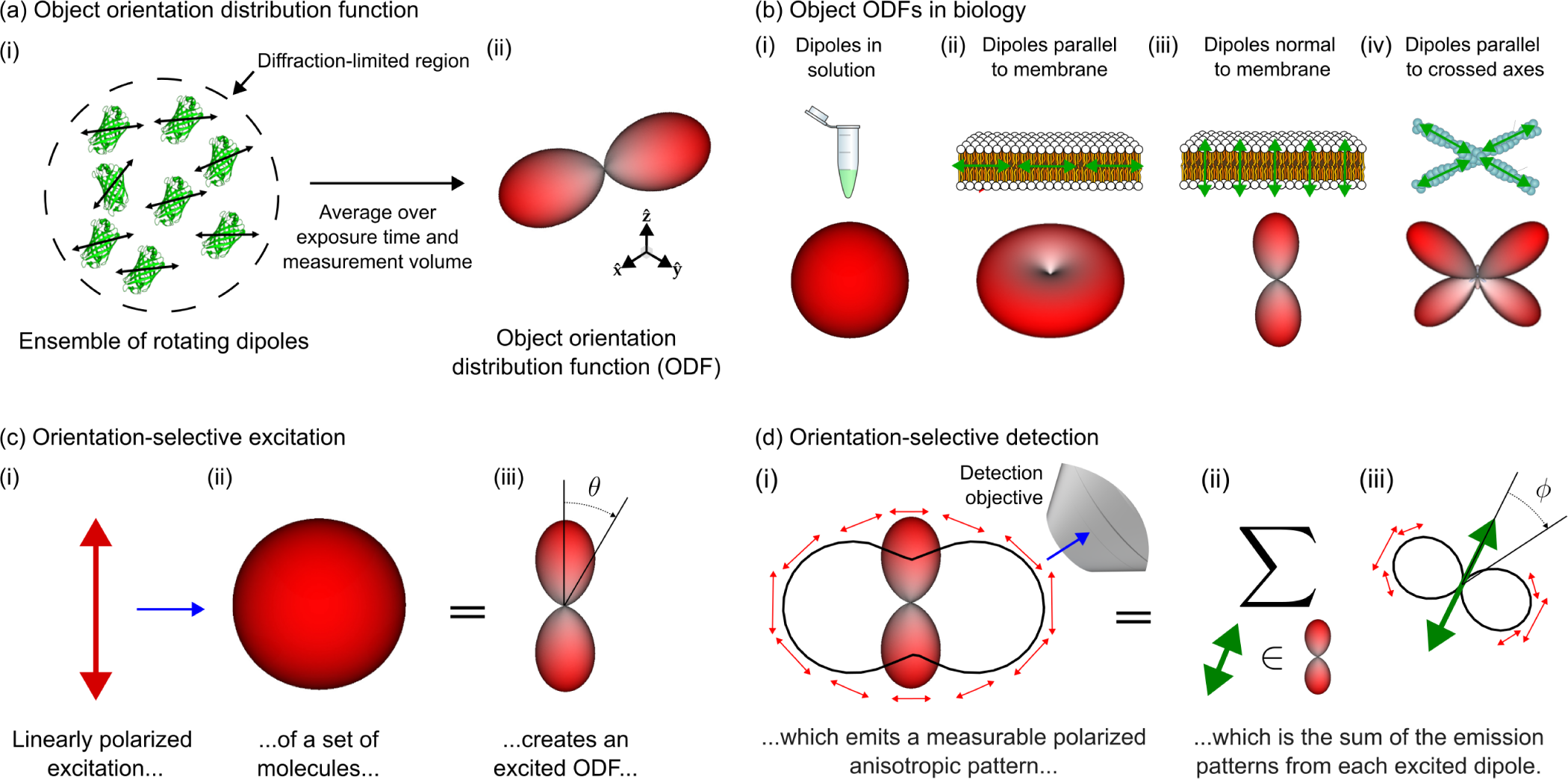
Orientation distribution functions (ODFs) can model ensembles of oriented fluorophores that label biological structures, their excitation, and their detection. **(a)** (i) Fluorescent samples consist of molecules that move and rotate in three dimensions (e.g. green fluorescent protein molecules pictured), and many of the most common fluorescent molecules’ excitation and emission behavior can be described by a single 3D dipole axis (double-sided black arrows overlaid on each molecule). Our instrument excites and measures emissions from diffraction-limited regions that contain many fluorescent molecules (dashed circle), so (ii) we simplify our model of individual emitters to a coarse-grained model called an object orientation distribution function (ODF). An ODF is a spherical function that we depict as a surface with a radius proportional to the number of dipoles in the measurement volume that are oriented along each direction. **(b)** Dipole distributions (top row) can be modelled by object ODFs (bottom row). (i) Fluorescent dipoles in solution typically rotate rapidly during the measurement time of fluorescent microscopes, so the corresponding ODFs are isotropic, depicted as a surface with constant radius. (ii-iv) When fluorescent dipoles (green double-sided arrows) are spatially and rotationally constrained, their corresponding object ODFs report the orientation of labelled biomolecules. **(c)** We can probe an object ODF by exciting a subset of molecules with polarized light. For example, when (i) linearly polarized light (red arrow) illuminates (blue arrow) an (ii) isotropic object ODF, (iii) the resulting subset of excited molecules, which we call an excited ODF, will have a cos^2^ *θ* dependence where *θ* is the angle between the incident polarization and the excitation dipole moment of the individual fluorophores in the distribution. Selectively exciting molecules creates contrast between different object ODFs. **(d)** We can create more contrast by selectively detecting an excited ODF’s emissions. (i) An excited ODF (red glyph) emits a polarized emission pattern (red arrows, perpendicular to the emission direction) that is anisotropic (solid black line, radius is proportional to the emitted power along each direction) which encodes information about the excited ODF. Selectively detecting emissions with an objective (blue arrow) creates contrast between excited ODFs. (ii) The emission pattern in (i) is the sum (Σ) of the emissions from each dipole (green double-sided arrow) in (*∈*) the excited ODF. (iii) Similar to (i), each dipole emits a polarized emission pattern that is anisotropic, with each dipole emitting in a sin^2^ *ϕ* intensity pattern where *ϕ* is the angle between the emission dipole moment and the emission direction.

Object ODFs can describe a wide range of realistic 3D dipole distributions (**Figure 1(b)**). For example, dipoles that rotate freely in solution have an isotropic (spherical) ODF, dipoles that lie flat in the plane of a membrane have a pancake-shaped ODF with more tightly constrained dipoles having correspondingly flatter ODFs, dipoles that are oriented normal to a membrane have a dumbbell-shaped ODF with more tightly constrained dipoles having correspondingly sharper ODFs, and dipoles that lie parallel to multiple axes within a diffraction-limited region have a multi-lobed ODF.

We have developed a *quasi-static* model of fluorescent ensembles, where angular diffusion during a measurement is included within each object ODF, and object ODFs do not change during a measurement. While this model is reasonably accurate for many polarized fluorescence experiments, it ignores effects from fluorescence lifetime, saturation, and spatial diffusion among others. For a complete account of the assumptions leading to our quasi-static model of fluorescent ensembles, we direct readers to **Supplement 5** and [30].

Our primary strategy for generating contrast between different object ODFs is orientation-selective excitation (**Figure 1(c)**). Orientation-selective excitation uses polarized light to excite a subset of an object ODF’s molecules, creating a distribution of excited molecules that we call an excited ODF. Linearly polarized illumination excites an ensemble of dipole absorbers with an efficiency that is proportional to cos^2^ *θ*, where *θ* is the angle between the illumination polarization axis and the excitation dipole moment of individual fluorophores in the distribution. Hence, the excited ODF is the product of the object ODF and the excitation’s cos^2^ *θ* efficiency. When more molecules are excited, more fluorescence can be emitted and collected, so the largest signals will come from object ODFs that are parallel to the illumination polarization. Light is polarized perpendicular to its direction of propagation, so we preferentially excite emitters whose dipole moments are perpendicular to the optical axis of the excitation objective.

Once we have an excited ODF, we can use orientation-selective detection to generate more contrast (**Figure 1(d)**). Ensembles of excited fluorophores emit polarized anisotropic patterns that report their underlying excited ODF. Many instruments use polarization filters/splitters to probe these polarization patterns, but here we rely only on the intensity anisotropy of the emission pattern—a sin^2^ *ϕ* intensity distribution where *ϕ* is the angle between the emission dipole moment and the emission direction. By detecting light with an objective that does not collect light from the entire half space (numerical aperture *<* index of refraction), we preferentially detect emitters whose dipole moments are perpendicular to the optical axis of the detection objective. Therefore, we will measure the largest signals from excited ODFs that lie entirely in the plane perpendicular to the detection objective’s optical axis.

A single ODF can be modelled mathematically as *a function on a sphere*, where the value of the function along a specific direction corresponds to the number of dipoles along that direction. This means that we can write arbitrary ODFs as *f* (**ŝ***_o_*), where **ŝ***_o_*is a coordinate on a two-dimensional sphere S^2^. For example, dipoles in solution (**Figure 1(c)(ii)**) can be represented by a constant-valued function *f* ^(solution)^(**ŝ***_o_*) = *C*, where *C* is constant, and the corresponding excited ODF (**Figure 1(c)(iii)**) can be represented by *f* ^(excited)^(**ŝ***_o_*) = *C|***ŝ***_o_ ·* **p^***|*^2^ = *C* cos^2^ *θ*, where **p^** is the illumination polarization and *θ* is the angle between **ŝ***_o_* and **p^**.

### 2.2 Polarized dual-view light-sheet microscopy enables selective excitation, selective detection, and reconstruction of fluorescent ensembles

In this section we use ODFs as a tool to describe our imaging system, its contrast generation mechanisms, its limits, our reconstruction algorithms, and our visualizations.

**Figure 2(a)** summarizes our dual-view excitation and detection strategy. Our core instrumentation (**Supplement 1.1**) consists of an asymmetric diSPIM frame equipped with a pair of water immersion objectives, each capable of excitation and detection [32].

**Fig. 2.**
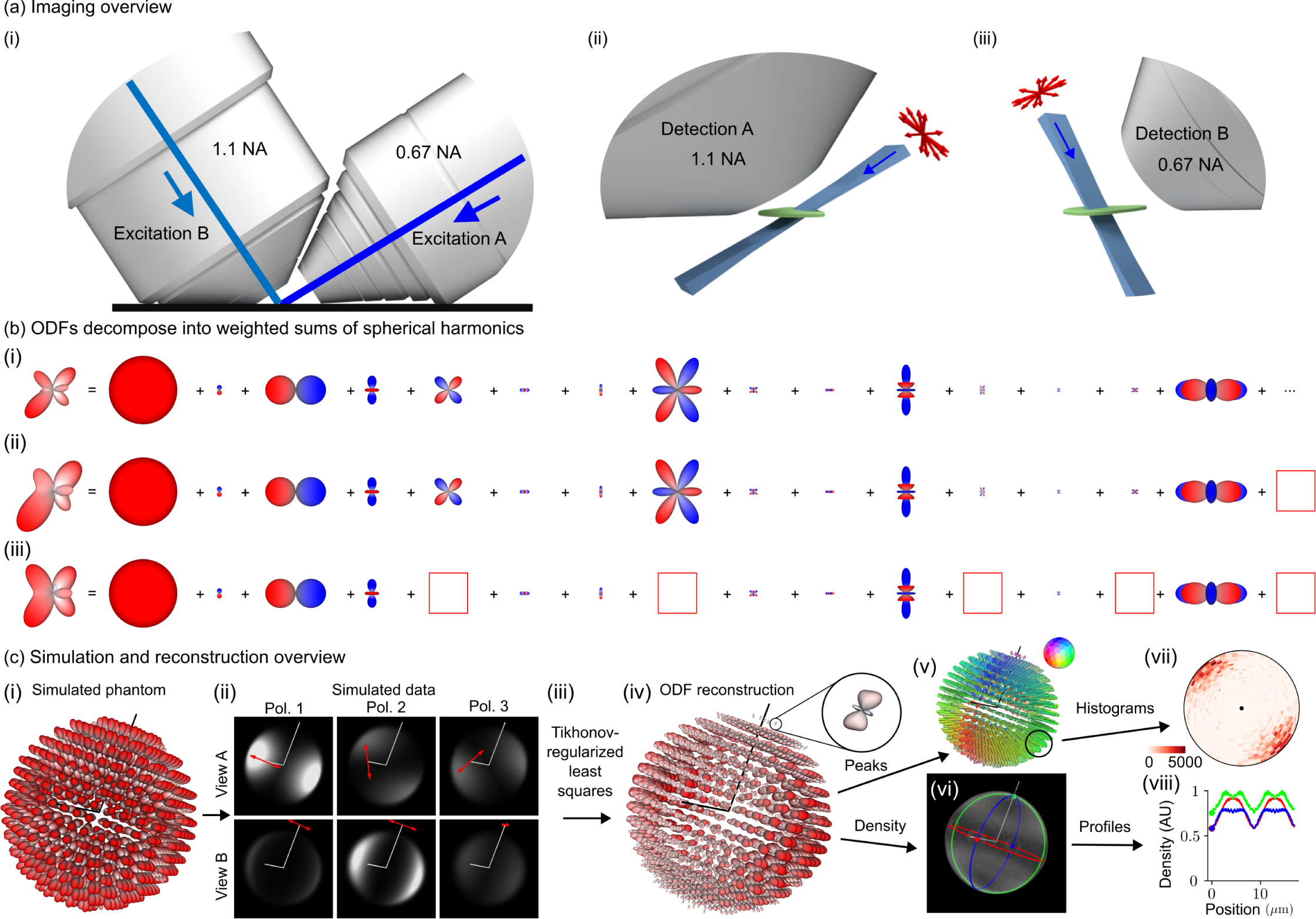
Polarized dual-view inverted selective-plane illumination microscope (pol-diSPIM) data together with a physics-informed reconstruction enables volumetric measurement of three-dimensional orientation distribution functions. **(a)** (i) We imaged our samples with an asymmetric pair of objectives, each capable of excitation and detection. (ii) Illuminating our sample (green) with a light sheet (blue) from the 0.67 numerical aperture (NA) objective and detecting the emitted light from the 1.1 NA objective allows us to make planar measurements of diffraction-limited regions. Modulating the illumination polarization (red arrows) allows us to selectively excite ODFs within each diffraction-limited region, and orthogonal detection allows selective detection. (iii) Excitation from the 1.1 NA objective and detection from the 0.67 NA objective creates additional selective-excitation and selective-detection contrast and complementary spatial resolution. Scanning the sample through these polarized light sheets allows orientation-resolved volumetric acquisitions with more isotropic spatial resolution than detection from a single objective. **(b)** We used spherical harmonic decompositions of ODFs to simulate, reconstruct, and interpret our designs. (i) An example ODF is decomposed into the sum of an infinite number of spherical harmonics with the 15 smoothest non-zero terms shown. (ii) Truncating the infinite sum (red box at right) smooths the ODF while preserving its overall shape, demonstrating the angular resolution our instrument can recover. (iii) Removing more terms (five red boxes) distorts the ODF and increases its symmetry, demonstrating the effect of missing components in the spatio-angular transfer function. **(c)** (i) A simulated phantom of radially oriented ODFs on the surface of a sphere are used to (ii) simulate a dataset. Each volume is simulated with a different illumination objective (rows) and illumination polarization (columns, red arrows indicate polarization, Pol. = Polarization), illustrating how selective excitation and detection (with optical axes indicated by white lines) results in contrast that encodes spatio-angular information. (iii) A physics-informed reconstruction algorithm allows us to recover (iv) ODFs in volumetric regions (inset, a single ODF corresponding to a diffraction-limited volume). We reduce these reconstructions to lower-dimensional visualizations including (v) peak orientations, where the orientation and color of each cylinder indicates the direction along which most dipoles are oriented, and (vi) density, a scalar value indicating the total number of values within each voxel. We further summarize distributions of peak orientations with (vii) angular histograms, where the central dot indicates the viewing axis, and density with (viii) spatial profiles, where the colored profiles correspond to the circumferential profiles in (vi).

We use an excitation-path MEMS mirror (**Supplement 1.2**) to illuminate the sample with a light sheet from the 0.67 numerical aperture (NA) objective, detect the emitted light with the 1.1 NA objective, then scan the sample through the stationary light sheet to acquire an imaging volume. We repeat the acquisition with the objectives’ roles swapped, illuminating with the 1.1 NA objective and detecting with the 0.67 NA objective.

We added a liquid crystal module to both excitation arms (**Supplement 1.2**), enabling our choice of arbitrary transverse polarization illumination. Instead of exploring all possible illumination polarizations, we restricted our possible choices to six linear polarization states, maximizing contrast while still enabling two-fold oversampling of the underlying signals (**Supplement 2.1**).

Our complete acquisition consists of a calibration procedure (**Supplement 3.2**) and the following data acquisition loops from fastest to slowest (**Supplement 3.3**): (*xy*) camera frame, (*z*) stage scan positions, (v) views, (*p*) illumination polarization, (*c*) colors, and (*T*) time points. Our fastest single-time point, single-color acquisition consists of six volumes (three illumination polarizations per view) acquired within 3.6 seconds.

After deskewing (**Supplement 4.1**) and registering (**Supplement 4.2**) the raw data, we collect our irradiance measurements into a single function *g_p_*_v_(**r***_d_*), where *p* is a polarization index, v is a view index, and **r***_d_ ∈* ℝ^3^ is a three-dimensional detector coordinate (**Supplement 5.1**). Next, we model the object that we are trying to estimate, a spatial distribution of ODFs, as a function *f* (**r***_o_,* **ŝ***_o_*), where **r***_o_ ∈* ℝ^3^ is a three-dimensional object-space coordinate and **ŝ***_o_ ∈* S^2^ is an orientation coordinate. Finally, we model the relationship between our data and our object as a shift-invariant integral transform

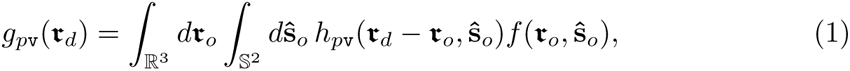

where *h_p_*_v_(**r***_d_ −* **r***_o_,* **ŝ***_o_*) is a *spatio-angular point-response function* (compare with **Supplement 5.8**, see **Supplement 5.1** for indices *p* and *j*). The key features of **Equation 1** are (1) *linearity*: doubling the number of fluorophore doubles the detected irradiance; (2) *3D spatial shift-invariance*: a spatial shift of a fluorophore results in a spatial shift of its irradiance response; and (3) *spatio-angular coupling*: the spatial point-response function depends on the dipole orientation. In other words, *h_p_*_v_(**r***_d_ −* **r***_o_,* **ŝ***_o_*) cannot be factored into a spatial part and an angular part. We assume that the thickness of the light sheet is approximately uniform over the field of view and that the detection-side point-response function is axially Gaussian over the width of the excitation light sheet, assumptions that we find to be true of our light sheets (further assumptions and details are provided in **Supplements 5.2–7**).

Our goal is to estimate the spatial distribution of ODFs, *f* (**r***_o_,* **ŝ***_o_*), from the measured data, *g_p_*_v_(**r***_d_*), but computing and inverting **Equation 1** is extremely computationally expensive. We reformulate **Equation 1** using *spatio-angular transfer functions* to simplify our computations and inversions with the additional benefit of improving our intuition about the imaging system’s limits (**Supplement 6**). We apply *spatial and spherical Fourier transforms* to exploit the symmetries and bandlimits of **Equation 1** to rewrite it as

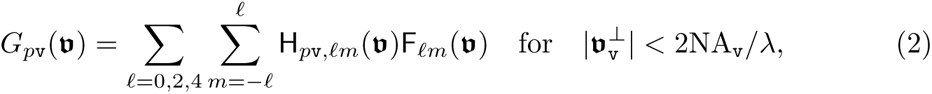

where *G_p_*_v_(**v**) = *F*_R_3 *{g_p_*_v_(**r**)*}* is the *irradiance spectrum*, the 3D spatial Fourier transform of the measured irradiance; H*_p_*_v_*_,ℓm_*(**v**) = *F*_R_3 *_×_*_S_2 *{h_p_*_v_(**r**, **ŝ***_o_*)*}* is the *dipole spatio-angular transfer function*, the 3D spatial and spherical Fourier transform of the spatio-angular point response function; F*_ℓm_*(**v**) = *F*_R_3 *_×_*_S_2 *{f* (**r**, **ŝ***_o_*)*}* is the sample’s *dipole spatio-angular spectrum*; **v** *∈* R^3^ is a three-dimensional spatial-frequency coordinate; **v***^⊥^ ∈* R^2^ is a two-dimensional transverse spatial-frequency coordinate for each view; *ℓ* is the *spherical harmonic band index*, interpretable as the sharpness of an angular component; *m* is the *spherical harmonic intra-band index*, interpretable as the index over all orientation components at a specific angular sharpness *ℓ*; NA*_A_* = 1.1 and NA*_B_* = 0.67; and *λ* is the detection wavelength. The key features of **Equation 2** are (1) *spatial band limits*: transverse spatial frequencies beyond 2NA*/λ* are not detected; (2) *angular discreteness*: instead of the continuous integral over the angular coordinate in **Equation 1** the transfer function formulation uses a discrete sum over spherical harmonic coefficients; and (3) *angular band limits*: angular frequencies from the *ℓ* = 0, 2, and 4 bands are the only terms transmitted. **Figure 2(b)** demonstrates that an arbitrary ODF can be decomposed into a weighted sum of spherical harmonics, that a bandlimited version of an ODF is a smoother version of the original, and that missing intra-band components can distort an ODF.

With an efficient forward model (**Equation 2**) in hand, we used simulations (**Figure 2(c)(i, ii)**) to develop a Tikhonov-regularized least-squares reconstruction algorithm (**Figure 2(c)(iii, iv), Supplements 7.1 and 7.2**). The spatio-angular coupling of the point-response function implies that we need to solve a small inverse problem for each spatial frequency—we cannot solve a small angular problem then solve a separate spatial problem. Therefore, our core algorithm consists of (1) applying a 3D spatial Fourier transform to the deskewed, registered, and calibrated volumes, (2) collecting the Fourier coefficients from each polarization and view into a 6 *×* 1 vector, one vector for each spatial frequency, then (3) multiplying each vector by a precomputed spatial-frequency-specific 15 *×* 6 matrix before (4) applying an inverse 3D Fourier transform and storing the result: a set of 15 spherical harmonic coefficients for each spatial point (**Supplement 7.3 and Supplement Table 4**).

We found that directly visualizing the complete reconstruction, a 3D spatial distribution of ODFs (**Figure 2(c)(iv)**) provided the most information about the sample but was visually overwhelming for most applications. We developed several tools for reducing the visual complexity of the reconstructions (**Supplements 7.4 and 7.5**) including peak-cylinder visualizations where the color and orientation of each cylinder encodes the axis along which most dipoles are oriented (**Figure 2(c)(v)**), histogram visualizations showing peak orientations in larger regions (**Figure 2(c)(vii)**), and scalar metrics including density, proportional to the number of dipoles in each region (**Figures 2(c)(vi, viii)**) and generalized fractional anisotropy (GFA) [33].

### 2.3 Light-sheet tilting enables recovery of all three-dimensional orientations

The angular band limit of our transfer function (**Equation 2**) deserves additional interpretation. Selective excitation with linearly polarized light will generate excited ODFs of the form cos^2^ *θ* multiplied by the object ODF, which means that angular components of degree two, the *ℓ* = 2 spherical harmonics, from the object ODF can be encoded into detected irradiances. Similarly, selective detection will generate irradiance patterns of the form *C*_1_ + *C*_2_ sin^2^ *ϕ*multiplied by the excited ODF (the constants *C*_1_ and *C*_2_ are due to the finite detection numerical aperture), meaning that selective excitation can encode the angular components of degree zero and two, the *ℓ* = 0 and *ℓ* = 2 spherical harmonics. When we combine selective excitation and detection, the imaging system can encode the *ℓ* = 0, 2 and 4 components of the object ODF into the measured irradiance patterns. Similar to spatial structured illumination microscopy (SIM), angularly structured (polarized) illumination aliases high angular frequency components into the detection pass band. Additionally, only even *ℓ* terms are transmitted—antipodally symmetric ODFs mean that ODFs consist of only even-*ℓ* terms.

This argument led us to expect that we could recover all orientations from our sample by exploiting selective excitation and detection, using oversampled illumination polarizations if necessary. We found our intuition to be incorrect, finding that no number of illumination polarizations was enough to recover all orientations from our imaging system. A close inspection of our angular transfer function revealed that we did not properly consider *intra-band angular holes*.

**Figure 3(a)** shows all fifteen *ℓ* = 0, 2 and 4 spherical harmonics along with the four intra-band angular holes in our transfer function. Mathematically, the spherical harmonic functions are grouped into (2*ℓ* +1)-dimensional bands that form rotationally invariant subspaces of the spherical functions, so if a single member of a band is missing we cannot expect rotationally invariant angular resolution. The *ℓ* = 2 and *m* = 1 spherical harmonic, shown in **Figure 3(b)**, is a particularly consequential angular hole because it is the single missing member of the lowest non-zero-order *ℓ* = 2 band, implying that there are some orientations that we cannot recover. We can use an abstract argument to predict this null function by inspection: all multiples of this spherical harmonic are invisible to our imaging system because excitation/detection of positive-valued lobes is always cancelled by excitation/detection of negative-valued lobes.

**Fig. 3.**
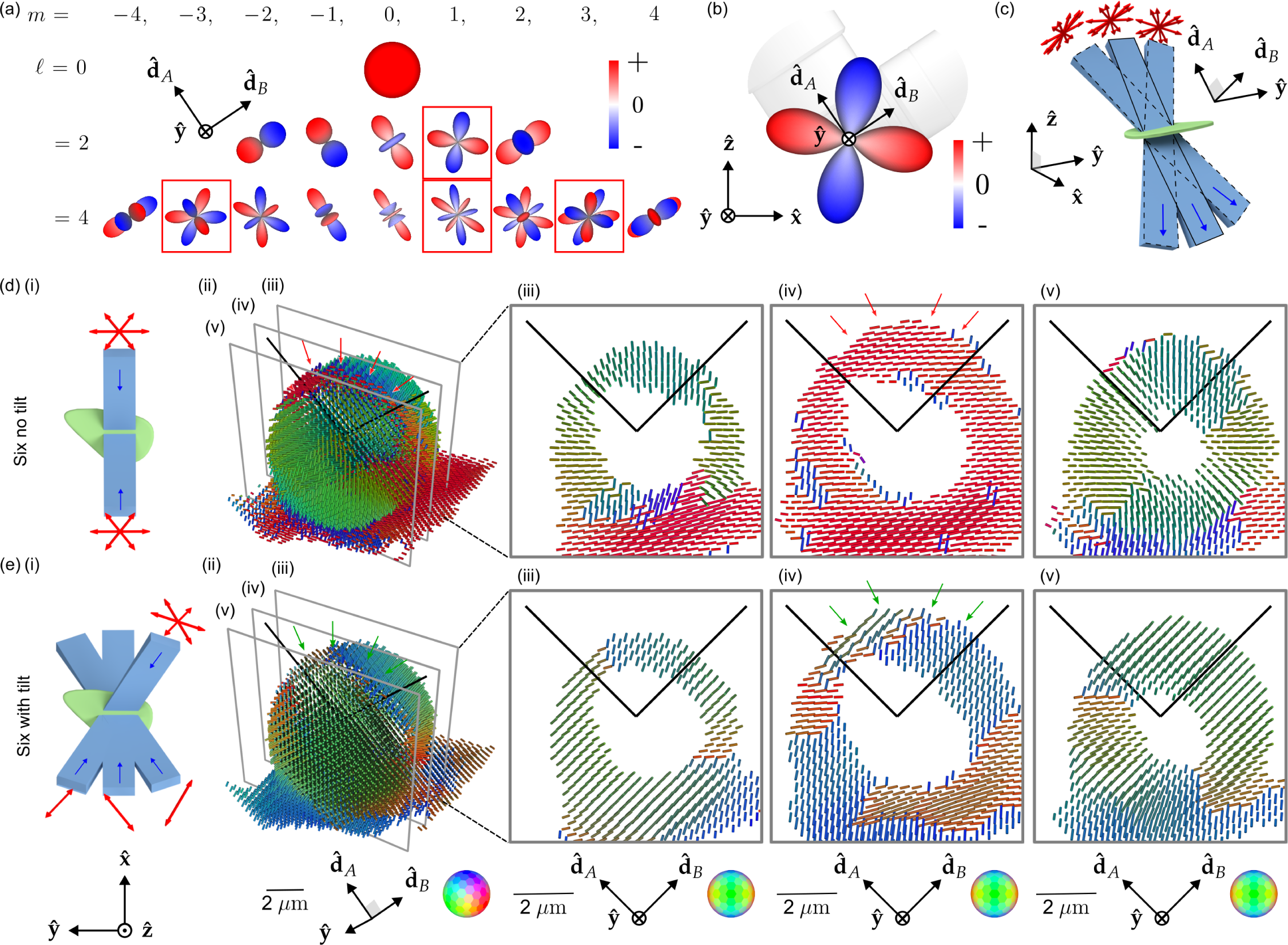
Light-sheet tilting enables experimental recovery of second-order spherical harmonic coefficients and all peak orientations. **(a)** We found that our spatio-angular transfer function had “angular holes” when expressed in a basis of spherical harmonics aligned with the detection axes. Red boxes indicate null functions, spherical harmonics that are not passed to the detected data. **(b)** The second-order angular null function is particularly problematic because it prevents the completion of the *ℓ* = 2 band, causing angular blind spots. Adding any multiple of an angular null function to the object creates identical data, so this angular null function is effectively invisible to our imaging system. **(c)** We added a MEMS mirrors to each excitation arm, enabling light-sheet illumination in the the typical straight-through configuration (blue rectangle with solid outline) and the new tilted configurations (blue rectangles with dashed outlines). Tilting the light sheet makes new polarization orientations (red arrows) accessible while illuminating the same positions in the sample. **(d)** (i) A schematic of our **Six no tilt** acquisition scheme, where the sample (green) is illuminated with light sheets (light blue) propagating parallel to the optical axes of the objectives (dark blue arrows) under three different polarization illuminations per light sheet (red arrows). (ii) Peak cylinder reconstruction from experimental data acquired from a giant-unilamellar vesicle (GUV), where color and orientation encodes the most frequent dipole orientation from within each voxel, spaced by 260 nm. We expect the dipole orientations to be everywhere normal to the GUV, but instead see a red stripe across the top of the reconstructed GUV (see red arrows). (iii-v) Slices through the peak cylinder reconstruction, with incorrect orientations marked with red arrows. **(e)** (i) A schematic of our **Six with tilt** acquisition scheme, which uses a view-asymmetric combination of polarization and tilted light sheets to acquire more angular information from six illumination samples. (ii-v) Peak cylinder reconstruction using tilted light sheets shows recovery of all peak orientations (see green arrows in (ii) and (iv)). Each column of (d) and (e) uses a single coordinate system described below the column where d^^^*_A_* and d^^^*_B_* are the detection optical axes.

We asked what a minimally modified version of our imaging system without an *ℓ* = 2 angular hole would look like, and we used the abstract argument above to propose that *light-sheet tilting* (**Figure 3(c), Supplement 2.2, Supplemental Figures S2 and S3**), enabled by a MEMS mirror placed conjugate to the illumination pupil, would allow us to fill the angular hole in the transfer function while illuminating the same spatial region.

We need to make at least 6 tilting polarization-diverse measurements to recover components from all members of the *ℓ* = 0 and 2 bands. We simulated transfer functions and optimized the condition number of our sampling schemes, searching through *∼* 2 *×* 10^6^ possible sampling schemes to settle on the schemes depicted in **Figures 3(d, e)(i)**. **Supplements 2 and 8** provide more detail about our sampling choices.

Although we measure some components from the *ℓ* = 4 band, we do not recover all of its components. Our imaging system can access all orientations by recovering components from all members of a non-zero band. Recovering the *ℓ* = 4 band would enable better angular resolution, but it would not enable access to more orientations than the *ℓ* = 2 band.

**Figures 3(d, e)** compare a peak-cylinder reconstruction of a GUV with and without light-sheet tilting. The GUV is labelled with FM1-43, a membrane-crossing dye with a dipole transition moment oriented normal to the membrane [34], so we expect a pin-cushion-like reconstruction. **Figures 3(d)(ii, iv)** show incorrect peak cylinder orientations in areas marked with red arrows—peak cylinders lie flat on the surface of the GUV when they should point radially outward. **Figures 3(e)(ii, iv)** show that light-sheet tilting corrects the problem, with continuously radial peak cylinders across the surface of the GUV, highlighted in regions with green arrows. All subsequent data and reconstructions are performed with light-sheet tilting.

### 2.4 pol-diSPIM measurements of fixed samples validate and extend our knowledge of oriented biological structures

Having demonstrated that light-sheet tilting enables recovery of all orientations in GUVs, we proceeded to validate and apply our method to other three-dimensional samples (**Figure 4, Supplemental Movies M1–6**) including GUV, xylem, and actin samples. **Figure 4(a)** shows ODF, peaks, density, and radial profile views of the FM1-43-labelled GUV from **Figure 3**. While the peak-cylinder visualization (**Figure 4(a)(ii)**) is the easiest-to-interpret reconstruction, the ODF reconstruction can reveal the subtlest changes that our system can measure, e.g. two different ODFs can have identical peak cylinders. The density reconstruction (**Figure 4(a)(iii)**, **Supplemental Movies M1–2**) gives a view that is familiar to fluorescence microscopists, with brightness encoding the density. Finally, we measure radial profiles (**Figure 4(a)(iv)**) of the density (**Figure 4(b)(v)**) indicating that the fluorophores are most dense near the GUV’s surface with two-fold variation in intensity due to the non-uniform spatial transfer function of our imaging system—although the transfer function is non-zero for the second-order harmonics, it remains non-uniform. We also measure radial profiles of the generalized fractional anisotropy (GFA) (**Figure 4(a)(vi)**) a scalar measure with GFA = 1 indicating a strongly anisotropic structure and GFA = 0 indicating an isotropic structure. The GFA profiles show similar behavior for all orientations, starting at *∼* 0.6 near the GUV’s center, dipping to *∼* 0.25, then reaching peaks of *∼* 0.75 at the GUV’s surface before dropping again. The behavior near the peak can be interpreted as the increase in oriented structures compared to the random orientations nearby, but the high GFA value in the center of the GUV indicates the effects of noise in low density regions. We consistently observed experiment-to-experiment variation in the value of GFA and orientations in background regions, leading us to only draw conclusions from GFA and peaks in regions with a density above a background threshold.

**Fig. 4.**
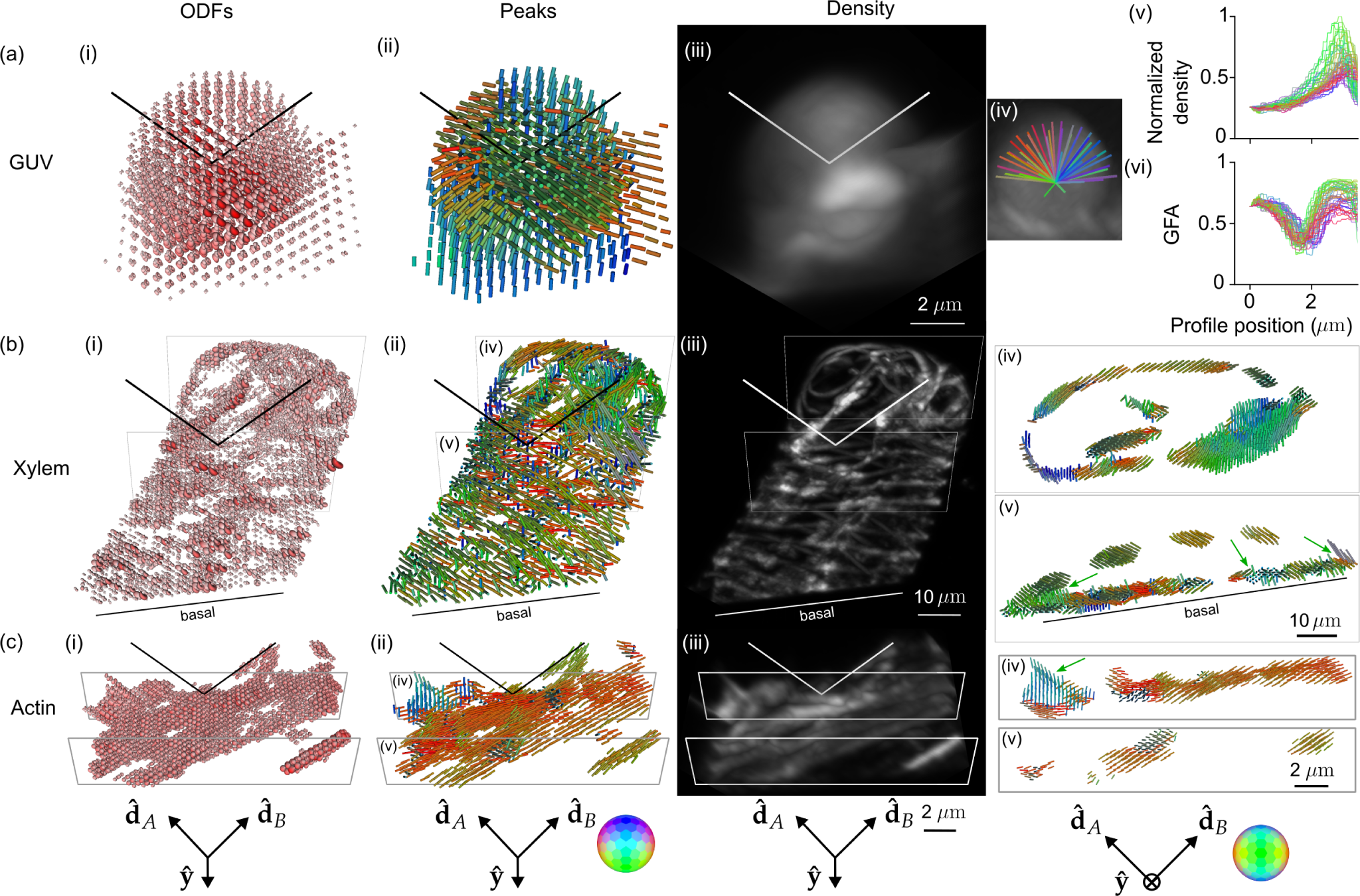
Reconstruction of GUV, xylem, and actin samples validate pol-diSPIM’s accuracy and extend known 2D orientation results to 3D. **(a)** A *∼* 6 *µ*m-diameter GUV labelled with FM1-43 with (i) ODFs and (ii) peak cylinders separated by 650 nm. Radial profiles through the density map (iv) are used to plot density (v) and (vi) generalized fractional anisotropy (GFA) as a function of distance from the center of the GUV. **(b)** A xylem cell with its cellulose labelled by fast scarlet with (i) ODFs and (ii) peak cylinders separated by 1.56 *µ*m. Slices (iv, v) show peak cylinders separated by 650 nm and depict the dipole orientations tracking parallel to the helical cellulose structure, with different orientations on the basal and apical surfaces and spatially merging fibers distinguishable by their orientations (green arrows). **(c)** U2OS cells with actin labelled by phalloidin 488 with (i) ODFs and (ii) peak cylinders separated by 390 nm. Slices (iv, v) show peak cylinders separated by 260 nm and depict out-of-plane (green arrow) and variable in-plane orientations of fixed actin. Each column’s camera orientation and orientation-to-color map is displayed in the bottom row. See also, **Supplemental Movies M1–6**.

**Figure 4(b)** shows fast-scarlet labelled cellulose in a xylem cell (**Supplemental Movies M3–4**), showing dipole orientations parallel to the long axis of the cellulose fibers as expected from 2D studies [35]. We observe 3D orientations tracking the cellulose fibers as they curve in space, spatially-disorganized orientation regions on the basal surface near the cover slip, and the ability to distinguish cellulose fibers that were indistinguishable on the basis of their merged density, but exhibited distinct orientations (**Figure 4(b)(v)**). The disorganization on the basal side is consistent with damage caused by air drying during sample preparation. The thinness of the helices likely indicate a cell in the early stages of differentiation, which would make these cells particularly susceptible to damage via air drying.

**Figure 4(c)** shows Alexa Fluor 488 phalloidin-labelled actin in a U2OS cell (**Supplemental Movies M5–6**), showing dipole orientations parallel to the long axis of the actin filaments as expected [22]. We observe distinct actin filaments, some lying flat in the plane of the coverslip while others reach off the coverslip oriented nearly normal to its surface.

### 2.5 pol-diSPIM measurements of cells grown on nanowires show local-global alignment correlations

Having validated our system’s ability to measure 3D orientations in actin, we used our system to study actin orientations with respect to fixed landmarks by imaging phalloidin-labelled 3T3 mouse fibroblasts grown on nanowire arrays (**Figures 5 and 6**), a model system for studying cell migration.

**Fig. 5.**
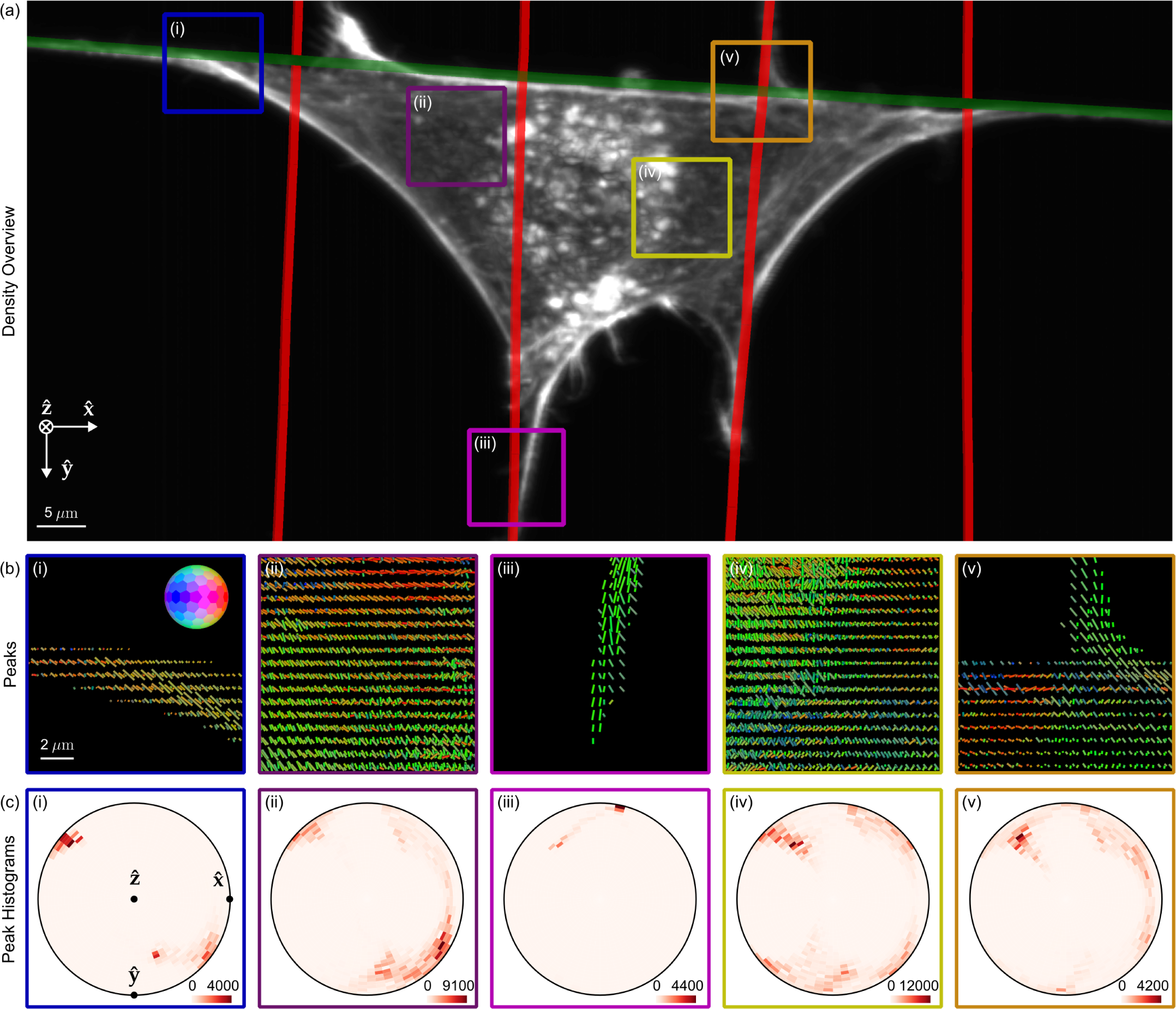
pol-diSPIM measurements of phalloidin-labelled 3T3 mouse fibroblasts grown on nanowires show dipoles oriented parallel to their nearest nanowires and reveal distinct out-of-plane dipole populations across the cell. **(a)** Reconstructed density maximum intensity projection of a cell grown on crossed nanowires, with hand-annotated wires measured from a wire-specific channel highlighted with red and green lines. ROIs (i-iii) are outlined in color and examined in subsequent panels. **(b)** Peak cylinders drawn every 780 nm in regions with total counts *>* 5000, colored by orientation (see inset color hemisphere), with lengths proportional to the maximum diameter of the corresponding ODF. **(c)** Histogram of all peak cylinders with total counts *>* 5000 in each ROI. Bins near the edge of the circle indicate in-plane orientations, bins near the center indicate out-of-plane orientations, and dots mark the Cartesian axes on the histogram.

**Fig. 6.**
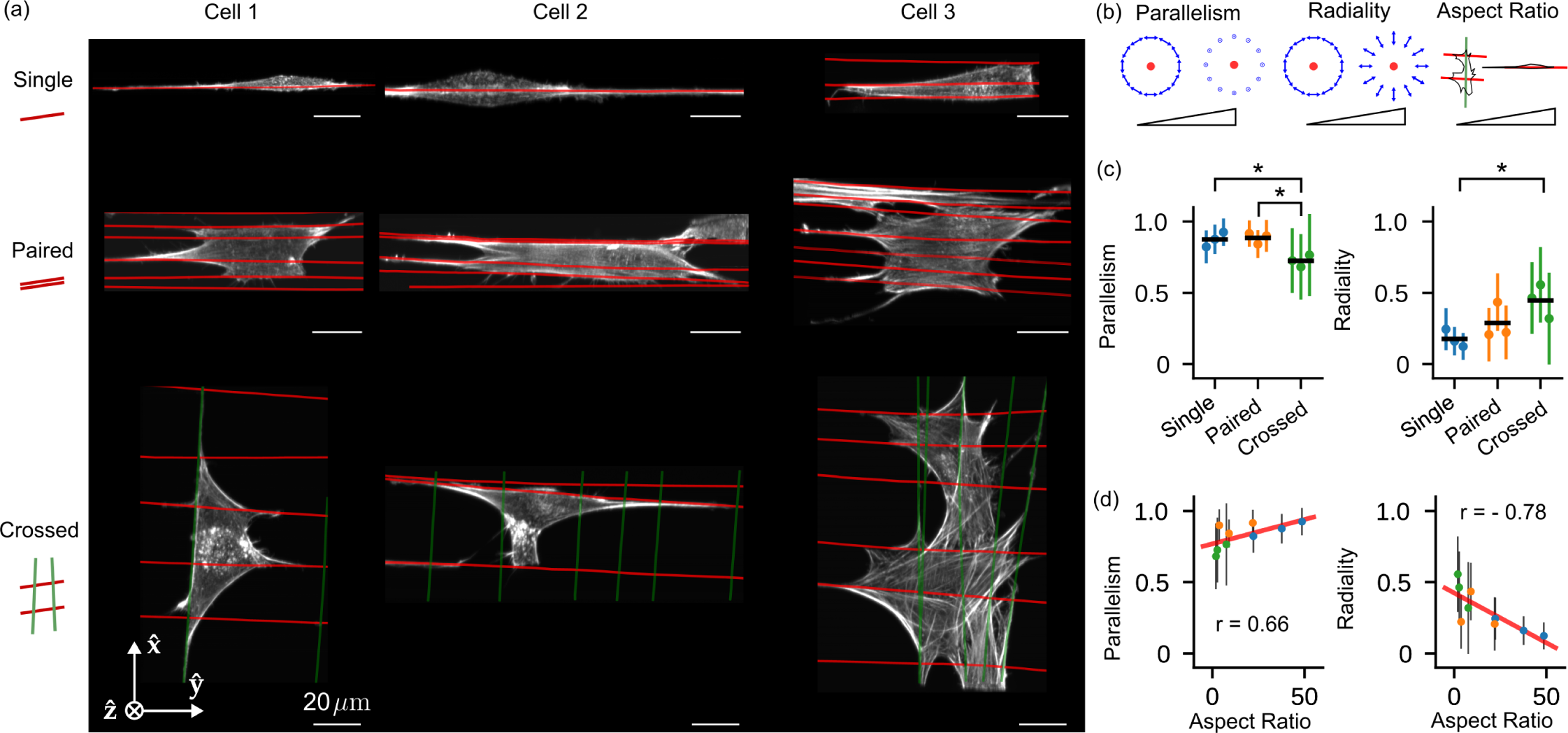
Measurements of 3T3 mouse fibroblasts grown on different nanowire arrangements show correlations between voxel- and cell-scale orientations. **(a)** Reconstructed density maximum intensity projections of three cell repeats (columns) grown on varying nanowire arrangements (rows) named “Single”, “Paired”, and “Crossed” (cartoons at left). Wires are overlaid as red and green lines. **(b)** We collected reconstructed peak directions in voxels that were *<* 5 *µ*m from a wire and had total counts *>* 5000, calculated their parallelism and radiality with respect to their nearest wire (see inset cartoons where the red dot indicates a wire, and blue arrows indicate the neighboring peak directions for strongly parallel and radial peaks), and plotted their mean (dots) and standard deviation (error bars) for each cell and nanowire arrangement (colors). Additionally, we calculated each cell’s “Aspect Ratio”, the ratio of the largest and smallest eigenvalues of the cell’s moment of inertia tensor (with the reconstructed density as a proxy for mass). **(c)** We compared population means (horizontal black lines) with a *t*-test and marked *p <* 0.05 - significant differences with asterisks. **(d)** We compared our local voxel-wise parallelism and radiality metrics to the cell’s global aspect ratio. We found positive and negative correlations between the aspect ratio and the parallelism and radiality, respectively, indicating local-global correlations in cellular behavior. Colored dots match (c), the red line is a linear fit to all nine data points, and the annotated *r* values are Pearson correlation coefficients.

Cells’ immediate environment, the extracellular matrix (ECM), is fibrous, consisting of individual fibrils and bundled fibers ranging in diameter from a few hundred nanometers to several micrometers [36–38], organized in a diverse range of fiber densities, pore sizes, and network architectures, including aligned [39] and crossed-fiber arrangements [40–43]. The complexity of studying cell migration in 3D motivated studies with simpler 1D substrates, which were found to replicate many of the features of 3D matrices while differing from observations on conventional 2D culture systems [44]. Further, in fibrous environments with large pore sizes, cells contact only a few fibers, signifying that cells in vivo can be suspended [45–48] and that imaging cells suspended on 1D and 2D wire arrays can provide biological insight.

Previously, we showed that cellular shapes can be tuned to the underlying fiber network, with actin stress fibers aligning along ECM-fibers in cells attached to single and multiple parallel fibers, and intersecting actin networks in cells attached to a crosshatch network of fibers [49, 50]. In a subsequent study, the underlying actin networks were found to regulate mitotic outcomes through actin retraction fibers connecting the rounded mitotic cell body to interphase adhesion sites [51]. In both studies, while the patterning of actin networks matched the arrangement of underlying ECM networks, it was unclear how far the effect of adhesions on fibers was extended to shape the actin networks.

Here we used pol-diSPIM imaging to investigate long-range adhesion effects in 3D. We deposited suspended fibers *∼* 200 nm in diameter, imaged phalloidin-labelled 3T3 mouse fibroblasts grown on these nanowire arrays, and measured the local orientation of actin networks. We imaged in two channels, a 488 nm channel for wire annotation, and a 561 nm channel for orientation measurements (**Supplement 4.3**).

**Figure 5(a)** shows a density reconstruction of a single cell with ROIs highlighted for closer inspection. The peak cylinders (**Figure 5(b)**) indicate that actin filaments and the dipoles that label them coalign with their nearest nanowires, and the histograms (**Figure 5(c)**) show distinct in-plane and out-of-plane filament populations from within each ROI, including out-of-plane populations that would be invisible to traditional anisotropy measurements. Additionally, we observe more disorder and larger out-of-plane populations for ROIs near the center of the cell (**Figure 5(b-c)(ii, iv)**) than regions near nanowires (**Figure 5(b-c)(i), (iii), (v)**).

To evaluate dipole orientations of F-actin with respect to their nearest wires, we developed a pair of scalar metrics, parallelism and radiality (**Supplement 7.6**), and computed maps of these metrics in regions with a total number of counts across polarizations and views greater than 5000, a threshold that rejects background regions.

We used these metrics to further investigate nine FOVs, with three FOVs for each of three different nanowire arrangements: single, paired, and crossed nanowires (**Figure 6(a)**). Across each FOV we calculated parallelism and radiality for voxels *<* 5 *µ*m from their nearest wire, and we compared these metrics between FOVs and wire arrangements (**Figure 6(b)**). We found the FOVs with the same wire arrangement clustered at distinct parallelism and radiality values and showed no significant differences (colors in **Figure 6(c)**), while FOVs with different wire arrangement showed significantly different values for parallelism and radiality. The crossed wire arrangement showed significantly lower parallelism (0.70 *±* 0.24, mean *±* standard deviation across voxels) than single (0.88 *±* 0.11) and paired (0.89 *±* 0.11) arrangements, indicating local-scale disorder of actin filaments created by the presence of wires in multiple orientations. Consistent with this observation, cells on crossed wires showed significantly increased radiality (0.51 *±* 0.28) over cells on single wires (0.17 *±* 0.13). **Figure 5(v)** shows an ROI consistent with these broader conclusions, where peaks show increased disorder near a wire crossing.

Further, we calculated each FOV’s aspect ratio, a global measure of the effect of nanowire topology on cell polarization, and found that the aspect ratio was positively and negatively correlated with parallelism (*r* = 0.66) and radiality (*r* = *−*0.78), respectively. Both of these correlations are interpretable as evidence for local-global coordination in the architecture of these cells as they grow on their extra-cellular matrix substrates.

Taken together, our measurements demonstrate, for the first time to our knowledge, that the effect of parallel ECM networks in orienting actin is felt over cell-scale distances, and topographical intersections diminish the orientation. Our findings are consistent with previous studies of 2D actin filament orientation and cell shape [52], and we are excited by our system’s ability to extend these findings to 3D.

## 3 Discussion

This work develops a theoretical and experimental bridge between 2D anisotropy measurements and 3D single-molecule orientation measurements. ODFs and the spatio-angular transfer function formalism helped us identify the limits of our imaging system, and we used theory to improve our design with light-sheet tilting. A similar light-sheet tilting scheme was used to reduce absorptive-streaking artefacts [53], but here we use tilting to increase the angular diversity of polarized illumination, allowing us to recover all orientations to extend and draw new conclusions about oriented fluorescent samples in 3D.

Spatio-angular transfer functions point us towards further improvements. Designs that use polarization splitting to make simultaneous selective-detection measurements will give access to the *ℓ* = 4 band without trading off speed, and designs that have a more uniform angular response than our system will improve on our ability to draw conclusions from samples in all orientations. Polarized two-photon excitation and emission [54] provide cos^4^ *θ*-behavior, sharper than the single-photon cos^2^ *θ*-behavior considered here, leading to potential for accessing the *ℓ* = 6 and 8 bands.

Imaging speed limits our ability to draw conclusions from living cells. We measure orientation signals via serial polarized illumination measurements taken over a few seconds, so translational motion on this timescale is indistinguishable from an oriented sample. Although we made many measurements of living cells with our system, we decided to withhold these data from publication as the possibility of spatial motion repeatedly called our conclusions into question. Replacing our polarized illumination strategy with a detection-side polarization splitting strategy provides one path for speed improvement.

We were constrained by a limited palette of fluorescent reporters that rigidly attach to biological structures. Although we are encouraged by recent developments of genetically encoded actin orientation probes [55], we see room for development of bright oriented probes across the spectrum in more biological structures. We see probes as the major limitation in this field, not instrumentation.

Here we made steady-state measurements, considering only quasi-static fluorescent reporters. Time-resolved orientation measurements provide a large set of possibilities for this field [56, 57], and combining time-resolved measurements with reversibly switchable proteins allows measurement of a wide range of reorientation timescales [58], giving access to reorientation timescales of large protein complexes. We are excited for future developments that probe long-timescale reorientation of fluorophores trapped in 3D angular potentials.

Fluorescence anisotropy can be used for homoFRET measurements of molecular binding [59], and an early non-tilting variant of the system described here was used to make such measurements [60]. We hope that future efforts will encode additional physical parameters, such as force and voltage, into the orientation and rotational mobility of fluorescent probes with readouts enabled by systems like the one described here.

## 4 Online Methods

This section describes all sample-preparation protocols. All imaging and analysis details can be found in the supplementary materials.

### 4.1 Bead samples

Glass coverslips (24 mm *×* 60 mm, #1.5, Electron Microscopy Sciences, 63793-01) were cleaned with clean water and coated with 0.1% poly(l-lysine) (Sigma-Aldrich, P8920) for 10 minutes. 100 nm diameter yellow-green beads (Thermo Fisher Scientific, F8803) were diluted *∼* 10^5^-fold, and 20 *µ*L added to the coverslip. After 10 minutes, the coverslip was washed three times with clean water before imaging. Beads were used to obtain measured estimates of the system PSF, which in turn were used to guide the generation of theoretical PSFs.

### 4.2 Fluorescence slides

For system calibration, a fluorescent plastic slide (Chroma, 92001) was carefully cut into small pieces (*∼* 4 *×* 5 mm^2^)) and glued to a glass coverslip (24 mm *×* 60 mm, #1.5, Electron Microscopy Sciences, 63793-01). Then the coverslip was mounted to a chamber (Applied Scientific Instrumentation, I-3078-2460) and imaged with diSPIM objectives to measure fluorescence changes as we varied the excitation modulation.

### 4.3 Giant unilamellar vesicles samples

We prepared giant unilamellar vesicles (GUVs) via electroformation [61, 62]. We coated a coverslip with 20 *µ*L cBSA, waited for *∼* 15 minutes at room temperature for it to dry into a thin layer, then washed three times with distilled water. We mixed 2 *µ*L of FM1-43 (ThermoFisher, a membrane-crossing dye with a dipole transition moment oriented normal to the membrane [63]) and 40 *µ*L of GUV solution in a 1.5 mL tube, transferred the solution to the cBSA coated coverslip, and waited for *∼* 20 minutes for GUVs to settle. Finally, we placed the coverslip in the imaging chamber, filled it with sucrose solution, and waited *∼* 12 hours, covered with a thin film to reduce evaporation, before imaging.

### 4.4 Fixed plant xylem samples

Xylem cells were prepared by inducing tobacco (Nicotiana tabacum) BY-2 cells to differentiate into tracheary elements, as described by Yamaguchi et al. [64]. Briefly, cells were cultured with standard methods for BY-2 [65]. A stable cell line was generated in which a transcription factor (VND7), driven by an inducible promoter (dexamethasone), had been integrated into the genome. Four days after adding 1 *µ*M dexamethasone to the culture, cells were collected, stained for 30 minutes with 0.02% fast scarlet in growth medium, rinsed in growth medium, adhered to poly-L-lysine coated coverslips, and air dried. Fast scarlet binds cellulose in an oriented manner [66].

### 4.5 Fixed U2OS cells with labelled actin

U2OS cells (American Type Culture Collection, HTB-96) were cultured in DMEM media (Lonza, 12-604F) supplemented with 10% FBS (Thermo Fisher Scientific, A4766801) at 37*^◦^*C and 5% CO_2_ on coverslips. Cells were fixed by 2% paraformaldehyde (Electron Microscopy Sciences, 15711) in 1*×* PBS at room temperature for 15 minutes and rinsed three times with 1*×* PBS. Cells were incubated with Alexa Fluor 488 phalloidin (Invitrogen, A12379, 1:50 dilution in 1× PBS) for 1 hour at room temperature and rinsed three times with 1*×* PBS before imaging.

### 4.6 Fiber network manufacturing

Polystyrene fibers were manufactured using the non-electrospinning spinneret-based tunable engineered parameters (STEP) platform as previously reported [67, 68]. Polystyrene of two different molecular weights (Agilent, *M_w_* = 15 *×* 10^6^ g/mol and Polystyrene Standard, *M_w_* = 2.5*×*10^6^ g/mol) was dissolved in xylene (Carolina Chemicals) to form polymeric solutions at 5% (w/w). Additionally, 20 *µ*L of 1 mg/mL of BDP FL Maleimide dye (Lumiprobe) was added to the polymer solutions to get fluorescent fibers. Fibers were spun on hollow 5 *×* 5 mm metal scaffolds. The first layer of fibers deposited were large diameter fibers *∼* 2 *µ*m (*M_w_* = 15 *×* 10^6^ g/mol) followed by an orthogonal layer of 200 nm (*M_w_* = 2.5 *×* 10^6^ g/mol) fibers with spacing varying from 7 to 20 *µ*m to achieve a variety of cell shapes (elongated on single fibers and parallel-shaped cells on two or more fibers) [51, 69, 70]. Additionally, crosshatch networks of 200 nm fiber diameters were also prepared with spacing varying from 7 to 20 *µ*m [50, 71] to achieve polygonal and kite-shaped cells on multiple fibers. The fiber networks were fused at junctions using a custom-built chemical fusing chamber.

### 4.7 Cell culture and seeding on fiber networks

3T3 mouse fibroblasts (ATCC) were grown in Dulbecco’s modified Eagle’s medium (Corning) supplemented with 10% fetal bovine serum (Corning) in T25 flasks (Thermo Scientific). The cells were grown in an incubator kept at 37*^◦^*C and 5% CO_2_. The nanofiber network scaffolds were tacked on a cover glass (VWR, 24 *×* 60 mm No. 1.5) with the help of high-vacuum grease (Dow Corning). Next, the scaffolds were sterilized with 70% ethanol for 10 minutes followed by Phosphate Buffer Solution (PBS) washes (two times). Next, the scaffold was coated with 4 *µ*g/mL bovine fibronectin (Sigma Aldrich) in PBS for at least one hour to promote cell adhesion. Cells were then seeded onto the scaffolds with a seeding density of 300,000 cells/mL and were allowed to spread onto the fibers for a few hours followed by the addition of 3 mL of media. Cells were allowed to further spread for an additional 24 hours before fixation.

### 4.8 Immunostaining cells on fiber networks

Cells were fixed with 4% paraformaldehyde in PBS (Santa Cruz Chemicals) for 15 minutes. The cells were then washed with PBS two times and then permeabilized with 0.1% Triton X-100 solution. Following two PBS washes the cells were blocked with 5% goat serum (Fisher Scientific) for 30 minutes. Next, conjugated antibody Alexa Fluor 568 Phalloidin (1:100, Thermo Fisher) diluted in antibody dilution buffer was added to the cells. After one hour, the cells were washed with PBS (3*×*, 5 minutes each). The sample was then covered in 2 mL of Live Cell Imaging Media (Thermo Fisher) for imaging.

## Acknowledgements

We thank Applied Scientific Instrumentation (ASI) for sharing drawings and assisting with hardware.

## Supplementary information

This manuscript is accompanied by a supplementary document with 9 text supplements, 4 tables, 13 figures, and 6 movies.

## Ethics approval and consent to participate

Not applicable.

## Consent for publication

Not applicable.

## Data availability

Registered data, analysis scripts, reconstructions, and visualizations of all samples described here are available on the BioImage Archive at https://www.ebi.ac.uk/biostudies/bioimages/studies/S-BIAD1055.

## Materials availability

Not applicable.

## Code availability

Pre-processing analysis software is available at https://github.com/eguomin/microImageLib. Reconstruction and visualization software is available at https://github.com/talonchandler/polaris. Several visuals were enabled by Dipy [72].

## Author contributions

T. C. developed theory, analysis, and visualization tools; suggested theory-motivated experimental improvements; performed all post-registration analysis; and drafted the paper, figures, and supplements. M. G. built the microscope, aligned and calibrated its optical system, modified its hardware and software to support light-sheet tilting, acquired all data, and performed all data pre-processing. Y. S., J. C., and V. J. prepared cellular samples. J. L. and H. L. contributed to analysis pipelines. Y. W. contributed to preprocessing software and an early prototype system. A. A. prepared and interpreted nanowire samples. S. B. M. and A. K. contributed to an early prototype of the microscope. T. B. prepared and interpreted xylem samples. R. F., A. N., and V. S. contributed biological context, guided our nanowire investigations, and interpreted nanowire and live-cell samples. R. O. prepared GUV samples and provided experimental, theoretical, and interpretation guidance. P. L. R. and H. S. oversaw the work, contributing to theory, experiment, analysis, visualization, interpretation, and writing. All coauthors contributed to revisions.

