## Supplemental Document for "Three-dimensional spatio-angular fluorescence microscopy with a polarized dual-view inverted selective-plane illumination microscope (pol-diSPIM)"

[1]CZ Biohub SF, San Francisco, 94158, California, USA.
[2]Department of Radiology, University of Chicago, Chicago, 60637, Illinois, USA.
[3]State Key Laboratory of Extreme Photonics and Instrumentation, College of Optical Science and Engineering, Zhejiang University, Hangzhou, 310027, Zhejiang, China.
[4]Laboratory of High Resolution Optical Imaging, National Institute of Biomedical Imaging and Bioengineering, National Institutes of Health, Bethesda, 20892, Maryland, USA.
[5]Advanced Imaging and Microscopy Resource, National Institutes of Health, Bethesda, 20892, Maryland, USA.
[6]Janelia Research Campus, Howard Hughes Medical Institute, Ashburn, 20147, Virginia, USA.
[7]Department of Mechanical Engineering, Virginia Tech, Blacksburg, 24061, Virginia, USA.
[8]Cell Biology and Physiology Center, National Heart, Lung, and Blood Institute, National Institutes of Health, Bethesda, 20892, Maryland, USA.
[9]Bell Center, Marine Biological Laboratory, Woods Hole, 02543, Massachusetts, USA.

[10]Biology Department, University of Massachusetts, Amherst, 01003, Maryland, USA.
[11]Whitman Center, Marine Biological Laboratory, Woods Hole, 02543, Massachusetts, USA.
[12]Department of Molecular Biology and Biochemistry, Simon Fraser University, Burnaby, V5A 1S6, British Columbia, Canada.
[13]Department of Clinical Sciences, Lund University, Lund, SE-221 00, Scania, Sweden.
[14]Wallenberg Centre for Molecular Medicine, Lund University, Lund, SE-221 00, Scania, Sweden.
[15]Department of Biomedical Engineering and Mechanics, Virginia Tech, Blacksburg, 24061, Virginia, USA.

†These authors contributed equally to this work.

### 1 Instrumentation

#### 1.1 Core components

The pol-diSPIM system, shown in **Figure S1(a)**, is built on an asymmetric diSPIM frame [1] equipped with a pair of water-immersion objectives: a 25×, 1.1 NA lens (Nikon, MRD77220, $f = 8$ mm), and a 28.6×, 0.67 NA lens (Special Optics, 54-10-7@488-910nm, $f = 7$ mm). We mounted our samples on glass coverslips (24 × 60 mm, #1.5, Electron Microscopy Sciences, 63793-01) and placed the coverslips in an imaging chamber (Applied Scientific Instrumentation, I-3078-2460). We mounted the imaging chamber to an XY piezo stage (Physik Instrumente, P-545.2C7, 200 $\mu$m × 200 $\mu$m) that we bolted to a motorized XY stage (Applied Scientific Instrumentation, RAMM and MS-2500). We used the motorized stage for coarse sample positioning before imaging, and we used the piezo stage to step our samples through stationary light sheets to create imaging volumes.

#### 1.2 Excitation optics

To excite the sample, we combined 488 nm, 561 nm, and 640 nm lasers (Coherent, OBIS models 1277611, 1280720 and 1185055) with two dichroic mirrors (Semrock, Di02-R488-25x36 and Di01-R488/561-25x36), passed the combined beam through an acousto-optic tunable filter (AOTF, Quanta Tech, AOTFnC-400.650-TN) for power and shuttering control, split the beam into two paths with a 50/50 beam splitter

(Chroma, 21014), then guided these two beams to our excitation arms with single-mode and polarization-preserving fibers (Excelitas Technologies, kineFLEX Fiber Delivery System, 012486). In each arm, we used a pair of MEMS mirrors (Applied Scientific Instrumentation, anti-striping fiber-coupled laser scanner) to scan and tilt the laser beam, **Figure S1(b)**. We used the first MEMS mirror, slightly offset from a conjugate image plane to avoid potential burning of the mirror by a concentrated focus, to tilt the beam in the plane of the sample; and we used the second MEMS mirror, conjugate to the back focal plane of each objective, to scan the beam to create a virtual light sheet in the plane of the sample. See **Figure S2(e, f)** to see how these mirrors affect the light sheet in sample space.

To modulate the excitation polarization, we placed a liquid crystal (LC) module in each arm between a 300 mm tube lens (for excitation path, Applied Scientific Instrumentation, C60-TUBE-E-300) and the objective. Each LC module, **Figure S1(c)**, contains a wire-grid linear polarizer followed by two stacked liquid crystal variable retarders (LCVRs, Meadowlark Optics, LVR-200) assembled in a custom acrylic housing with the first LCVR's slow axis oriented at 45 degrees relative to the linear polarizer's transmission axis and the second LCVR's slow axis parallel to the linear polarizer's transmission axis [2]. The linear polarizer ensures that the beam reaching the LCVRs is polarized with a high purity and fixed orientation, and the LCVR's orientations are chosen so that varying their voltages allows us to illuminate our sample with any polarization state perpendicular to the direction of the beam propagation. We used a four-channel LC digital controller (Meadowlark Optics, D3050) to apply voltages to both LCVRs on both ilumination arms.

We achieved additional polarization modulation by tilting the beam with MEMS mirror 1, see **Figure S1(b)**. Tilting the illumination beam changes the beam's propagation direction and the polarization states that are accessible via the LCVRs, so we varied the first MEMS mirrors and the LCVRs together to explore a large set of possible illumination polarizations. We describe our specific polarization samples in more detail in **Supplement 2**.

##### 1.3 Detection optics

For each arm, we collected fluorescent emissions with an objective, blocked reflections and background with dichroic mirrors (Semrock, Di03-R405/488/561/635-t1-25x36) and a quad-notch filter (Semrock, NF03-405/488/561/635E-25), and imaged with a tube lens (400 mm tube lens for the 1.1 NA detection path, Applied Scientific Instrumentation, C60-TUBE-400, and 200 mm tube lens for the 0.67 NA detection path, Applied Scientific Instrumentation, C60-TUBE-B) onto sCMOS cameras (Hamamatsu, ORCA Flash 4.0 v2). The effective magnifications for the 1.1 NA and 0.67 NA detection paths are 50× and 28.6×, respectively, so the resulting object-space pixel sizes are 6.5 $\mu$m/50 = 130 nm, and 6.5 $\mu$m/28.6 = 227 nm.

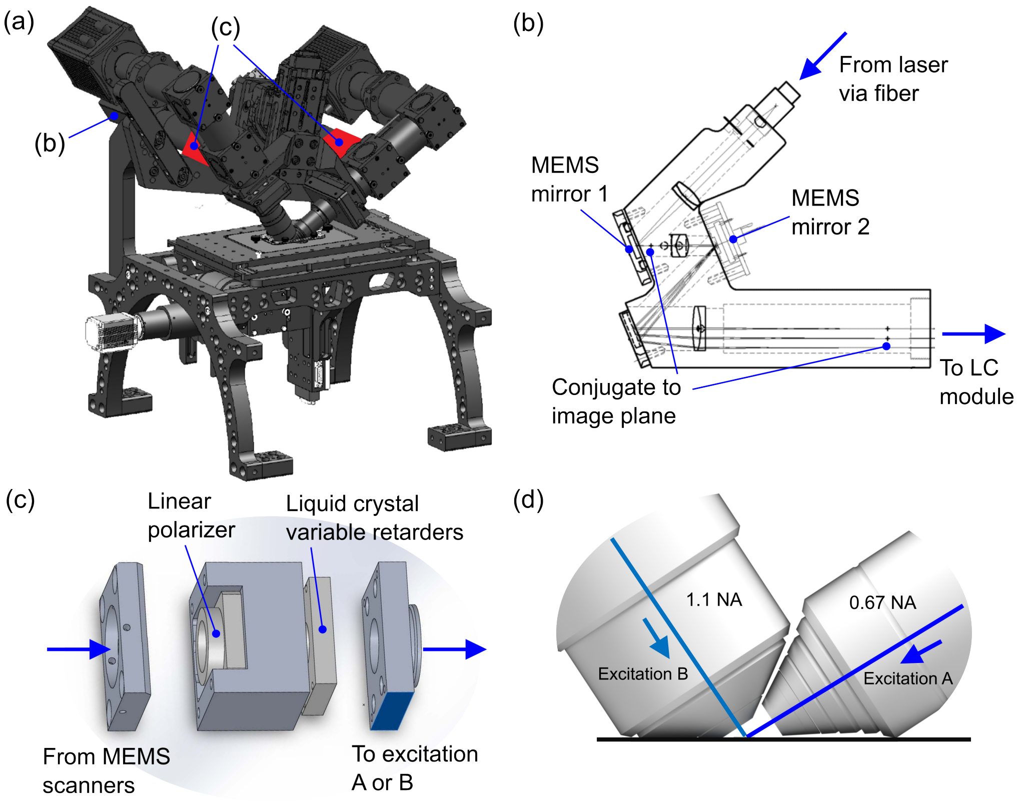


**Fig. S1 Key hardware and schematics for polarized dual-view inverted selective plane illumination microscope (pol-diSPIM).** (a) Overview of the microscope with the MEMS scanners detailed in (b) and the liquid crystal (LC) modules (red box in each arm) detailed in (c). (b) Schematic of the MEMS scanner module (Applied Scientific Instrumentation). The beam arrives from a fiber-coupled laser; reflects from MEMS mirror 1, positioned slightly offset from the position conjugate to the image plane for tilting the beam; reflects from MEMS mirror 2, positioned conjugate to the back focal plane of the objective for scanning the beam to create a light sheet; then continues to the tube lens and an LC module. (c) Exploded drawing of an LC module. The beam arrives from the MEMS scanner where it is linearly polarized before its polarization is modulated by a stacked pair of liquid-crystal variable retarders (LCVRs). The beam continues to excitation objective A or B where it forms a polarization- and tilt-controlled light sheet in the sample. (d) A view of the asymmetric objective pair (1.1 NA and 0.67 NA) with arrows indicating the excitation beams' propagation directions.

### 2 Degrees of freedom & finite sampling

The pol-diSPIM system has three degrees of freedom that allow it to acquire data that can be used to estimate the orientation of the molecules it images: its transverse illumination polarizations, its illumination tilt angles, and its views.

#### 2.1 Transverse illumination polarization

We modulate the illumination beam's transverse polarization with a liquid crystal (LC) module **Figure S1(c)**, which allows us to illuminate with arbitrary transverse polarizations. Although the LC module is capable of generating any transverse polarization state, we restrict our samples to six linear polarization states in each arm for the following reasons:

1. We restrict our illumination polarization states to **linear polarizations only**. The large majority of fluorescent emitters used in biological applications are excited via a linear dipole moment, so they are excited most efficiently by linearly polarized light. Illuminating the sample with elliptically polarized light would reduce contrast.
2. We restrict our linear polarization states to just **six states at 0°, 45°, 60°, 90°, 120°, and 135°** with respect to the $y$-axis (see **Figure S3(a, b)** for angle conventions). We know that dipoles are excited proportional to $\cos^2\theta$ where $\theta$ is the angle between the illumination polarization and the excitation dipole moment, so an arbitrary ensemble of dipoles is excited with a functional dependence of the form $a\cos^2(\theta-\phi)+b$, where $a$, $b$, and $\phi$ are unknowns. Three noise-free samples at different values of $\theta$ are sufficient to recover $a$, $b$, and $\phi$, but in practice we often operate in a noise-limited regime where some degree of oversampling is beneficial. Therefore, we chose to illuminate our sample with as many as six transverse polarization states.

#### 2.2 Illumination tilt angles

We modulate the tilt of the illumination beams with the MEMS mirror 1 in either arm. The MEMS mirrors allow us to tilt both illumination beams up to $\sim 10°$ in any direction, but we restrict our samples to three tilt angles in the plane of the light sheet, see **Figure S2(e, f)** for the orientation of the tilt, for the following reasons:

1. We restrict our tilt samples to just **three tilt angles** because only three samples are required to recover $a$, $b$, and $\phi$ from an $a\cos^2(\theta-\phi)+b$ signal. Unlike the 6-sample oversampling for the transverse polarization states, tilt angles beyond $\sim 10°$ are unavailable to the MEMS mirrors, so the benefits of oversampling are small.
2. We restricted our tilt samples to tilts **in the plane of the light sheet** so that the light sheet remains in the focal plane of the fixed perpendicular imaging arm.
3. We chose our tilt samples **at the nominal untilted angle (labelled 0), and at the maximum tilt angles in either direction (labelled ±1)** that did not create noticeably aberrated images. These tilt angles maximize the contrast available to the imaging system.

#### 2.3 Views

The pol-diSPIM employs an asymmetric two-arm geometry with a 1.1 NA objective and a 0.67 NA objective (**Figure S2**). Similar to conventional diSPIM [3, 4], one objective generates light-sheet illumination while the other objective collects the fluorescence (e.g., view A: 0.67 NA excitation, 1.1 NA detection, **Figure S2(a)**), and after an acquisition the roles of the two objectives are switched (e.g., view B: 1.1 NA excitation, 0.67 NA detection, **Figure S2(b)**).

While these two views are primarily for improving axial spatial resolution, they also provide additional orientation information. For example, a dipole oriented parallel to the 1.1 NA objective's optical axis will emit more light in the direction of the 0.67 NA view, so, all else equal, a larger signal in the 0.67 NA objective will lead us to estimate that a dipole is oriented closer to parallel with the optical axis of the 1.1 NA objective.

##### 2.4 Finite sampling and notation

We reduced a large set of possible samples down to a set of **36 possible configurations**—all combinations of 6 transverse polarizations $p \in \{0°, 45°, 60°, 90°, 120°, 135°\}$, 3 tilt angles $\mathtt{t} \in \{-1, 0, +1\}$, and 2 views $\mathtt{v} \in \{A, B\}$. See **Figure S3(c, d)** for an overview of all 36 configurations.

We can describe a single volumetric acquisition with a tuple $(p, \mathtt{t}, \mathtt{v})$ describing its polarization, tilt, and view. For example, $(45°, 0, A)$ refers to the volumetric sample acquired under polarized illumination 45° from the $\hat{\mathbf{y}}$-axis with no tilt from view $A$. In **Supplement 5** we will find it convenient to combine the $p$ and $\mathtt{t}$ indices into a single index $j$.

##### 2.5 Excitation sampling schemes

We proceeded to collect combinations of our 36 possible polarization-tilt-view configurations into what we call *excitation sampling schemes*, summarized in **Table S1**.

Our first scheme, named **All**, consists of a loop through all six polarizations including a repeat of the 0° polarization for bleaching estimation, all three tilts, and both views for a total of 42 samples. The **All** scheme oversamples fluorescent dipole emitters, but it is useful for calibrating with fixed samples like GUVs.

Our second scheme, named **Six no tilt**, consists of three transverse polarizations and both views for a total of six samples. The **Six no tilt** scheme undersamples the fluorescent dipole emitters, and it leaves some orientations unmeasured—see main text **Figure 3**.

Our final scheme, named **Six with tilt**, consists of six total samples, each acquired with a different transverse polarization, tilt, and view. The **Six with tilt** scheme is an optimized sampling scheme chosen to maximize the angular information we desire with a minimal number of samples. We performed an exhaustive search through all $\binom{36}{6} \approx 2 \times 10^6$ possible 6-sample schemes available to our instrument, and we chose the scheme that optimized the singular spectrum of the instrument—see **Supplement 8.3** for details.

#### 3 Data acquisition

To implement polarization imaging, we extended our original LabVIEW control software [4] to incorporate LC module control, beam tilting control, and additional functions that facilitated system alignment and calibration. Before imaging biological

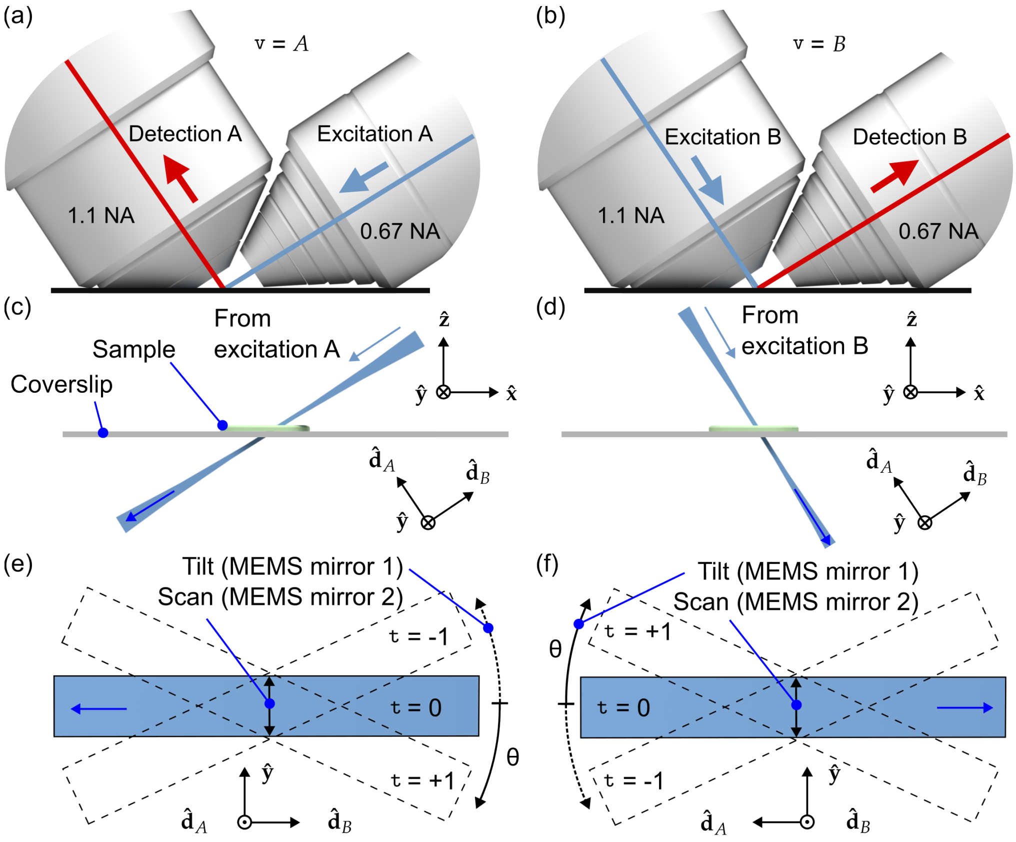


**Fig. S2 Schematic of light-sheet views, scanning, and tilting.** Columns: The first and second columns describe data acquired from views A and B, respectively. (a) View A illuminates the sample with a light-sheet from the 0.67 NA objective (blue arrow) and detects emitted light with the 1.1 NA objective (red arrow). (b) Similarly, view B illuminates with the 1.1 NA objective and detection with the 0.67 NA objective. (c) A view of a stationary Gaussian beam from excitation objective A illuminating a sample (green) on a coverslip. Here and throughout the figure, the blue arrow indicates the direction of light propagation. (d) Similarly, a view of a stationary Gaussian beam from excitation objective B. Subfigures (a)–(d) share a coordinate system with an $\hat{\mathbf{x}}$-$\hat{\mathbf{y}}$-$\hat{\mathbf{z}}$ coordinate system defined with respect to the coverslip and a $\hat{\mathbf{d}}_A$-$\hat{\mathbf{d}}_B$-$\hat{\mathbf{y}}$ coordinate system defined with respect to the objectives optical axis ($\hat{\mathbf{d}}_A$ and $\hat{\mathbf{d}}_B$ are shorthand for detection objective A and B, respectively). (e) A view of the illumination pattern A looking down the optical axis of detection objective A. The stationary Gaussian beam depicted in (c) and (d) can be scanned with MEMS mirror 2 to form a light sheet (blue rectangle), and tilted with MEMS mirror 1 to form tilted light sheets (dashed lines) labelled with their shorthand indices $\mathtt{t} \in \{-1, 0, +1\}$. Alternatively, the light-sheet tilt angles can be described by their tilt angle $\theta$ where positive/negative angles are measured from the $\hat{\mathbf{d}}_B$ axis to the negative/positive $\hat{\mathbf{y}}$ axis shown with a solid/dotted arrow. Note that the tilt angles are exaggerated for clarity; the real tilt angles are less than half the angles shown here. (f) Similarly, a view of illumination pattern B looking down the optical axis of detection objective B.

samples, the system was aligned (**Supplement 3.1**) and calibrated (**Supplement 3.2**).

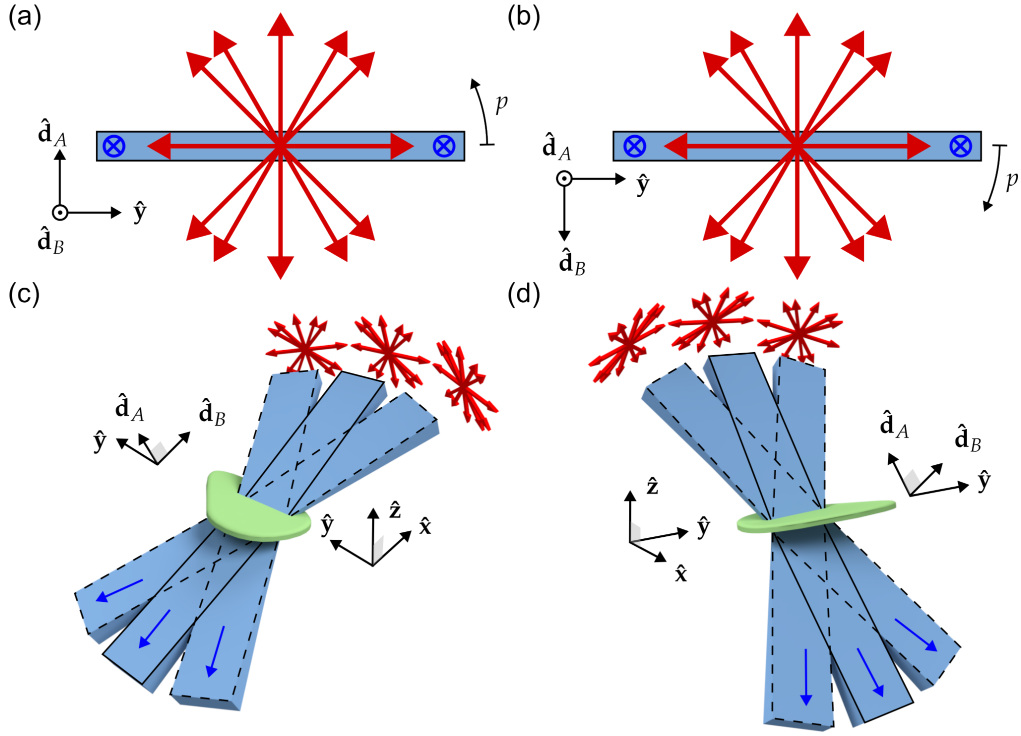


**Fig. S3 Schematic of light-sheet polarization settings.** Columns: The first and second columns describe data acquired from views A and B, respectively. (a) A view of a non-tilted illumination light sheet A (blue rectangle) looking down the optical axis of excitation objective A ($\hat{\mathbf{d}}_B$) with light propagating into the page (blue arrows). Red arrows indicate the linear transverse polarization states selected by the LCVR at orientations $p \in \{0°, 45°, 60°, 90°, 120°, 135°\}$, where $p$ is the angle from the $\hat{\mathbf{y}}$ axis in the direction of the $\hat{\mathbf{d}}_A$ axis. (b) Similarly, a view of a non-tilted illumination light sheet B looking down the optical axis of excitation objective B ($\hat{\mathbf{d}}_A$), where $p$ is the angle between the $\hat{\mathbf{y}}$ axis and the $\hat{\mathbf{d}}_B$ axis. (c) An overview of all 18 possible samples from view A—six polarization settings for each of three tilt settings. Notice that tilting the light sheet makes new polarization orientations accessible while illuminating the same positions in the sample. (d) Similarly, an overview of all 18 possible samples from view B.

#### 3.1 Alignment

To set up the imaging system, we perform an alignment routine to find the voltages to apply to the LC modules and the MEMS mirrors.

1. **Transverse polarization settings**: We use a wire-grid linear polarizer and a power meter to determine the LCVR voltages required to generate the polarization states, see **Figure S4**.
   (a) Mount the polarizer and power meter so that they are aligned with the optical axis of view A's illumination objective (the propagation direction of the beam from 0.67 NA objective when no tilt or scanning is applied), see **Figure S4(a)**.

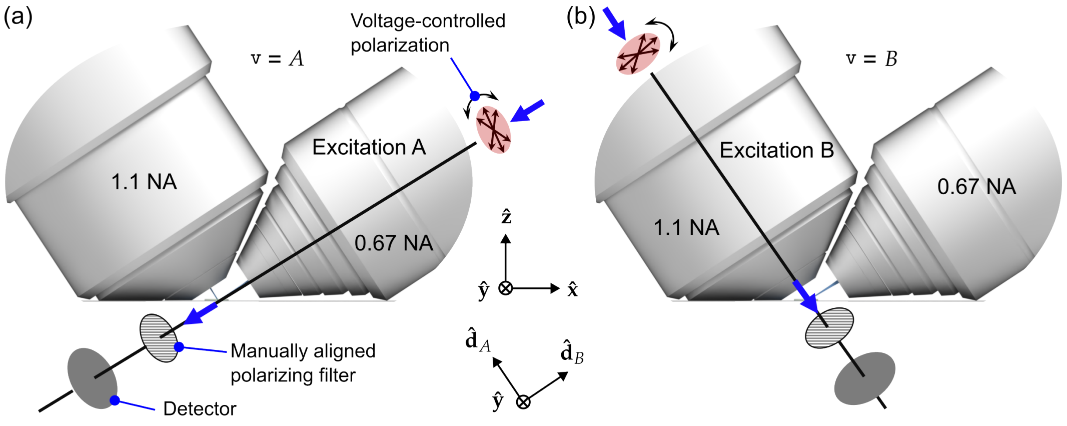


**Fig. S4 Procedure for finding the LCVR voltages that generate different illumination polarizations.** We mount a wire-grid linear polarizer and a power detector along the (a) view A illumination path, then (b) view B illumination path. To determine the LCVR voltages for generating the 6 polarization states, we orient the polarizer's transmission axis perpendicular to the desired illumination polarization orientation, then vary the LCVR voltages to extinguish the beam at the detector.

(b) For a specific polarization configuration, e.g., 0°, orient the polarizer's transmission axis perpendicular to the desired polarization orientation.

(c) With the laser on, vary the voltages on the two LCVRs (in the arm of view A illumination) until the readout of the detector is minimized (i.e find the voltage at which extinction is achieved, usually with a polarization extinction ratio >200:1), then record the voltages for this polarization configuration.

(d) Repeat steps (b) and (c) for all six polarization configurations and record the voltages.

(e) Unmount the polarizer and power meter from view A, and remount them to view B as shown in **Figure S4(b)**. Repeat steps (b), (c), and (d) to find the voltages for the 6 polarization configurations of view B.

2. **Beam tilt settings**: We choose and measure our illumination tilt angles by imaging a fluorescent bead solution (100 nm yellow-green beads, 1000-fold dilution) then:

(a) Starting with the view A illumination path, we illuminate with a non-scanned Gaussian beam and increase the tilt angle until the image quality noticeably degrades near the edge of the field of view. We record the MEMS mirror voltage, record an image of the stationary beam (see **Figure S5**) and measure its tilt angle from the image.

(b) Repeat step (a) for the opposite tilt and non-tilted conditions.

(c) Repeat steps (a) and (b) for view B.

#### 3.2 Calibration

Although our alignment routine gives us some confidence that the polarization orientations and tilts are where we expect them to be, in practice we find that day-to-day use and realignment of the microscope causes measurable drift of the illumination states.

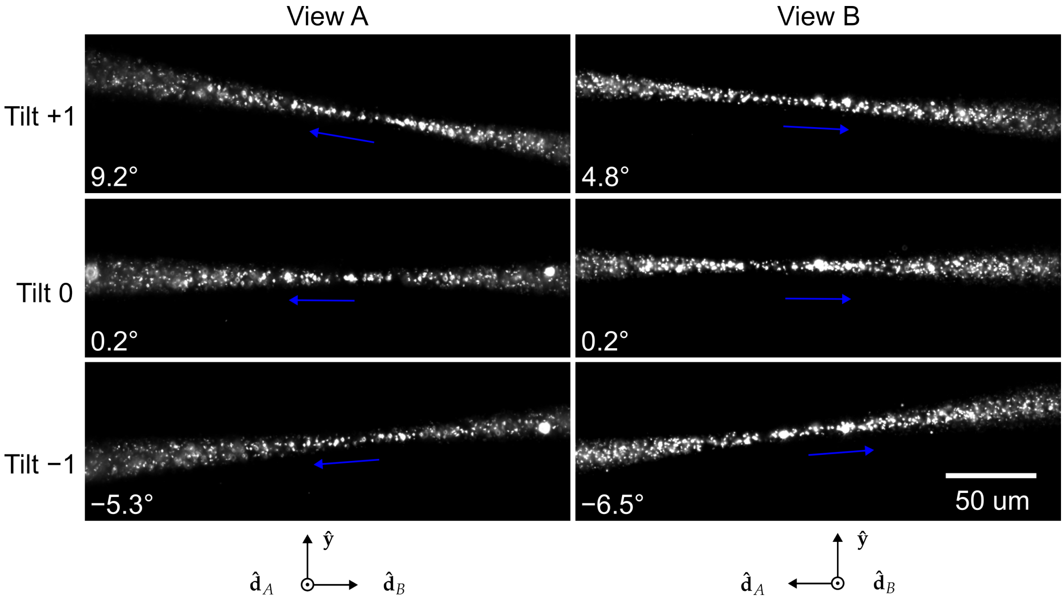


**Fig. S5** Images of fluorescent beads in solution under different illumination beam tilts (rows) and views (columns). Axis orientations for each view are shown at the bottom of each column, and the propagation directions are indicated with blue arrows. The orientation of the columns in this figure match **Figure S2(e, f)**.

.

In addition, we find empirically that the number of counts measured on each arm drifts independently. These variations led us to develop a calibration routine where we acquire data from a spatially and angularly uniform sample, an autofluorescent plastic slide (Chroma 92001). Immediately preceding each acquisition:

1. We mount an autofluorescent plastic slide in the microscope chamber.
2. We acquire three-dimensional volumes from within the fluorescent slide under all 36 illumination configurations. We record these data in a four-dimensional array $g^{(\text{cal})}_{(p,\mathtt{t},\mathtt{v})}(\mathbf{r}_d)$ where the tuple $(p, \mathtt{t}, \mathtt{v})$ indexes the 36 illumination configurations, and $\mathbf{r}_d$ is a three-dimensional detection coordinate.
3. We average the measured intensities over a volume $V$ from deep within the sample, and we record the 36 averaged calibration values in a vector $\bar{g}^{(\text{cal})}_{(p,\mathtt{t},\mathtt{v})} = \frac{1}{|V|}\sum_{\mathbf{r}_d \in V} g^{(\text{cal})}_{(p,\mathtt{t},\mathtt{v})}(\mathbf{r}_d)$.

We use the calibration values $\bar{g}^{(\text{cal})}_{(p,\mathtt{t},\mathtt{v})}$ to correct our raw data immediately before performing a reconstruction, see **Supplement 7.2** for details.

Note that we performed the alignment procedure approximately twice a year, while we performed the calibration procedure immediately before or after every acquisition session.

#### 3.3 Acquisition order

We acquired volumetric data by scanning the sample stage through a stationary light sheet, acquiring fluorescence images with 15–50 ms exposures for each image. Unless stated otherwise, the step size for all data was set to 1 $\mu$m per stage step (0.549 $\mu$m after deskewing in View A, 0.836 $\mu$m in View B). We acquired image volumes in both views before we changed the polarization/tilt state and acquired the next pair of volumes from both views. Our fastest volume acquisitions required 15 ms per slice for 40 slices for a total acquisition time of 0.6 s per volume. Our fastest complete acquisition required 0.6 s per volume for 6 volumes, making our fastest 3D-orientation-resolved time resolution 3.6 s.

For multicolor imaging, all views, polarizations, and tilts for one color were acquired, followed by all views, polarizations, and tilts for the next color until all colors were acquired.

In summary, we acquired datasets with as many as eight dimensions, and each dimension was collected in a loop in the following order from fastest to slowest: $(xy)$ camera frame, $(z)$ stage scan positions, $(\mathtt{v})$ views, $(p)$ polarization, $(\mathtt{t})$ tilts, $(c)$ colors, $(T)$ time points.

### 4 Preprocessing

#### 4.1 Deskewing

Since volumetric data were acquired by scanning the sample stage, raw images were deskewed, see Kumar et al. for details [5], then cropped to save storage and computational expense.

#### 4.2 Registration

After deskewing, the two view images were interpolated and upsampled to an isotropic voxel size of 130 nm $\times$ 130 nm $\times$ 130 nm. Then view B (0.67 NA objective illumination and 1.1 NA objective detection) images were rotated by $-90$ degrees about the $\hat{\mathbf{y}}$-axis so that they were coarsely aligned to View B (1.1 NA objective illumination and 0.67 NA objective detection) images. To estimate a registration transformation, we averaged the volumes acquired under different illumination conditions (polarizations, tilts) into a single volume for each view, then estimated the 12-dimensional affine transformation that maximized the cross correlation of the two volumes. Finally, we applied the estimated transformation to all of raw data volumes acquired in view B. All optimizations and transformations were performed with GPU-based 3D affine registration routines [6].

We found that averaging each view over illumination polarizations and tilts before estimating a registration transformation led to the most accurate registrations. When imaging a static sample with spatially varying oriented fluorescent dipoles, changing the illumination polarization will result in different regions of the sample appearing with different intensities. These shifts in intensity can be incorrectly identified as mis-registrations, which can lead to incorrect registration transformations. We reasoned

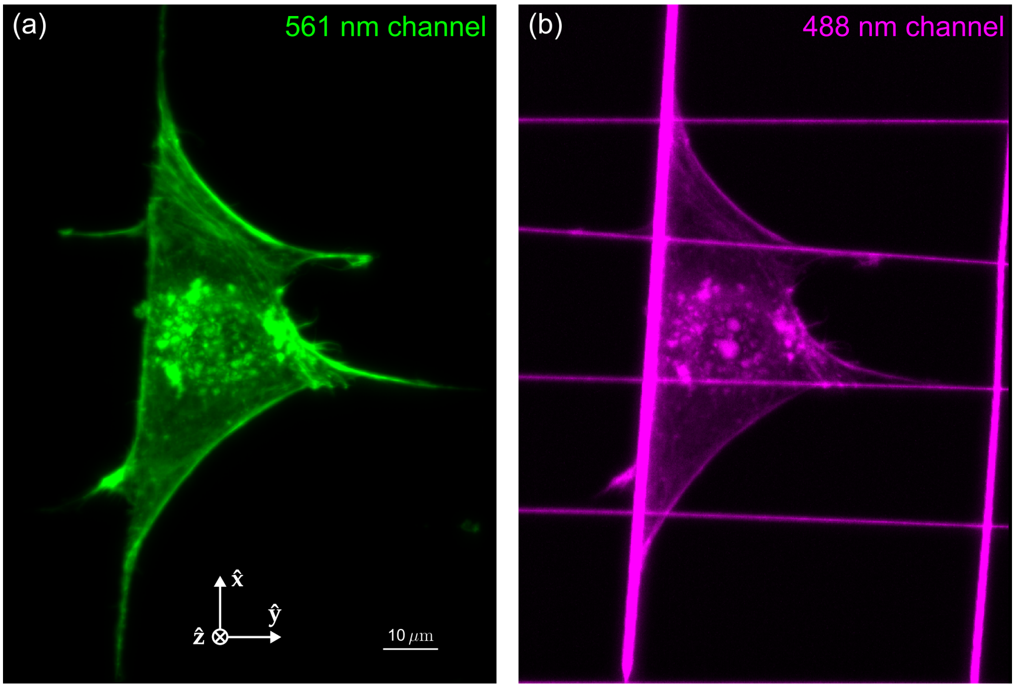


**Fig. S6 MIPs of raw nanowire data in two channels.** (a) 561 nm channel showing Alexa Fluor 568 phalloidin in one of six polarization channels. (b) 488 nm channel showing nanowires and crosstalk from the 561 nm channel. We hand annotated nanowires from the high-contrast 488 nm channel, then used the wire locations for further nanowire analysis.

that our registration algorithm should not be sensitive to any apparent shifts in intensity from a single view, so we averaged over illumination polarizations and tilts before estimating a registration transformation.

#### 4.3 Nanowire annotation

We manually annotated nanowires in 3D from a dedicated 488 nm channel. For example, **Figure S6(b)** shows the raw data we used to annotate nanowires for **main-text Figure 4**.

For each nanowire we loaded the 488 nm volume in 3D, viewed slices of the volumes with normals approximately parallel to the the long axes of the wire (e.g. $\hat{\mathbf{y}} - \hat{\mathbf{z}}$ slices for the wires along the $\hat{\mathbf{x}}$ axis, $\hat{\mathbf{x}} - \hat{\mathbf{z}}$ slices for the wires along the $\hat{\mathbf{y}}$ axis), then clicked the wire every $\sim 20\ \mu$m. We estimate that we were able to locate the centers of the nanowires within $\sim 2$ voxels $= 260$ nm.

### 5 Point-response functions

In the upcoming sections we describe our model of the imaging system. We begin with notation tables in **Tables S2 and S3** before starting with our point-response function calculations.

#### 5.1 Description of the data

Our imaging system illuminates the object with a sheet of light, collects a two-dimensional irradiance image, then scans the object through the stationary light sheet to build a three-dimensional dataset that we can index with a three-dimensional coordinate $\mathfrak{r}_d \in \mathbb{R}^3$. We repeat these three-dimensional acquisitions for different combinations of transverse polarizations $p$, tilts $\mathtt{t}$, and views $\mathtt{v}$—that we write together in a tuple $(p, \mathtt{t}, \mathtt{v})$. This means that we can describe a complete set of irradiance measurements with $g_{(p,\mathtt{t},\mathtt{v})}(\mathfrak{r}_d) \in \mathbb{L}_2(\mathbb{R}^3)^N$, where $N$ is the total number of volume acquisitions.

All of our datasets consist of an equal number of transverse-polarization and tilt settings for each view, so it is convenient to combine the transverse-polarization index $p$ and the tilt index $\mathtt{t}$ into a single polarization index $j$. This allows us to rewrite our complete dataset as $g_{j\mathtt{v}}(\mathfrak{r}_d) \in \mathbb{L}_2(\mathbb{R}^3)^N$—see **Supplement 2.4**.

#### 5.2 General relationship between the object and the data

We start by creating a reasonably general description of fluorescent dipoles in a sample. First, consider an ensemble of two-state fluorescent molecules with aligned dipole absorption and emission moments. These molecules undergo spatial and angular diffusion while being excited with illumination light and decaying to emit photons that we can manipulate and detect. We make multiple measurements of the object by manipulating the excitation light or by manipulating the emitted photons onto our detectors, but we can not control the position, diffusion, or decay dynamics of the molecules. Our goal is to describe the relationship between the object and the data we measure, so that we can recover as much as possible about the position, orientation, and dynamics of these fluorescent molecules. Ideally enough information can be recovered that the experimentalist can draw new conclusions about the molecules and their environment.

We can describe this ensemble of molecules using two functions $f^{(\text{gr})}(\mathfrak{r}_o, \hat{\mathbf{s}}_o, t)$ and $f^{(\text{ex})}(\mathfrak{r}_o, \hat{\mathbf{s}}_o, t)$ that describe the number of molecules per unit volume at position $\mathfrak{r}_o \in \mathbb{R}^3$ per unit solid angle oriented along $\hat{\mathbf{s}}_o \in \mathbb{S}^2$ per unit time at time $t$ in the ground and excited states, respectively. We assume that these molecules are diffusing in a spatio-angular potential $v(\mathfrak{r}_o, \hat{\mathbf{s}}_o)$, that they are decaying with a rate constant $\kappa^{(\text{d})}(\mathfrak{r}_o, \hat{\mathbf{s}}_o)$, and that during measurement $g_{j\mathtt{v}}(\mathfrak{r}_d)$ they are being excited with a rate constant $h^{(\text{ex})}_{j\mathtt{v}}(\mathfrak{r}_d, \mathfrak{r}_o, \hat{\mathbf{s}}_o)$. These dynamics can be captured by the following pair of coupled differential equations

$$\frac{\partial}{\partial t}\begin{bmatrix} f^{(\text{ex})}(\mathfrak{r}_o, \hat{\mathbf{s}}_o, t) \\ f^{(\text{gr})}(\mathfrak{r}_o, \hat{\mathbf{s}}_o, t) \end{bmatrix} = \begin{bmatrix} \mathscr{D}_{\boldsymbol{v}} - \kappa^{(\text{d})}(\mathfrak{r}_o, \hat{\mathbf{s}}_o) & h^{(\text{ex})}_{j\mathtt{v}}(\mathfrak{r}_d, \mathfrak{r}_o, \hat{\mathbf{s}}_o) \\ \kappa^{(\text{d})}(\mathfrak{r}_o, \hat{\mathbf{s}}_o) & \mathscr{D}_{\boldsymbol{v}} - h^{(\text{ex})}_{j\mathtt{v}}(\mathfrak{r}_d, \mathfrak{r}_o, \hat{\mathbf{s}}_o) \end{bmatrix} \begin{bmatrix} f^{(\text{ex})}(\mathfrak{r}_o, \hat{\mathbf{s}}_o, t) \\ f^{(\text{gr})}(\mathfrak{r}_o, \hat{\mathbf{s}}_o, t) \end{bmatrix}, \tag{S1}$$

where $\mathscr{D}_{\boldsymbol{v}}$ is a Smoluchowski operator that models spatio-angular diffusion

$$\mathscr{D}_{\boldsymbol{v}} = \nabla \cdot \exp[-\beta v(\mathbf{r}_o, \hat{\mathbf{s}}_o)]\mathbf{D}\exp[\beta v(\mathbf{r}'_o, \hat{\mathbf{s}}'_o)], \tag{S2}$$

where $\nabla$ is a gradient on $\mathbb{R}^3 \times \mathbb{S}^2$, $\mathbf{D}$ is a generalized diffusion tensor, and $\beta = 1/k_BT$.

If the molecules start in the ground state with spatial density $\rho(\mathbf{r}_o)$, then the initial condition is given by $f^{(\text{ex})}(\mathbf{r}_o, \hat{\mathbf{s}}_o, 0) = 0$ and $f^{(\text{gr})}(\mathbf{r}_o, \hat{\mathbf{s}}_o, 0) = \rho(\mathbf{r}_o)$. After solving **Equation S1** we can model the emission density during measurement $g_{j\mathtt{v}}(\mathbf{r}_d)$ from time $t_{j\mathtt{v}}^{(\text{start})}(\mathbf{r}_d)$ to $t_{j\mathtt{v}}^{(\text{end})}(\mathbf{r}_d)$ as

$$f_{j\mathtt{v}}^{(\text{em})}(\mathbf{r}_d, \mathbf{r}_o, \hat{\mathbf{s}}_o) = \int_{t_{j\mathtt{v}}^{(\text{start})}(\mathbf{r}_d)}^{t_{j\mathtt{v}}^{(\text{end})}(\mathbf{r}_d)} dt\, \kappa^{(\text{d})}(\mathbf{r}_o, \hat{\mathbf{s}}_o) f^{(\text{ex})}(\mathbf{r}_o, \hat{\mathbf{s}}_o, t). \tag{S3}$$

Finally, we can relate the irradiance measurements $g_{j\mathtt{v}}(\mathbf{r}_d)$ to object properties by integrating the emission density $f_{j\mathtt{v}}^{(\text{em})}(\mathbf{r}_d, \mathbf{r}_o, \hat{\mathbf{s}}_o)$ weighted by the point-response function of the imaging system $h_{j\mathtt{v}}^{(\text{det})}(\mathbf{r}_d, \mathbf{r}_o, \hat{\mathbf{s}}_o)$

$$g_{j\mathtt{v}}(\mathbf{r}_d) = \int_{\mathbb{R}^3} d\mathbf{r}_o \int_{\mathbb{S}^2} d\hat{\mathbf{s}}_o\, h_{j\mathtt{v}}^{(\text{det})}(\mathbf{r}_d, \mathbf{r}_o, \hat{\mathbf{s}}_o) f_{j\mathtt{v}}^{(\text{em})}(\mathbf{r}_d, \mathbf{r}_o, \hat{\mathbf{s}}_o). \tag{S4}$$

**Equations S3 and S4** describe a non-linear relationship between an object property, the excited state population $f^{(\text{ex})}(\mathbf{r}_o, \hat{\mathbf{s}}_o, t)$, and the measured data, irradiance measurements $g_{j\mathtt{v}}(\mathbf{r}_d)$. Without additional assumptions about the sample, this non-linearity will make it difficult to recover useful information about the sample.

#### 5.3 Linear relationship between the object and the data

One way to find a linear relationship between object properties and the data is to make assumptions about our object and measurements and arrange experimental conditions that meet those assumptions. In [7] we showed that if (a) spatial diffusion is negligible, (b) angular diffusion is consistent with a spherical rotor model, (c) excitation is weak to avoid saturation effects, (d) angular diffusion times are long compared to the fluorescence decay times, (e) exposure times are long compared to the diffusion and fluorescence decay times, (f) and the measurements are collected long after initial transient diffusion and fluorescence decay times (fluorescence dynamics have reached a steady state), then the following relationship holds

$$g_{j\mathtt{v}}(\mathbf{r}_d) = \int_{\mathbb{R}^3} d\mathbf{r}_o \int_{\mathbb{S}^2} d\hat{\mathbf{s}}_o\, h_{j\mathtt{v}}(\mathbf{r}_d, \mathbf{r}_o, \hat{\mathbf{s}}_o) f(\mathbf{r}_o, \hat{\mathbf{s}}_o), \tag{S5}$$

where

$$h_{j\mathtt{v}}(\mathbf{r}_d, \mathbf{r}_o, \hat{\mathbf{s}}_o) = h_{j\mathtt{v}}^{(\text{det})}(\mathbf{r}_d, \mathbf{r}_o, \hat{\mathbf{s}}_o) h_{j\mathtt{v}}^{(\text{exc})}(\mathbf{r}_d, \mathbf{r}_o, \hat{\mathbf{s}}_o) \tag{S6}$$

is the *spatio-angular point response function*, and

$$f(\mathbf{r}_o, \hat{\mathbf{s}}_o) = \rho(\mathbf{r}_o)\frac{\exp[-\beta v(\mathbf{r}_o, \hat{\mathbf{s}}_o)]}{\int_{\mathbb{S}^2} d\hat{\mathbf{s}}\ \exp[-\beta v(\mathbf{r}_o, \hat{\mathbf{s}})]} \tag{S7}$$

is the *spatio-angular Boltzmann density*—the product of the labeling density $\rho(\mathbf{r}_o)$ and the angular Boltzmann distribution at each spatial point. The relationship between the spatio-angular Boltzmann density $f(\mathbf{r}_o, \hat{\mathbf{s}}_o)$ and the data $g_{j\texttt{v}}(\mathbf{r}_d)$ is linear, so the spatio-angular Boltzmann density is a good candidate for us to reconstruct. Therefore, **Equation S7** relates the sample properties $\rho(\mathbf{r}_o)$ (the spatial labelling density) and $v(\mathbf{r}_o, \hat{\mathbf{s}}_o)$ (the spatio-angular potential) to our reconstruction target $f(\mathbf{r}_o, \hat{\mathbf{s}}_o)$.

By writing the sample properties $\rho(\mathbf{r}_o)$ and $v(\mathbf{r}_o, \hat{\mathbf{s}}_o)$ without any time dependence, we are making an additional assumption (g) that these sample properties do not change appreciably over the course of a complete set of measurements $g_{j\texttt{v}}(\mathbf{r}_d)$. When we measure living samples that move on timescales faster than it takes us to acquire a complete set of measurements, $\sim$ 3.6 s for our fastest acquisitions, this assumption is no longer true, and we are in danger of misinterpreting sample motion as intensity modulations that indicate an oriented sample.

We emphasize that the spatio-angular Boltzmann density is only a reasonable target for linear reconstruction when conditions (a)–(g) are satisfied, and throughout this paper we assume that conditions (a)–(g) are satisfied.

We now proceed to give explicit expressions for the excitation point-response function $h_{j\texttt{v}}^{(\text{exc})}(\mathbf{r}_d, \mathbf{r}_o, \hat{\mathbf{s}}_o)$ and the dectection point-response function $h_{\texttt{v}}^{(\text{det})}(\mathbf{r}_d, \mathbf{r}_o, \hat{\mathbf{s}}_o)$.

#### 5.4 Excitation point-response function

The excitation point-response function $h_{j\texttt{v}}^{(\text{exc})}(\mathbf{r}_d, \mathbf{r}_o, \hat{\mathbf{s}}_o)$ can be interpreted as the probability of exciting a dipole emitter at position $\mathbf{r}_o$ oriented along $\hat{\mathbf{s}}_o$ when the demagnified detector coordinate is in position $\mathbf{r}_d$ and the sample is illuminated with polarization $j$ and imaged with view $\texttt{v}$.

We create all of our illumination light sheets by scanning paraxial Gaussian beams, so we assume that our illumination polarization does not vary spatially across the illumination pattern. This allows us to factor the excitation point-response function into two functions

$$h_{j\texttt{v}}^{(\text{exc})}(\mathbf{r}_d, \mathbf{r}_o, \hat{\mathbf{s}}_o) = h_{\texttt{v}}^{(\text{exc, sp})}(\mathbf{r}_d, \mathbf{r}_o)h_{j\texttt{v}}^{(\text{exc, ang})}(\hat{\mathbf{s}}_o), \tag{S8}$$

where $h_{\texttt{v}}^{(\text{exc, sp})}(\mathbf{r}_d, \mathbf{r}_o)$ is the *spatial* excitation point-response function and $h_{j\texttt{v}}^{(\text{exc, ang})}(\hat{\mathbf{s}}_o)$ is the *angular* excitation point-response function.

The form of the spatial excitation point-response function is simple to write for each view individually. For view $A$, we illuminate the sample with a focused Gaussian beam propagating along the negative $\hat{\mathbf{d}}_B$ axis and scan the beam along the $\hat{\mathbf{y}}$ axis to create a light sheet in the $\hat{\mathbf{d}}_B$-$\hat{\mathbf{y}}$ plane, see **Figure S2(c, e)**, and we can model this with

$$h_A^{(\text{exc, sp})}(\mathbf{r}_d, \mathbf{r}_o) = \left[\frac{w_0}{w(\mathbf{r}_d \cdot \hat{\mathbf{d}}_B)}\right]^2 \exp\left[\frac{-2\left((\mathbf{r}_d - \mathbf{r}_o)\cdot \hat{\mathbf{d}}_A\right)^2}{w(\mathbf{r}_d \cdot \hat{\mathbf{d}}_B)^2}\right], \tag{S9}$$

where $w_0$ is the beam waist radius, $w(x) = w_0\sqrt{1+(x/x_R)^2}$ is the depth-dependent beam radius, and $x_R = \pi w_0^2 n_0/\lambda$ is the Rayleigh range. Similarly, the spatial excitation point-response function for view $B$ can be written by swapping $\hat{\mathbf{d}}_A$ and $\hat{\mathbf{d}}_B$ in **Equation S9**:

$$h_B^{(\text{exc, sp})}(\mathbf{r}_d, \mathbf{r}_o) = \left[\frac{w_0}{w(\mathbf{r}_d \cdot \hat{\mathbf{d}}_A)}\right]^2 \exp\left[\frac{-2\left((\mathbf{r}_d - \mathbf{r}_o)\cdot \hat{\mathbf{d}}_B\right)^2}{w(\mathbf{r}_d \cdot \hat{\mathbf{d}}_A)^2}\right]. \tag{S10}$$

We can combine **Equations S9 and S10** into a single equation by defining a pair of rotation matrices

$$\mathbf{R}_A = \begin{bmatrix} 1 & 0 & 0 \\ 0 & 1 & 0 \\ 0 & 0 & 1 \end{bmatrix}, \qquad \mathbf{R}_B = \begin{bmatrix} 0 & 0 & 1 \\ 0 & -1 & 0 \\ 1 & 0 & 0 \end{bmatrix}, \tag{S11}$$

and writing the complete spatial excitation point-response function as

$$h_{\text{v}}^{(\text{exc, sp})}(\mathbf{r}_d, \mathbf{r}_o, \hat{\mathbf{s}}_o) = \left[\frac{w_0}{w\left(\mathbf{R}_{\text{v}}^{-1}\mathbf{r}_d \cdot \hat{\mathbf{d}}_B\right)}\right]^2 \exp\left[\frac{-2\left(\mathbf{R}_{\text{v}}^{-1}(\mathbf{r}_d - \mathbf{r}_o)\cdot \hat{\mathbf{d}}_A\right)^2}{w\left(\mathbf{R}_{\text{v}}^{-1}\mathbf{r}_d \cdot \hat{\mathbf{d}}_B\right)^2}\right]. \tag{S12}$$

Finally, the normalized angular excitation point-response function is given by

$$h_{j\text{v}}^{(\text{exc,ang})}(\hat{\mathbf{s}}_o) = \frac{3}{\sqrt{4\pi}}(\hat{\mathbf{p}}_{j\text{v}} \cdot \hat{\mathbf{s}}_o)^2, \tag{S13}$$

where $\hat{\mathbf{p}}_{j\text{v}}$ is $j\text{v}$-th polarization axis, which models the $\cos^2\theta$-dependence of excitation where $\theta$ is the angle between the polarization axis $\hat{\mathbf{p}}_{j\text{v}}$ and the dipole axis $\hat{\mathbf{s}}_o$.

#### 5.5 Three-dimensional shift invariance from uniform-thickness illumination

We have assembled a complete excitation point-response function in **Equations S8, S12, and S13**, but this model is expensive to compute. Fortunately, we can make a reasonable approximation and find a three-dimensionally shift-invariant model that captures the most important features of the illumination.

We assume that the illumination light sheet does not broaden appreciably across our imaging field of view so that $w(z) \approx w_0$. Under this assumption the model in

**Equation S8** becomes shift-invariant, so we can make the substitution $\mathfrak{r} = \mathfrak{r}_d - \mathfrak{r}_o$ and write a simplified spatial excitation point-response function

$$h_{j\mathtt{v}}^{(\text{exc, sp})}(\mathfrak{r}, \hat{\mathbf{s}}_o) \overset{(\text{unif})}{=} \exp\Big[-2\big(r_{\mathtt{v}}^{\parallel}/w_0\big)^2\Big], \tag{S14}$$

where $r_{\mathtt{v}}^{\parallel}$ is a view-dependent axial coordinate given explicitly by

$$r_A^{\parallel} = \mathfrak{r} \cdot \hat{\mathbf{d}}_A, \tag{S15}$$

$$r_B^{\parallel} = \mathfrak{r} \cdot \hat{\mathbf{d}}_B. \tag{S16}$$

This uniform-sheet assumption is valid within approximately one Rayleigh range of the beam's focus.

#### 5.6 Detection point-response function

The detection point-response function $h_{j\mathtt{v}}^{(\text{det})}(\mathfrak{r}_d, \mathfrak{r}_o, \hat{\mathbf{s}}_o)$ can be interpreted as the probability of detecting a photon from a dipole emitter at position $\mathfrak{r}_o$ oriented along $\hat{\mathbf{s}}_o$ when the detector is in demagnified position coordinate $\mathfrak{r}_d$ and the sample is illuminated with polarization $j$ and imaged with view $\mathtt{v}$.

Our detection point-response function is three-dimensionally shift invariant, i.e. shifting $\mathfrak{r}_o$ and $\mathfrak{r}_d$ together will leave the function unchanged, so we can safely replace $\mathfrak{r}_d$ and $\mathfrak{r}_o$ with $\mathfrak{r} = \mathfrak{r}_d - \mathfrak{r}_o$. Also, our measurements do not change when we modify the illumination polarization or tilt, so we can safely drop the $j$ dependence. Together, these notational reductions allow us to seek a simplified form $h_{\mathtt{v}}^{(\text{det})}(\mathfrak{r}, \hat{\mathbf{s}}_o)$.

To model the detection point-response function, $h_{\mathtt{v}}^{(\text{det})}(\mathfrak{r}, \hat{\mathbf{s}}_o)$, we start by rewriting it in view-dependent coordinates so that we can reuse results from the literature. If we let $h^{(\text{det},4f)}(\mathbf{r}^{\perp}, r^{\parallel}, \hat{\mathbf{s}}, \text{NA})$ denote the irradiance created at a demagnified off-axis point $\mathbf{r}^{\perp}$ on a two-dimensional detector behind an aplanatic $4f$ optical system with a paraxial tube lens and a detection objective with numerical aperture NA by an on-axis dipole defocused by $r^{\parallel}$ with orientation $\hat{\mathbf{s}}$, then the detection point-response function $h_{\mathtt{v}}^{(\text{det})}(\mathfrak{r}, \hat{\mathbf{s}}_o)$ can be written in terms of $h^{(\text{det},4f)}(\mathbf{r}^{\perp}, r^{\parallel}, \hat{\mathbf{s}}, \text{NA})$ as

$$h_{\mathtt{v}}^{(\text{det})}(\mathfrak{r}, \hat{\mathbf{s}}_o) = h^{(\text{det},4f)}\big(\mathbf{r}_{\mathtt{v}}^{\perp}, r_{\mathtt{v}}^{\parallel}, \hat{\mathbf{s}}_{\mathtt{v}}, \text{NA}_{\mathtt{v}}\big)\,, \tag{S17}$$

where the first argument

$$\mathbf{r}_{\mathtt{v}}^{\perp} = \mathbf{R}_{\mathtt{v}}^{-1}\mathfrak{r} - \big[\mathbf{R}_{\mathtt{v}}^{-1}\mathfrak{r} \cdot \hat{\mathbf{d}}_A\big]\hat{\mathbf{d}}_A, \tag{S18}$$

is a view-dependent transverse coordinate, the second argument

$$r_{\mathtt{v}}^{\parallel} = \mathbf{R}_{\mathtt{v}}^{-1}\mathfrak{r} \cdot \hat{\mathbf{d}}_A \tag{S19}$$

is a view-dependent axial coordinate, the third argument

$$\hat{\mathbf{s}}_{\mathtt{v}} = \mathbf{R}_{\mathtt{v}}^{-1}\hat{\mathbf{s}}_o \tag{S20}$$

is a view-dependent angular coordinate, and the last argument is the view-dependent detection numerical aperture

$$\text{NA}_A = 1.1, \tag{S21}$$
$$\text{NA}_B = 0.67. \tag{S22}$$

The detection point spread function for an aplanatic $4f$ optical system with a paraxial tube lens written in demagnified detection coordinates is given by [8–11]

$$h^{(\text{det},4f)}(\mathbf{r}^{\perp}, r^{\parallel}, \hat{\mathbf{s}}, \text{NA}) = \sum_{n,n'=0,1,2} b_{nn'}(\mathbf{r}^{\perp}, r^{\parallel}, \text{NA})\, s_n s_{n'}, \tag{S23}$$

where

$$b_{nn'}(\mathbf{r}^{\perp}, r^{\parallel}, \text{NA}) = \sum_{i=0,1} \beta_{in}(\mathbf{r}^{\perp}, r^{\parallel}, \text{NA})\beta^*_{in'}(\mathbf{r}^{\perp}, r^{\parallel}, \text{NA}) \tag{S24}$$

is the irradiance created at position $\mathbf{r}^{\perp}$ on the detector when an $s_n s_{n'}$ angular distribution is placed at defocus position $r_o^{\parallel}$,

$$\beta_{in}(\mathbf{r}^{\perp}, r^{\parallel}, \text{NA}) = \int_{\mathbb{R}^2} d\boldsymbol{\tau}\, A(\boldsymbol{\tau}, \text{NA})\Phi(\boldsymbol{\tau}, r^{\parallel})\gamma_{in}(\boldsymbol{\tau}) \exp[i2\pi\boldsymbol{\tau}\cdot\mathbf{r}^{\perp}] \tag{S25}$$

is the $i$th component of the electric field created at position $\mathbf{r}^{\perp}$ on the detector by the $n$th component of a dipole,

$$A(\boldsymbol{\tau}, \text{NA}) = (1 - (|\boldsymbol{\tau}|/\nu_m)^2)^{-1/4}\Pi\left(|\boldsymbol{\tau}|/\nu_c(\text{NA})\right) \tag{S26}$$

is the aplanatic apodization function with full width $\nu_c(\text{NA}) = 2\text{NA}/\lambda$ and $\nu_m = n_0/\lambda$,

$$\Phi(\boldsymbol{\tau}, r^{\parallel}) = \exp\left[i2\pi r^{\parallel}\sqrt{\nu_m^2 - |\boldsymbol{\tau}|^2}\right] \tag{S27}$$

encodes the defocus phase, the functions $\gamma_{in}(\boldsymbol{\tau})$ model the $i$th field components in the pupil plane created by the $n$th component of a dipole where $|\boldsymbol{\tau}|$ and $\phi_\tau$ are polar coordinates in the pupil plane

$$\begin{aligned}
\gamma_{00}(\boldsymbol{\tau}) &= \sin^2\phi_\tau + \cos^2\phi_\tau\sqrt{1-(|\boldsymbol{\tau}|/\nu_m)^2}, & \gamma_{10}(\boldsymbol{\tau}) &= \frac{1}{2}\sin(2\phi_\tau)\left(\sqrt{1-(|\boldsymbol{\tau}|/\nu_m)^2}-1\right),\\
\gamma_{01}(\boldsymbol{\tau}) &= \frac{1}{2}\sin(2\phi_\tau)\left(\sqrt{1-(|\boldsymbol{\tau}|/\nu_m)^2}-1\right), & \gamma_{11}(\boldsymbol{\tau}) &= \cos^2\phi_\tau + \sin^2\phi_\tau\sqrt{1-(|\boldsymbol{\tau}|/\nu_m)^2},\\
\gamma_{02}(\boldsymbol{\tau}) &= |\boldsymbol{\tau}|\cos\phi_\tau, & \gamma_{12}(\boldsymbol{\tau}) &= |\boldsymbol{\tau}|\sin\phi_\tau,
\end{aligned} \tag{S28}$$

and $s_n$ is the $n$th component of the dipole orientation coordinate $\hat{\mathbf{s}}_o$.

#### 5.7 Gaussian axial response

We can write the complete point-response function under the uniform-sheet approximation as

$$h_{j\mathrm{v}}(\mathbf{r}, \hat{\mathbf{s}}_o) \overset{\text{(unif)}}{=} \exp\left[-2\left(r_\mathrm{v}^\parallel/w_0\right)^2\right] h_{j\mathrm{v}}^{\text{(exc, ang)}}(\hat{\mathbf{s}}_o)\, h^{(\text{det},4f)}\left(\mathbf{r}_\mathrm{v}^\perp, r_\mathrm{v}^\parallel, \hat{\mathbf{s}}_\mathrm{v}, \mathrm{NA}_\mathrm{v}\right). \tag{S29}$$

Next, we assume that the axial dependence of the $4f$ detection point-response function is approximately Gaussian over the width of the excitation light sheet. That is

$$\exp\left[-2\left(r_\mathrm{v}^\parallel/w_0\right)^2\right] h^{(\text{det},4f)}\left(\mathbf{r}_\mathrm{v}^\perp, r_\mathrm{v}^\parallel, \hat{\mathbf{s}}_\mathrm{v}, \mathrm{NA}_\mathrm{v}\right) \tag{S30}$$

$$\approx \exp\left[-2\left(r_\mathrm{v}^\parallel/w_*\right)^2\right] h^{(\text{det},4f)}\left(\mathbf{r}_\mathrm{v}^\perp, 0, \hat{\mathbf{s}}_\mathrm{v}, \mathrm{NA}_\mathrm{v}\right), \tag{S31}$$

where **Equation S30** is the unapproximated result, and **Equation S31** is the approximate result with the $4f$-detection point-response function evaluated at $r_\mathrm{v}^\parallel = 0$ and a new axial width $w_* > w_0$. Applying **Equation S31** to **Equation S29** yields

$$h_{j\mathrm{v}}(\mathbf{r}, \hat{\mathbf{s}}_o) \underset{\text{(Gauss)}}{\overset{\text{(unif)}}{=}} \exp\left[-2\left(r_\mathrm{v}^\parallel/w_*\right)^2\right] h_{j\mathrm{v}}^{\text{(exc, ang)}}(\hat{\mathbf{s}}_o)\, h^{(\text{det},4f)}\left(\mathbf{r}_\mathrm{v}^\perp, 0, \hat{\mathbf{s}}_\mathrm{v}, \mathrm{NA}_\mathrm{v}\right). \tag{S32}$$

We can restate this assumption by claiming that the point-response function of the entire imaging system (both excitation and detection together) is approximately axially Gaussian for both viewing axes. Empirically, we find this to be true of our light sheets.

This assumption is also pragmatic—it is difficult to model and/or estimate the direct excitation width of the light sheet $w_o$, but it is straightforward to observe the total width of the light sheet $w_*$.

#### 5.8 Summary of forward model and point-response function

We complete this section by summarizing our model of the imaging system. First, our high-level model of the imaging system is

$$g_{j\mathrm{v}}(\mathbf{r}_d) = \int_{\mathbb{R}^3} d\mathbf{r}_o \int_{\mathbb{S}^2} d\hat{\mathbf{s}}_o\, h_{j\mathrm{v}}(\mathbf{r}_d - \mathbf{r}_o, \hat{\mathbf{s}}_o) f(\mathbf{r}_o, \hat{\mathbf{s}}_o), \tag{S33}$$

where $g_{j\mathrm{v}}(\mathbf{r}_d)$ is the irradiance measured on the detector at position $\mathbf{r}_d$ under view $\mathrm{v}$ and illumination polarization $j$, $f(\mathbf{r}_o, \hat{\mathbf{s}}_o)$ is the Boltzmann density at position $\mathbf{r}_o$ and orientation $\hat{\mathbf{s}}_o$, and $h_{j\mathrm{v}}(\mathbf{r}_d - \mathbf{r}_o, \hat{\mathbf{s}}_o)$ is the point-response function under the uniform-thickness and axial-Gaussian approximations given explicitly by

$$h_{j\mathrm{v}}(\mathbf{r}, \hat{\mathbf{s}}_o) = \exp\left[-2\left(r_\mathrm{v}^\parallel/w_*\right)^2\right] h_{j\mathrm{v}}^{\text{(exc, ang)}}(\hat{\mathbf{s}}_o)\, h^{(\text{det},4f)}\left(\mathbf{r}_\mathrm{v}^\perp, 0, \hat{\mathbf{s}}_\mathrm{v}, \mathrm{NA}_\mathrm{v}\right), \tag{S34}$$

where $h_{j\mathrm{v}}^{\text{(exc, ang)}}(\hat{\mathbf{s}}_o)$ is given by **Equation S12** and $h^{(\text{det},4f)}\left(\mathbf{r}_\mathrm{v}^\perp, 0, \hat{\mathbf{s}}_\mathrm{v}, \mathrm{NA}_\mathrm{v}\right)$ is given by **Equation S17**.

Notice that the uniform-thickness and axial-Gaussian point-response function is three-dimensionally shift invariant, so we can write it in terms of $\mathbf{r}_d - \mathbf{r}_o$. We will exploit this fact in the next section.

### 6 Transfer functions

**Equations S33 and S34** model how a spatial distribution of fluorescent dipoles $f(\mathbf{r}_o, \hat{\mathbf{s}}_o)$ appear in our irradiance measurements $g_{j\text{v}}(\mathbf{r}_d)$, but this model is computationally inefficient because it requires expensive integrals over $\mathbb{R}^3$ and $\mathbb{S}^2$. If we would like to efficiently simulate and invert this model, we must find a simpler form.

#### 6.1 Reformulating in terms of a transfer function

We will use two tools to rewrite our model in a more computationally efficient form. First, the *spatial Fourier transform* will let us exploit the shift-invariance and spatial band-limits of the imaging system so that we can turn the expensive convolution integral over $\mathbf{r}_o$ into an inexpensive FFT, multiplication, and inverse FFT. Second, the *spherical Fourier transform* will let us exploit the band-limited excitation and emission of dipolar fluorophores so that we can turn the expensive integral over $\hat{\mathbf{s}}_o$ into an inexpensive sum over just fifteen terms.

The first key result is that we can rewrite **Equation S33** as

$$G_{j\text{v}}(\boldsymbol{\nu}) = \sum_{\ell=0}^{\infty} \sum_{m=-\ell}^{\ell} \mathsf{H}_{j\text{v},\ell m}(\boldsymbol{\nu}) \mathsf{F}_{\ell m}(\boldsymbol{\nu}), \tag{S35}$$

where

$$G_{j\text{v}}(\boldsymbol{\nu}) = \int_{\mathbb{R}^3} d\mathbf{r}\, g_{j\text{v}}(\mathbf{r}) \exp(-2\pi i \mathbf{r} \cdot \boldsymbol{\nu}) \tag{S36}$$

is the *irradiance spectrum*,

$$\mathsf{H}_{j\text{v},\ell m}(\boldsymbol{\nu}) = \int_{\mathbb{R}^3} d\mathbf{r} \int_{\mathbb{S}^2} d\hat{\mathbf{s}}_o\; h_{j\text{v}}(\mathbf{r}, \hat{\mathbf{s}}_o) \exp(-2\pi i \mathbf{r} \cdot \boldsymbol{\nu}) Y_{\ell m}(\hat{\mathbf{s}}_o) \tag{S37}$$

is the *dipole spatio-angular transfer function*, and

$$\mathsf{F}_{\ell m}(\boldsymbol{\nu}) = \int_{\mathbb{R}^3} d\mathbf{r} \int_{\mathbb{S}^2} d\hat{\mathbf{s}}_o\; f(\mathbf{r}, \hat{\mathbf{s}}_o) \exp(-2\pi i \mathbf{r} \cdot \boldsymbol{\nu}) Y_{\ell m}(\hat{\mathbf{s}}_o) \tag{S38}$$

is the sample's *dipole spatio-angular spectrum*, $\boldsymbol{\nu} \in \mathbb{R}^3$ is a three-dimensional spatial-frequency coordinate, and $Y_{\ell m}(\hat{\mathbf{s}}_o)$ are the real spherical harmonic functions.

We have shown elsewhere [7] how to rewrite **Equation S33** in the form of **Equation S35**. Briefly, starting with **Equation S33** we apply the convolution-multiplication theorem then apply a generalized Plancherel theorem for spherical functions

$$\int_{\mathbb{S}^2} d\hat{\mathbf{s}}\, p(\hat{\mathbf{s}})q(\hat{\mathbf{s}}) = \sum_{\ell=0}^{\infty}\sum_{m=\ell}^{\ell} P_{\ell m}Q_{\ell m}, \tag{S39}$$

where $p(\hat{\mathbf{s}})$ and $q(\hat{\mathbf{s}})$ are arbitrary functions on the sphere, and $P_{\ell m}$ and $Q_{\ell m}$ are their spherical Fourier transforms defined by

$$P_{\ell m} = \int_{\mathbb{S}^2} d\hat{\mathbf{s}}\, p(\hat{\mathbf{s}})Y_{\ell m}(\hat{\mathbf{s}}). \tag{S40}$$

One way to understand the equivalence of **Equations S33 and S35** is that they both represent the same integral transform expressed in different bases—**Equation S33** in a standard basis of delta functions, and **Equation S35** in a basis of complex exponentials and spherical harmonics.

In the next section we will evaluate the integrals in **Equation S37** to find an explicit form for the dipole spatio-angular transfer function, but for now we will skip to a second key result: $\mathsf{H}_{j\mathrm{v},\ell m}(\boldsymbol{\mathfrak{v}})$ is only non-zero when $|\boldsymbol{\mathfrak{v}}_{\mathrm{v}}^{\perp}| < 2\mathrm{NA}_{\mathrm{v}}/\lambda$ and $\ell = 0, 2$ and 4. These limits are due to the transverse diffraction limit and the band-limited angular excitation and emission of dipolar fluorophores, and they let us further simplify **Equation S35** to a finite sum over a finite region in frequency space

$$G_{j\mathrm{v}}(\boldsymbol{\mathfrak{v}}) = \sum_{\ell=0,2,4}\sum_{m=-\ell}^{\ell} \mathsf{H}_{j\mathrm{v},\ell m}(\boldsymbol{\mathfrak{v}})\mathsf{F}_{\ell m}(\boldsymbol{\mathfrak{v}}) \quad \text{for} \quad |\boldsymbol{\mathfrak{v}}_{\mathrm{v}}^{\perp}| < 2\mathrm{NA}_{\mathrm{v}}/\lambda, \tag{S41}$$

a computationally efficient way to simulate and invert our model.

#### 6.2 Setting up the transfer function calculation

Our goal in this section is to evaluate the integrals in the dipole spatio-angular transfer function

$$\mathsf{H}_{j\mathrm{v},\ell m}(\boldsymbol{\mathfrak{v}}) = \int_{\mathbb{R}^3} d\boldsymbol{\mathfrak{r}} \int_{\mathbb{S}^2} d\hat{\mathbf{s}}_o\; h_{j\mathrm{v}}(\boldsymbol{\mathfrak{r}}, \hat{\mathbf{s}}_o)\exp(-2\pi i\boldsymbol{\mathfrak{r}}\cdot\boldsymbol{\mathfrak{v}})Y_{\ell m}(\hat{\mathbf{s}}_o), \tag{S42}$$

where $h_{j\mathrm{v}}(\boldsymbol{\mathfrak{r}}, \hat{\mathbf{s}}_o)$ is given by **Equation S34**. The details in this section are presented for those who wish to compute dipole spatio-angular transfer functions efficiently using Gaunt coefficients and Wigner D-matrices. Most readers can skip to **Supplement 6.5** for a summary of the results.

We start by plugging **Equation S34** into S42 and rearranging terms

$$\begin{aligned}\mathsf{H}_{j\mathrm{v},\ell m}(\boldsymbol{\mathfrak{v}}) = &\int_{\mathbb{R}^3} d\boldsymbol{\mathfrak{r}}\exp\left[-2\left(r_{\mathrm{v}}^{\parallel}/w_*\right)^2\right]\exp(-2\pi i\boldsymbol{\mathfrak{r}}\cdot\boldsymbol{\mathfrak{v}})\times\\ &\int_{\mathbb{S}^2} d\hat{\mathbf{s}}_o\, h_{j\mathrm{v}}^{(\text{exc, ang})}(\hat{\mathbf{s}}_o)\, h^{(\text{det},4f)}\left(\boldsymbol{\mathfrak{r}}_{\mathrm{v}}^{\perp}, 0, \hat{\mathbf{s}}_{\mathrm{v}}, \mathrm{NA}_{\mathrm{v}}\right) Y_{\ell m}(\hat{\mathbf{s}}_o).\end{aligned} \tag{S43}$$

Next, we split the three-dimensional coordinate $\mathfrak{r}$ into $(\mathbf{r}_{\text{v}}^{\perp}, r_{\text{v}}^{\parallel})$ then evaluate the axial integral and rearrange

$$\mathsf{H}_{j\text{v},\ell m}(\mathfrak{v}) = \frac{\exp\left[-\left(w_* v_{\text{v}}^{\parallel}\right)^2/2\right]}{\sqrt{\pi/2}/w_*} \int_{\mathbb{S}^2} d\hat{\mathbf{s}}_o\, h_{j\text{v}}^{(\text{exc, ang})}(\hat{\mathbf{s}}_o) Y_{\ell m}(\hat{\mathbf{s}}_o) \times \int_{\mathbb{R}^2} d\mathbf{r}_{\text{v}}^{\perp} h^{(\det,4f)}\left(\mathbf{r}_{\text{v}}^{\perp}, 0, \hat{\mathbf{s}}_{\text{v}}, \text{NA}_{\text{v}}\right) \exp(-2\pi i \mathbf{r}_{\text{v}}^{\perp} \cdot \mathbf{v}_{\text{v}}^{\perp}), \tag{S44}$$

where we have split the three-dimensional spatial frequency coordinate into its transverse and axial components $\mathfrak{v} = (\mathbf{v}_{\text{v}}^{\perp}, v_{\text{v}}^{\parallel})$. We collect the spatial integral into its own function, dropping the axial coordinate for convenience

$$H^{(\det,4f)}(\mathbf{v}_{\text{v}}^{\perp}, \hat{\mathbf{s}}_{\text{v}}, \text{NA}_{\text{v}}) = \int_{\mathbb{R}^2} d\mathbf{r}_{\text{v}}^{\perp} h^{(\det,4f)}\left(\mathbf{r}_{\text{v}}^{\perp}, 0, \hat{\mathbf{s}}_{\text{v}}, \text{NA}_{\text{v}}\right) \exp(-2\pi i \mathbf{r}_{\text{v}}^{\perp} \cdot \mathbf{v}_{\text{v}}^{\perp}), \tag{S45}$$

then rewrite the complete transfer function as

$$\mathsf{H}_{j\text{v},\ell m}(\mathfrak{v}) = \frac{\exp\left[-\left(w_* v_{\text{v}}^{\parallel}\right)^2/2\right]}{\sqrt{\pi/2}/w_*} \int_{\mathbb{S}^2} d\hat{\mathbf{s}}_o\, h_{j\text{v}}^{(\text{exc, ang})}(\hat{\mathbf{s}}_o) H^{(\det,4f)}(\mathbf{v}_{\text{v}}^{\perp}, \hat{\mathbf{s}}_{\text{v}}, \text{NA}_{\text{v}}) Y_{\ell m}(\hat{\mathbf{s}}_o). \tag{S46}$$

This spherical integral is challenging to evaluate, so we will evaluate it in pieces. First, we notice that this integral is a spherical Fourier transform of the product of two functions, which we can simplify by using the spherical version of the convolution-multiplication theorem

$$\int_{\mathbb{S}^2} d\hat{\mathbf{s}}\, p(\hat{\mathbf{s}}) q(\hat{\mathbf{s}}) Y_{\ell m}(\hat{\mathbf{s}}) = \sum_{\ell' m'} \sum_{\ell'' m''} \mathscr{G}_{\ell\ell'\ell''}^{mm'm''} P_{\ell'}^{m'} Q_{\ell''}^{m''}, \tag{S47}$$

where

$$P_{\ell'}^{m'} = \int_{\mathbb{S}^2} d\hat{\mathbf{s}}\, p(\hat{\mathbf{s}}) Y_{\ell' m'}(\hat{\mathbf{s}}), \tag{S48}$$

$$Q_{\ell''}^{m''} = \int_{\mathbb{S}^2} d\hat{\mathbf{s}}\, q(\hat{\mathbf{s}}) Y_{\ell'' m''}(\hat{\mathbf{s}}), \tag{S49}$$

and

$$\mathscr{G}_{\ell\ell'\ell''}^{mm'm''} = \int_{\mathbb{S}^2} d\hat{\mathbf{s}}\, Y_{\ell m}(\hat{\mathbf{s}}) Y_{\ell' m'}(\hat{\mathbf{s}}) Y_{\ell'' m''}(\hat{\mathbf{s}}) \tag{S50}$$

are the real Gaunt coefficients [12]. This identity lets us evaluate the spherical Fourier transform of the individual functions $h_{j\text{v}}^{(\text{exc, ang})}(\hat{\mathbf{s}}_o)$ and $h^{(\det,4f)}\left(\mathbf{r}_{\text{v}}^{\perp}, 0, \hat{\mathbf{s}}_{\text{v}}, \text{NA}_{\text{v}}\right)$, then combine them using **Equation S47** to find the spherical Fourier transform of their product. We will evaluate these spherical Fourier transforms in the next two section before we complete the transfer function calculation.

#### 6.3 Angular excitation transfer function

In this section we evaluate the following integral

$$
\begin{aligned}
H_{j\mathrm{v},\ell m}^{(\mathrm{exc,\ ang})} &\equiv \int_{\mathbb{S}^2} d\hat{\mathbf{s}}_o\, h_{j\mathrm{v}}^{(\mathrm{exc,\ ang})}(\hat{\mathbf{s}}_o) Y_{\ell m}(\hat{\mathbf{s}}_o) \\
&= \frac{3}{\sqrt{4\pi}} \int_{\mathbb{S}^2} d\hat{\mathbf{s}}_o\, (\hat{\mathbf{p}}_{j\mathrm{v}} \cdot \hat{\mathbf{s}}_o)^2 Y_{\ell m}(\hat{\mathbf{s}}_o) \\
&= \frac{1}{\sqrt{4\pi}} \int_{\mathbb{S}^2} d\hat{\mathbf{s}}_o\, [P_0(\hat{\mathbf{p}}_{j\mathrm{v}} \cdot \hat{\mathbf{s}}_o) + 2P_2(\hat{\mathbf{p}}_{j\mathrm{v}} \cdot \hat{\mathbf{s}}_o)] Y_{\ell m}(\hat{\mathbf{s}}_o) \\
&= \sqrt{4\pi} \int_{\mathbb{S}^2} d\hat{\mathbf{s}}_o \left[ Y_{00}(\hat{\mathbf{p}}_{j\mathrm{v}}) Y_{00}(\hat{\mathbf{s}}_o) + \frac{2}{5} \sum_{m'=-2}^{2} Y_{2m'}(\hat{\mathbf{p}}_{j\mathrm{v}}) Y_{2m'}(\hat{\mathbf{s}}_o) \right] Y_{\ell m}(\hat{\mathbf{s}}_o) \\
&= \sqrt{4\pi} \left[ Y_{00}(\hat{\mathbf{p}}_{j\mathrm{v}}) \delta_{0\ell} + \frac{2}{5} \sum_{m'=-2}^{2} Y_{2m'}(\hat{\mathbf{p}}_{j\mathrm{v}}) \delta_{2\ell} \delta_{mm'} \right] \\
&= \sqrt{4\pi}\, Y_{\ell m}(\hat{\mathbf{p}}_{j\mathrm{v}}) \left( \delta_{0\ell} + \frac{2}{5} \delta_{2\ell} \right),
\end{aligned}
\tag{S51}
$$

where we have expanded in terms of Legendre polynomials $P_\ell(x)$, applied the spherical harmonic addition theorem $P_\ell(\hat{\mathbf{x}} \cdot \hat{\mathbf{y}}) = \frac{4\pi}{2\ell+1} \sum_{m'=-\ell}^{\ell} Y_{\ell m'}(\hat{\mathbf{x}}) Y_{\ell m'}(\hat{\mathbf{y}})$, exploited the orthonormality of the spherical harmonics $\int_{\mathbb{S}^2} d\hat{\mathbf{s}} Y_{\ell m}(\hat{\mathbf{s}}_o) Y_{\ell' m'}(\hat{\mathbf{s}}_o) = \delta_{\ell\ell'} \delta_{mm'}$ where $\delta_{\ell\ell'}$ is the Kronecker delta, then used the discrete sifting theorem $\sum_{m'} f_{m'} \delta_{mm'} = f_m$. **Equation S51** shows that the angular excitation transfer function contains at most six non-zero terms (one for $\ell = 0$, five for $\ell = 2$), and these terms can be found efficiently by evaluating the spherical harmonic functions along the illumination polarization axes $\hat{\mathbf{p}}_{j\mathrm{v}}$. We will make explicit choices for our illumination polarizations in **Supplement 8.3**.

#### 6.4 Detection transfer function

In this section we evaluate the following integral

$$
\mathsf{H}_{\mathrm{v},\ell m}^{(\mathrm{det},4f)}(\mathbf{v}_{\mathrm{v}}^{\perp}, \mathrm{NA}_{\mathrm{v}}) \equiv \int_{\mathbb{S}^2} d\hat{\mathbf{s}}_o\, H^{(\mathrm{det},4f)}\left(\mathbf{v}_{\mathrm{v}}^{\perp}, \mathbf{R}_{\mathrm{v}}^{-1} \hat{\mathbf{s}}_o, \mathrm{NA}_{\mathrm{v}}\right) Y_{\ell m}(\hat{\mathbf{s}}_o), \tag{S52}
$$

where we have explicitly written the angular coordinate as $\hat{\mathbf{s}}_{\mathrm{v}} = \mathbf{R}_{\mathrm{v}}^{-1} \hat{\mathbf{s}}_o$ so that we can evaluate the integral for both views.

In the same way that the Fourier-shift theorem can simplify spatial Fourier transforms, here we seek an analogous simplification that will let us efficiently compute the spherical Fourier transform of a rotated function. Suppose we have a spherical function $f(\hat{\mathbf{s}})$ and its spherical Fourier transform

$$F_{\ell m} = \int_{\mathbb{S}^2} d\hat{\mathbf{s}}\, f(\hat{\mathbf{s}}) Y_{\ell m}(\hat{\mathbf{s}}). \tag{S53}$$

The spherical Fourier transform of same function in rotated coordinates is given by

$$F'_{\ell' m'} = \int_{\mathbb{S}^2} d\hat{\mathbf{s}}\, f(\mathbf{R}^{-1}\hat{\mathbf{s}}) Y_{\ell' m'}(\hat{\mathbf{s}}), \tag{S54}$$

where $\mathbf{R} \in \mathbb{SO}(3)$ is a rotation matrix. After making a change of coordinates $\mathbf{R}^{-1}\hat{\mathbf{s}} \to \hat{\mathbf{s}}$

$$F'_{\ell' m'} = \int_{\mathbb{S}^2} d\hat{\mathbf{s}}\, f(\hat{\mathbf{s}}) Y_{\ell' m'}(\mathbf{R}\hat{\mathbf{s}}), \tag{S55}$$

we expand $f(\hat{\mathbf{s}})$ into a spherical-harmonic series

$$F'_{\ell' m'} = \int_{\mathbb{S}^2} d\hat{\mathbf{s}} \left[ \sum_{\ell=0}^{\infty} \sum_{m=-\ell}^{\ell} F_{\ell m} Y_{\ell m}(\hat{\mathbf{s}}) \right] Y_{\ell' m'}(\mathbf{R}\hat{\mathbf{s}}), \tag{S56}$$

then rearrange to find

$$F'_{\ell' m'} = \sum_{\ell=0}^{\infty} \sum_{m=-\ell}^{\ell} \left[ \int_{\mathbb{S}^2} d\hat{\mathbf{s}}\, Y_{\ell m}(\hat{\mathbf{s}}) Y_{\ell' m'}(\mathbf{R}\hat{\mathbf{s}}) \right] F_{\ell m}. \tag{S57}$$

The integral in square brackets is only non-zero when $\ell = \ell'$, so we perform the sum over $\ell$ and give the integral its own symbol

$$F'_{\ell m'} = \sum_{m=-\ell}^{\ell} \Delta^{\ell}_{mm'}(\mathbf{R}) F_{\ell m}\ , \tag{S58}$$

where

$$\Delta^{\ell}_{mm'}(\mathbf{R}) = \int_{\mathbb{S}^2} d\hat{\mathbf{s}}\, Y_{\ell m}(\hat{\mathbf{s}}) Y_{\ell m'}(\mathbf{R}\hat{\mathbf{s}}) \tag{S59}$$

are the *real Wigner D-matrices*—see the appendix in Kautz et al. for a similar result [13]. The Wigner D-matrices are square $(2\ell + 1) \times (2\ell + 1)$ matrices for each rotation $\mathbf{R}$ and band $\ell$, and these matrices can be used to calculate the spherical harmonic coefficients of a rotated function. Notice that rotations only change the spherical harmonic coefficients within each band since each band of spherical harmonics spans an $(2\ell + 1)$-dimensional rotationally invariant subspace of $\mathbb{L}_2(\mathbb{S}^2)$.

We can apply **Equation S58** to simplify our target integral **Equation S52** into

$$\mathsf{H}_{\ell m}^{(\text{det},4f)}(\mathbf{v}_\text{v}^\perp, \text{NA}_\text{v}) = \sum_{m=-\ell}^{\ell} \Delta^\ell_{mm'}(\mathbf{R}_\text{v}) \int_{\mathbb{S}^2} d\hat{\mathbf{s}}_o \, H^{(\text{det},4f)}\big(\mathbf{v}_\text{v}^\perp, 0, \hat{\mathbf{s}}_o, \text{NA}_\text{v}\big)\, Y_{\ell m}(\hat{\mathbf{s}}_o). \tag{S60}$$

Our only remaining task is to evaluate the spherical Fourier transform of $h^{(\text{det},4f)}\big(\mathbf{r}_\text{v}^\perp, 0, \hat{\mathbf{s}}_o, \text{NA}_\text{v}\big)$. Proceeding step by step, we start with our target integral

$$= \int_{\mathbb{S}^2} d\hat{\mathbf{s}}_o \, h^{(\text{det},4f)}\big(\mathbf{r}_\text{v}^\perp, 0, \hat{\mathbf{s}}_o, \text{NA}_\text{v}\big)\, Y_{\ell m}(\hat{\mathbf{s}}_o), \tag{S61}$$

substitute **Equation S23** and separate the angular integral

$$= \sum_{nn'=0,1,2} \left[\int_{\mathbb{S}^2} d\hat{\mathbf{s}}_o \, s_n s_{n'} Y_{\ell m}(\hat{\mathbf{s}}_o)\right] b_{nn'}(\mathbf{r}_\text{v}^\perp, 0, \text{NA}_\text{v}), \tag{S62}$$

then notice that we can rewrite the integral in terms of the Gaunt coefficients

$$= \frac{4\pi}{3} \sum_{nn'=0,1,2} \mathscr{G}_{\ell 11}^{m\epsilon_n\epsilon_{n'}} b_{nn'}(\mathbf{r}_\text{v}^\perp, 0, \text{NA}_\text{v}), \tag{S63}$$

where $\epsilon_0 = 1$, $\epsilon_1 = -1$, $\epsilon_2 = 0$.

We can now complete our calculation of the detection transfer function by substituting **Equation S63** into **Equation S60** to give the main result for this section

$$\mathsf{H}_{\ell m}^{(\text{det},4f)}(\mathbf{r}_\text{v}^\perp, \text{NA}_\text{v}) = \frac{4\pi}{3} \sum_{m=-\ell}^{\ell} \Delta^\ell_{mm'}(\mathbf{R}_\text{v}) \sum_{nn'=0,1,2} \mathscr{G}_{\ell 11}^{m\epsilon_n\epsilon_{n'}} B_{nn'}(\mathbf{r}_\text{v}^\perp, 0, \text{NA}_\text{v}). \tag{S64}$$

Calculating $\Delta^\ell_{mm'}(\mathbf{R})$ efficiently for arbitrary $\mathbf{R}$, $\ell$, $m$, and $m'$ is challenging—see Pinchon and Hoggan for one approach [14]. Fortunately, we only have two rotation matrices $\mathbf{R}_\text{v}$, and $\mathscr{G}_{\ell 11}^{m\epsilon_n\epsilon_{n'}}$ is only non-zero for $\ell = 0$ and $\ell = 2$ terms (see Homeier for Gaunt coefficient selection rules [12]). This means we only need to calculate $2(1^2+5^2) = 52$ integrals, which is feasible symbolically. Most of these integrals are trivial and can be calculated by hand

$$\Delta^0_{00}(\mathbf{R}_\text{v}) = 1, \quad \Delta^2_{mm'}(\mathbf{R}_A) = \delta_{mm'}. \tag{S65}$$

The remaining integrals can be evaluated with a computer algebra package, and we write the values $\Delta^2_{mm'}(\mathbf{R}_B)$ in matrix notation as

$$\boldsymbol{\Delta}^2(\mathbf{R}_B) = \begin{bmatrix} 0 & -1 & 0 & 0 & 0 \\ -1 & 0 & 0 & 0 & 0 \\ 0 & 0 & -1/2 & 0 & \sqrt{3}/2 \\ 0 & 0 & 0 & 1 & 0 \\ 0 & 0 & \sqrt{3}/2 & 0 & 1/2 \end{bmatrix}. \tag{S66}$$

As expected this matrix is involutary $[\boldsymbol{\Delta}^2(\mathbf{R}_B)]^{-1} = \boldsymbol{\Delta}^2(\mathbf{R}_B)$, because the matrix $\mathbf{R}_B$ is involutary.

##### 6.5 Complete spatio-angular transfer function

We now have all of the pieces for our complete spatio-angular transfer function.

$$\mathsf{H}_{j\mathsf{v},\ell m}(\boldsymbol{\mathfrak{v}}) = \frac{\exp\left[-\left(w_* v_{\mathsf{v}}^{\parallel}\right)^2/2\right]}{\sqrt{\pi/2}/w_*} \sum_{\ell' m'} \sum_{\ell'' m''} \mathscr{G}_{\ell\ell'\ell''}^{mm'm''} H_{j\mathsf{v},\ell' m'}^{(\text{exc, ang})} \mathsf{H}_{\ell'' m''}^{(\text{det},4f)}(\mathbf{v}_{\mathsf{v}}^{\perp}, \text{NA}_{\mathsf{v}}), \tag{S67}$$

where

$$H_{j\mathsf{v},\ell m}^{(\text{exc, ang})} = \sqrt{4\pi}\, Y_{\ell m}(\hat{\mathbf{p}}_{j\mathsf{v}}) \left(\delta_{0\ell} + \frac{2}{5}\delta_{2\ell}\right), \tag{S68}$$

and

$$\mathsf{H}_{\ell m}^{(\text{det},4f)}(\mathbf{r}_{\mathsf{v}}^{\perp}, \text{NA}_{\mathsf{v}}) = \frac{4\pi}{3} \sum_{m=-\ell}^{\ell} \Delta_{mm'}^{\ell}(\mathbf{R}_{\mathsf{v}}) \sum_{nn'=0,1,2} \mathscr{G}_{\ell 11}^{m\epsilon_n\epsilon_{n'}} b_{nn'}(\mathbf{r}_{\mathsf{v}}^{\perp}, 0, \text{NA}_{\mathsf{v}}). \tag{S69}$$

From our previous work we know that the excitation and detection transfer functions are only non-zero for $\ell = 0$, 2, so the complete transfer function is only non-zero for $\ell = 0$, 2, 4, which is at most 15 non-zero angular terms.

### 7 Reconstruction algorithm

##### 7.1 Theoretical motivation

The spatio-angular transfer function $\mathsf{H}_{j\mathsf{v},\ell m}(\boldsymbol{\mathfrak{v}})$ tells us how an object's spatio-angular spectrum $\mathsf{F}_{\ell m}(\boldsymbol{\mathfrak{v}})$ is transmitted to the data's spectrum $G_{j\mathsf{v}}(\boldsymbol{\mathfrak{v}})$ with the following relationship

$$G_{j\mathsf{v}}(\boldsymbol{\mathfrak{v}}) = \sum_{\ell m} \mathsf{H}_{j\mathsf{v},\ell m}(\boldsymbol{\mathfrak{v}}) \mathsf{F}_{\ell m}(\boldsymbol{\mathfrak{v}}). \tag{S70}$$

Our goal in this section is to find an efficient way to estimate the object's spatio-angular spectrum, $\hat{\mathsf{F}}_{\ell m}(\boldsymbol{\mathfrak{v}})$, from a noise-corrupted measurement of the data spectrum.

**Equation S70** shows that $\mathsf{H}_{j\mathsf{v},\ell m}(\boldsymbol{\mathfrak{v}})$ can be interpreted as a matrix for each spatial frequency $\boldsymbol{\mathfrak{v}}$, with rows indexed by $j\mathsf{v}$ and columns indexed by $\ell m$. This observation lets us temporarily lift the notational burden of coordinates to rewrite **Equation S70** in matrix-vector form

$$\mathbf{g} = \mathcal{H}\mathbf{f}, \tag{S71}$$

where the matrix multiplication is implied.

A reasonable starting place for estimating the object $\mathbf{f}$ from the data $\mathbf{g}$ is to solve the least-squares optimization problem

$$\hat{\mathbf{f}}^{\,(\mathrm{LS})} = \operatorname*{argmin}_{\mathbf{f}} |\mathbf{g} - \mathcal{H}\mathbf{f}|^2, \tag{S72}$$

where $|\mathbf{g}|$ is the $\mathbb{L}^2$ norm of $\mathbf{g}$. This optimization problem has a closed-form solution that is most easily expressed in terms of the singular system of $\mathcal{H}$

$$\hat{\mathbf{f}}^{\,(\mathrm{LS})} = \sum_{k=1}^{R} \frac{1}{\sqrt{\mu_k}} \mathbf{u}_k \left(\mathbf{v}_k \cdot \mathbf{g}\right), \tag{S73}$$

where $R$ is the rank of $\mathcal{H}$, $(\mathbf{v}_k \cdot \mathbf{g})$ is an inner product between vectors, and $(\mu_k, \mathbf{u}_k, \mathbf{v}_k)$ is the singular system of $\mathcal{H}$ that satisfies

$$\mathcal{H}^T\mathcal{H}\mathbf{u}_k = \mu_k\mathbf{u}_k, \tag{S74}$$

$$\mathcal{H}\mathcal{H}^T\mathbf{v}_k = \mu_k\mathbf{v}_k, \tag{S75}$$

where $\mathcal{H}^T$ is the transpose of $\mathcal{H}$. This solution can be statistically justified as the maximum-likelihood estimator for data corrupted by uncorrelated Gaussian noise [15, ch. 13.3.4], but in practice division by small eigenvalues $\mu_k$ can amplify noise to unacceptable levels.

We address this problem by adding a *Tikhonov regularization* term to the optimization problem

$$\hat{\mathbf{f}}^{\,\eta} = \operatorname*{argmin}_{\mathbf{f}} |\mathbf{g} - \mathcal{H}\mathbf{f}|^2 + \eta|\mathbf{f}|^2, \tag{S76}$$

where $\eta$ is a positive constant. Once again, this optimization problem has a closed-form solution in terms of the singular system of $\mathcal{H}$

$$\hat{\mathbf{f}}^{\,\eta} = \sum_{k=1}^{R} \frac{\sqrt{\mu_k}}{\mu_k + \eta} \mathbf{u}_k \left(\mathbf{v}_k \cdot \mathbf{g}\right). \tag{S77}$$

Statistically, adding a Tikhonov regularizer can be interpreted as applying a Bayesian prior that assumes the unknown parameters to be independent, zero-mean, Gaussian-distributed random variables with variance $1/(2\eta)$ [15, ch. 15.3.3]. Even when this assumption is not strictly true, adding a Tikhonov regularizer is a practical way to control noise amplification.

We can rewrite **Equation S77** with coordinates as

$$\hat{\mathsf{F}}^{\eta}_{\ell m}(\mathfrak{v}) = \sum_{k=1}^{R} \frac{\sqrt{\mu_k(\mathfrak{v})}}{\mu_k(\mathfrak{v}) + \eta} U_{k,\ell m}(\mathfrak{v}) \sum_{j\mathtt{v}} V_{k,j\mathtt{v}}(\mathfrak{v}) G_{j\mathtt{v}}(\mathfrak{v}), \tag{S78}$$

where $U_{k,\ell m}(\mathfrak{v})$ and $V_{k,j\mathtt{v}}(\mathfrak{v})$ are matrices with rows consisting of the right- and left-singular vectors of the spatio-angular transfer function $\mathsf{H}_{j\mathtt{v},\ell m}(\mathfrak{v})$, respectively.

#### 7.2 Practical description of the reconstruction algorithm

After deskewing and registering our raw volumes (see **Supplement 4**), we renormalize each volume using our initial calibration measurements (see **Supplement 3.2**). **Figure S7** shows an example of volume-averaged calibration measurements from a fluorescent lake. We found that our calibration data showed the expected $\cos^2\theta$-type intensity variation across illumination polarizations, but we found the intensity variations between tilts and views to vary between experimental runs, driven by changes in alignment of the two arms. Additionally, we found our imaging configuration introduced a $\sim$ 10-15 degree polarization shift compared to the alignment configuration (**Figure S4**), measurable by fitting curves to the calibration data points. We suspect the detection-side dichroic filter is responsible for this polarization phase shift.

To correct for these effects we applied volume-wise calibration factors to the raw data. First, we used the curve-fit polarization phase shift and our model of the imaging system to calculate an expected set of intensities from a lake $H^{(\text{cal})}_{j\mathtt{v},00}(\mathbf{0})$. Second, we calculated volume-averaged measurements $\bar{g}^{(\text{cal})}_{j\mathtt{v}}$ from our calibration data (see **Supplement 3.2**). Finally, we reasoned that we should apply normalized calibration factors to each volume, so we normalized both terms by their first $j\mathtt{v}$ terms, specifically, $\bar{g}^{(\text{cal})}_{00}$ and $H^{(\text{cal})}_{00,00}(\mathbf{0})$. Altogether, our calibration correction takes the form

$$g_{j\mathtt{v}}(\mathbf{r}_d) = g^{(\text{raw})}_{j\mathtt{v}}(\mathbf{r}_d) \frac{\bar{g}^{(\text{cal})}_{00}}{\bar{g}^{(\text{cal})}_{j\mathtt{v}}} \frac{H^{(\text{cal})}_{j\mathtt{v},00}(\mathbf{0})}{H^{(\text{cal})}_{00,00}(\mathbf{0})}. \tag{S79}$$

**Equation S79** applies a volume-wise correction to bridge the gap between our model of the instrument and our calibration measurements taken with a known sample. **Figure S8** shows the marginal improvement that the calibration procedure makes on the reconstructions we report here. In early iterations of the instrument we found the calibration procedure to be essential, but as the instrument stabilized and we refined our models we found the calibration procedure to be less necessary.

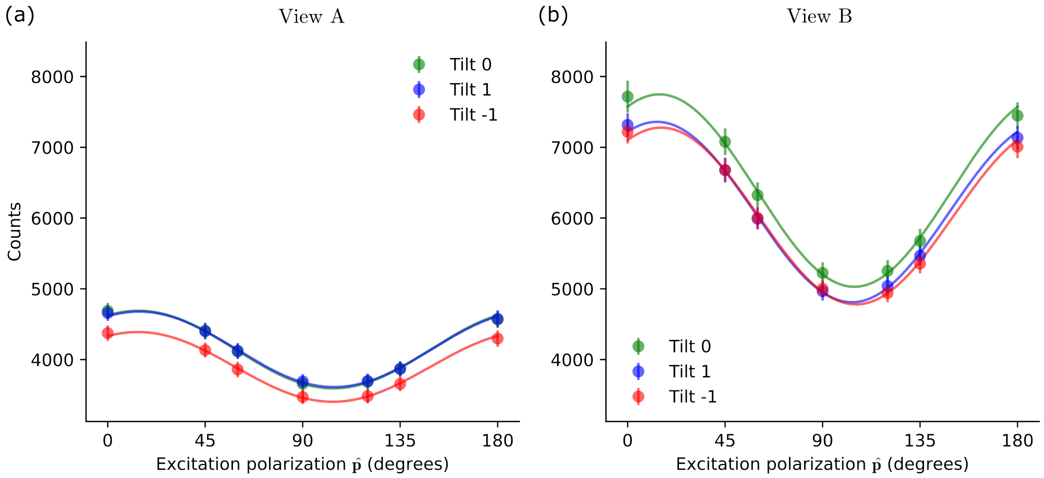


**Fig. S7** Volume-averaged calibration measurements from a fluorescent lake. We acquired volumes under seven polarization (dots), three different tilt angles (colors) with (a) view A and (b) view B. Each data point is the mean over an $\sim 3.5{\times}3.5{\times}3.5\ \mu\text{m}^3$ volume from deep within the fluorescent lake, and the error bars indicate the standard deviation of intensity values across the volume. For each set of polarization measurements we fit a curve of the form $y = a\cos^2(x - b) + c$ (solid lines), then averaged the $b$ estimates across tilts and views to estimate the system's polarization phase shift, here $\sim 11$ degrees corresponding to the peak of the fit.

Next, we take the three-dimensional Fourier transform of each calibration-corrected volume

$$G_{j\mathrm{v}}(\boldsymbol{\mathfrak{v}}) = \int_{\mathbb{R}^3} d\boldsymbol{\mathfrak{r}}_d\, g_{j\mathrm{v}}(\boldsymbol{\mathfrak{r}}_d) \exp[-2\pi i \boldsymbol{\mathfrak{r}}_d \cdot \boldsymbol{\mathfrak{v}}], \tag{S80}$$

apply a Tikhonov-regularized pseudoinverse using a pre-computed singular system, see **Supplement 7.1** for theoretical details and **Supplement 7.3** for practical tips,

$$\hat{\mathsf{F}}^{\eta}_{\ell m}(\boldsymbol{\mathfrak{v}}) = \sum_{k=1}^{R} \frac{\sqrt{\mu_k(\boldsymbol{\mathfrak{v}})}}{\mu_k(\boldsymbol{\mathfrak{v}}) + \eta} U_{k,\ell m}(\boldsymbol{\mathfrak{v}}) \sum_{j\mathrm{v}} V_{k,j\mathrm{v}}(\boldsymbol{\mathfrak{v}}) G_{j\mathrm{v}}(\boldsymbol{\mathfrak{v}}), \tag{S81}$$

then we take an inverse three-dimensional Fourier transform

$$\hat{F}^{\eta}_{\ell m}(\boldsymbol{\mathfrak{r}}_o) = \int_{\mathbb{R}^3} d\boldsymbol{\mathfrak{v}}\, \hat{\mathsf{F}}^{\eta}_{\ell m}(\boldsymbol{\mathfrak{v}}) \exp[2\pi i \boldsymbol{\mathfrak{r}}_o \cdot \boldsymbol{\mathfrak{v}}], \tag{S82}$$

and store the result. At visualization time, we complete our reconstruction by calculating the spatio-angular Boltzmann distribution with

$$\hat{f}^{\,\eta}(\boldsymbol{\mathfrak{r}}_o, \hat{\mathbf{s}}_o) = \sum_{\ell m} \hat{F}^{\eta}_{\ell m}(\boldsymbol{\mathfrak{r}}_o) Y_{\ell m}(\hat{\mathbf{s}}_o). \tag{S83}$$

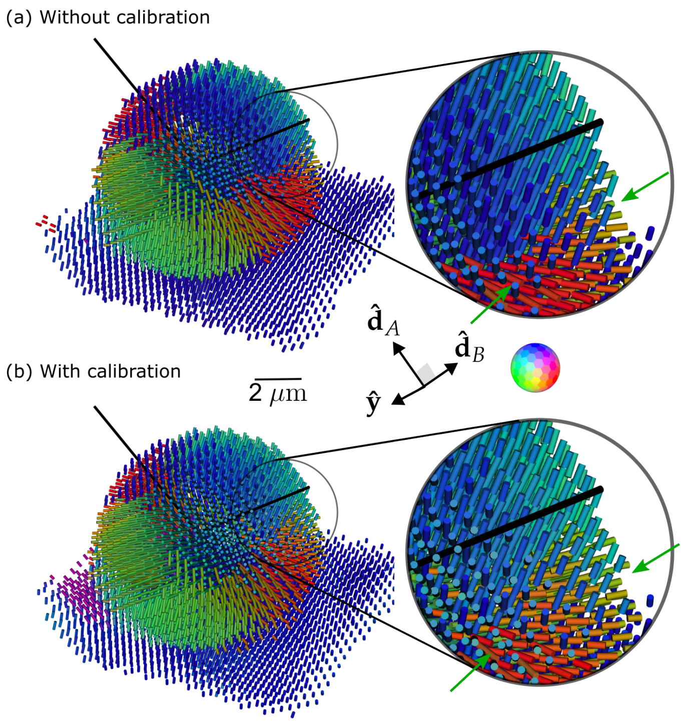


**Fig. S8** GUV peak reconstruction from "All" polarization measurements (a) without and (b) with applying the calibration algorithm. We find that when the measurements match the model well, the calibration procedure makes only marginal improvements on the reconstruction, see inset and green arrows where the transition from blue to red orientations is smoother with calibration.

#### 7.3 Practical precomputations

Our datasets commonly reach spatial sizes of $1000 \times 1000 \times 1000 = 10^9$ voxels, so 6 acquired volumes can fill $6 \times (4 \text{ bytes/value}) \times (10^9) \approx 24$ GB. If we naively precomputed the entire singular system, we would need to store 15 spherical harmonic coefficients, 6 data-space coefficients, and 1 singular value for each of the 6 non-zero singular values at each spatial frequency totalling $6(15+6+1)\times(10^9)\times(4 \text{ bytes/value}) \approx$ 500 GB of data to perform a reconstruction. This is a significant burden that can be alleviated with on-the-fly computation of the transfer functions.

First, we exploit the separability of the transfer function to compute and store a small number of values that can be combined to generate all of the entries $\mathsf{H}_{j\mathrm{v},\ell m}(\boldsymbol{\mathfrak{v}})$. **Equation** S67 shows a natural way to decompose the complete transfer function into five parts—an spatial excitation part, an angular excitation part, a detection part, real Gaunt coefficients, and real Wigner D-matrices—and each of these can be precalculated, stored, and combined efficiently.

At reconstruction time, we use **Equation** S67 to combine our stored values into a complete transfer function at a given spatial frequency. We can take this $6 \times 15$ matrix, inexpensively compute its singular value decomposition, perform the reconstruction

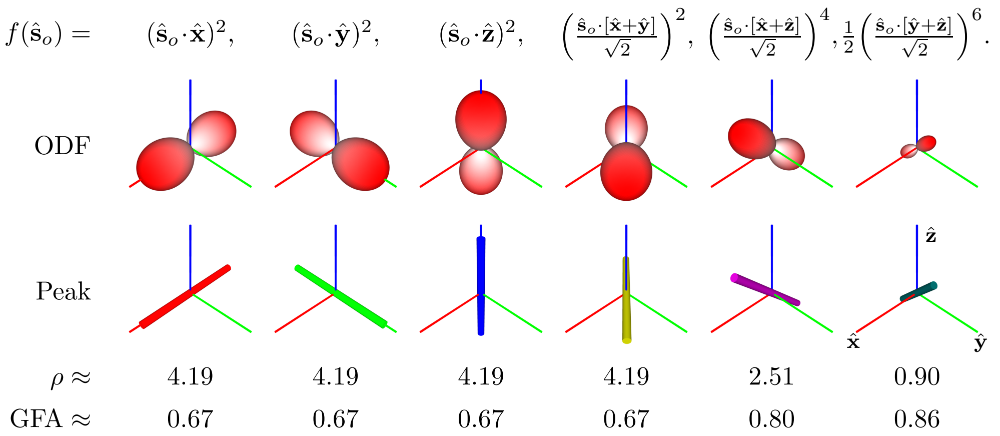


**Fig. S9** Demonstration of angular visualization schemes. Each column shows a different angular function for a single spatial point $f(\hat{\mathbf{s}}_o)$ specified by the upper label. The first row shows an orientation distribution function (ODF) visualization that doubly encodes the magnitude of $f(\hat{\mathbf{s}}_o)$ into the glyph radius and glyph color. Notice that red encodes the maximum for each glyph instead of a fixed value. The second row shows the peak directions visualized with an oriented cylinder colored using the absolute value method. Once again, the peak orientation is doubly encoded into the orientation and color of the cylinder. Finally, we calculate the density $\rho$ and the generalized fractional anisotropy (GFA). These scalar values can be visualized using any color map.

using **Equation S81**, then repeat the process on the next spatial frequency. This approach reduces the precomputation storage burden while increasing computational demands at reconstruction time.

#### 7.4 Orientation distribution functions and summary statistics

In this section we look at several ways to visualize and summarize the spatio-angular Boltzmann distributions that we estimate $\hat{f}^{\eta}(\mathbf{r}_o, \hat{\mathbf{s}}_o)$. **Figure S9** summarizes four ways to visualize individual spatial points from a spatio-angular reconstruction. We have found these visuals to be the most useful for understanding and interpreting reconstructions.

Starting with the stored reconstruction $\hat{F}^{\eta}_{\ell m}(\mathbf{r}_o)$, our goal is to calculate and plot useful visuals. To avoid storage inflation we have postponed our conversion to a standard basis until visualization time. Our first step is to choose a set of $\mathtt{N}$ points on the sphere $\{\hat{\mathbf{s}}_{o,\mathtt{n}}\}$ that we would like to visualize. The Fibonacci lattice is an attractive choice because it leads to well-spaced points that are inexpensive to compute for arbitrary $\mathtt{N}$. The polar angles $\{\theta_\mathtt{n}\}$ and azimuthal angles $\{\phi_\mathtt{n}\}$ of the Fibonacci lattice with $\mathtt{N}$ points are given by [16, 17]

$$\theta_\mathtt{n} = \cos^{-1}(1 - (2\mathtt{n} + 1)/\mathtt{N}), \tag{S84}$$

$$\phi_\mathtt{n} = \pi(3 - \sqrt{5})\mathtt{n}. \tag{S85}$$

Choosing a larger `N` will make the final visuals appear smoother at additional computational expense. Empirically we have found that `N = 500` is an appropriate starting point for most visualizations.

Next, we choose a set of spatial points $\{\mathbf{r}_{o,\text{n}}\}$ where we would like to visualize the object. We recommend starting with a modest number of spatial points by downsampling or thresholding the reconstruction because visualizing every spatial point is visually overwhelming and computationally expensive. Empirically we have found that visualizing more than $10^4$ spatial points overwhelms most users who are trying to interpret the results and most computers that are trying to render them without a dedicated graphics card.

With our spherical points $\{\hat{\mathbf{s}}_{o,\text{n}}\}$ and spatial points $\{\mathbf{r}_{o,\text{n}}\}$ we can calculate the *orientation distribution functions (ODFs)* at each point

$$\hat{f}^{\eta}(\mathbf{r}_{o,\text{n}}, \hat{\mathbf{s}}_{o,\text{n}}) = \sum_{\ell m} \hat{F}^{\eta}_{\ell m}(\mathbf{r}_{o,\text{n}}) Y_{\ell m}(\hat{\mathbf{s}}_{o,\text{n}}). \tag{S86}$$

Notice that the spherical harmonics $Y_{\ell m}(\hat{\mathbf{s}}_{o,\text{n}})$ can be computed once then reused. The ODFs can be visualized by drawing a glyph at each point $\{\mathbf{r}_{o,\text{n}}\}$ with a radius along each direction $\{\hat{\mathbf{s}}_{o,\text{n}}\}$ given by $\hat{f}^{\eta}(\mathbf{r}_{o,\text{n}}, \mathbf{s}_{o,\text{n}})$. We use a blue-white-red color map to doubly encode the value of $\hat{f}^{\eta}(\mathbf{r}_{o,\text{n}}, \hat{\mathbf{s}}_{o,\text{n}})$ into the radius and the color of the glyph.

In addition to drawing a complete glyph at each point $\{\mathbf{r}_{o,\text{n}}\}$, we have found that drawing a cylinder or thin line at each point $\{\mathbf{r}_{o,\text{n}}\}$ along the direction where the function is largest

$$\hat{\mathbf{s}}^{\eta,(\text{pk})}(\mathbf{r}_{o,\text{n}}) = \underset{\hat{\mathbf{s}}_o}{\text{argmax}}\, \hat{f}^{\eta}(\mathbf{r}_{o,\text{n}}, \hat{\mathbf{s}}_o) \tag{S87}$$

is a good way to summarize and understand the reconstruction. In many cases the viewing direction obscures the direction of the cylinder or line, so encoding the direction of the line in color can reduce visual degeneracy. In all of our reconstructions we have colored the $\hat{\mathbf{x}}$, $\hat{\mathbf{y}}$, and $\hat{\mathbf{z}}$ components of $\hat{\mathbf{s}}^{\eta,(\text{pk})}(\mathbf{r}_{o,\text{n}})$ with weighted red, green, and blue color channels, respectively. In the computer graphics and magnetic resonance imaging (MRI) literature this color mapping is usually called the *absolute value method* [18]. Although the absolute value method is widely used and easy to understand, it suffers from ambiguities that can be avoided by using more sophisticated color maps [19].

We can also calculate and plot scalar summary statistics for each spatial point. The estimated number of fluorophores at each point or *fluorophore density* is given directly by

$$\hat{\rho}^{\eta}(\mathbf{r}_o) = \hat{F}^{\eta}_{00}(\mathbf{r}_o). \tag{S88}$$

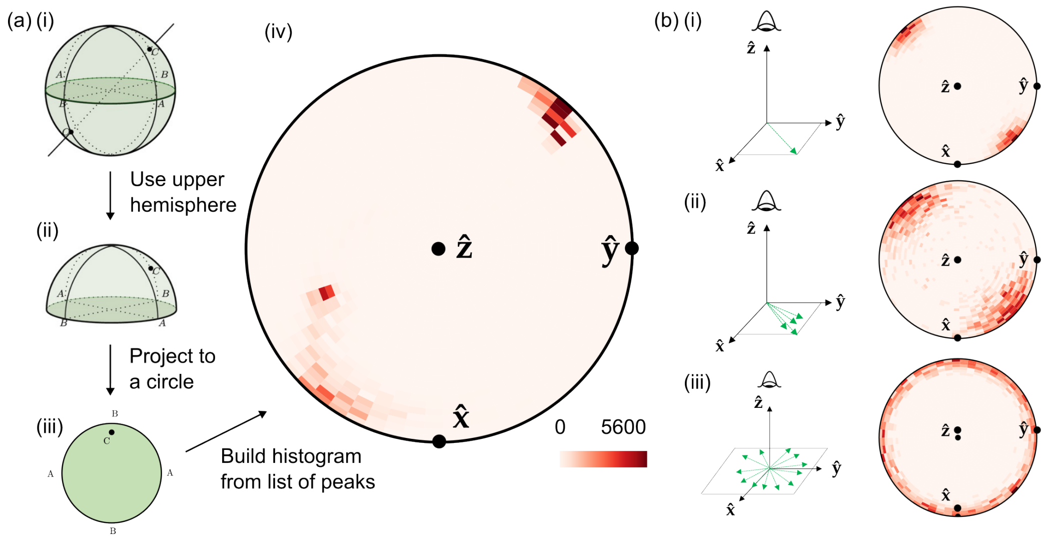


**Fig. S10 Peak histograms.** We build a peak histogram by (i) starting with a list of normalized 3D vectors, e.g. A, B, and C. A 3D vector and its antipodal point vector represent the same peak (e.g. A, B, and C appear twice and represent the same peak), so we an choose a viewing direction and (ii) flip vectors to the upper hemisphere to remove the ambiguity, then (iii) project the 3D vector to a 2D circle, then (iv) increment bins for each vector in the list. (b) Example histograms of dipole distributions viewed along the $\hat{\mathbf{z}}$ axis. Left, cartoons illustrating a small number of a draws from an example distribution; right, simulated histograms with 1000 samples from Watson distributions $f(\hat{\mathbf{s}}) \propto \exp\left[\kappa(\hat{\boldsymbol{\mu}} \cdot \hat{\mathbf{s}})\right]$, where $\hat{\boldsymbol{\mu}}$ is a direction and $\kappa$ is a spread parameter. (i) $\kappa = 20$, $\hat{\boldsymbol{\mu}} = (\hat{\mathbf{x}} + \hat{\mathbf{y}})/\sqrt{2}$ (ii) $\kappa = 5$, $\hat{\boldsymbol{\mu}} = (\hat{\mathbf{x}} + \hat{\mathbf{y}})/\sqrt{2}$, (iii) $\kappa = -40$, $\hat{\boldsymbol{\mu}} = \hat{\mathbf{z}}$. Subfigure (a) modified with permission from Günther Eder [22]
.

Another useful scalar summary statistic is the *generalized fractional anisotropy* [20, 21], which is

$$\widehat{\text{GFA}}^{\eta}(\mathfrak{r}_o) = \sqrt{1 - \frac{\left[\hat{F}^{\eta}_{00}(\mathfrak{r}_o)\right]^2}{\sum_{\ell m}\left[\hat{F}^{\eta}_{\ell m}(\mathfrak{r}_o)\right]^2}}. \tag{S89}$$

Although both of these parameters are useful for summarizing the data, strictly speaking neither is estimable since they do not live in the measurement space of the imaging system [15, ch. 15.1.3]. We know that estimates of $\rho$ and GFA can be biased, so we need to be skeptical of any conclusions drawn from them. Averaging over larger spatial regions can reduce (but not eliminate) these biases.

#### 7.5 Peak histograms

**Figure S10** illustrates how we build peak histograms from a list of peaks. Starting with a list of 3D vectors representing peaks, we choose a viewing direction (e.g. the $\hat{\mathbf{z}}$

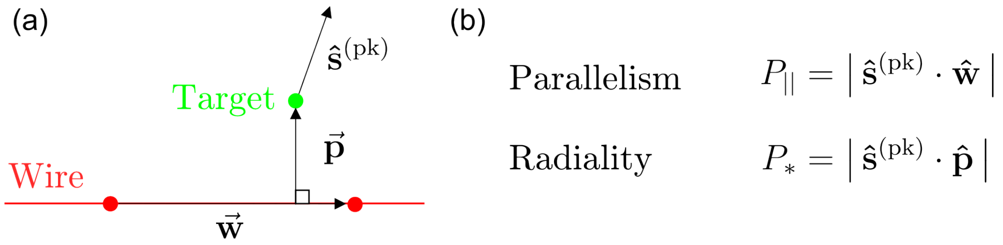


**Fig. S11 Peak metrics.** (a) For each target point (green dot), we find the nearest pair of annotated wire points (red dots). We use these three points to define two vectors: $\vec{\mathbf{w}}$, the vector from one of the wire points to the other, and $\vec{\mathbf{p}}$, the shortest vector from the wire to the target point. Additionally, we have the target point's peak direction $\hat{\mathbf{s}}^{(\mathrm{pk})}$ as the orientation along which the ODF is maximized. (b) We normalize $\vec{\mathbf{w}}$ and $\vec{\mathbf{p}}$ then define a pair of scalar metrics that we call parallelism and radiality.

axis in **main-text Figure 4**), flip the vectors to the upper hemisphere with respect to the viewing direction, project the resulting vectors to a circle by dropping the component along the viewing axis, then increment bins in a polar plot for each peak in the list. We emphasize that the peak histogram depends on the viewing orientation.

Points near the center of the peak histogram represent peaks that have a large component along the viewing axis, while points on the outer rim of the peak histogram represent peaks that lie in the plane perpendicular to the viewing direction. For example, **Figure S10(b)(ii)** shows a histogram with peaks in the $\hat{\mathbf{x}} - \hat{\mathbf{y}}$ plane appearing on the outer rim of the histogram, and peaks with a significant $\hat{\mathbf{z}}$ component appearing closer to the center of the histogram, while **Figure S10(b)(iii)** shows a histogram where all peaks are nearly in the $\hat{\mathbf{x}} - \hat{\mathbf{y}}$ plane and appear on the outer rim of the histogram.

We are representing axes, not vectors, so in-plane peaks can be equivalently represented by points on opposite sides of the peak histogram. This means that a tightly grouped set of nearly in-plane peaks with some peaks above and below the plane normal to the viewing axis will appear as two populations on opposite sides of the peak histograms (see **Figure S10(a)(iv) and (b)** for examples).

Mathematically, the peaks are members of the real projective plane $\mathbb{RP}^2$, and we are drawing histograms on a minimal 2D surface that represents this space.

#### 7.6 Summary statistics with respect to nanowires

In **main-text Figure 6** we measure peak orientations with respect to the nearest nanowires using a pair of metrics we call radiality and parallelism. After annotating the nanowires manually from a separate channel (see **Supplement 4.3**), we calculated each point's peak direction $\hat{\mathbf{s}}^{(\mathrm{pk})}$, nearest wire direction $\vec{\mathbf{w}}$, and the nearest wire's normal direction pointing toward the target point $\vec{\mathbf{p}}$, see **Figure S11(a)**. We use these vectors to calculate parallelism and radiality, see **Figure S11(b)**.

Parallelism and radiality are scalar values between 0 and 1. A parallelism value of 1 (0) indicates a peak direction that is exactly parallel (perpendicular) to the nearest wire, and a radiality value of 1 (0) indicates a peak direction that is exactly parallel (perpendicular) to lines that point radially outwards from the wire.

#### 7.7 Reconstruction and visualization summary

We complete this section by summarizing our reconstruction and visualization algorithms in **Table S4**. We accompany each step with numpy pseudocode to aid implementations, and we highlight the use of `np.einsum` [23], an efficient way to program multidimensional array multiplications inspired by Krister Åhlander's C++ library [24].

### 8 Choosing polarization and tilt samples

In **Supplement 2.5** we described the three polarization-tilt sampling schemes that we used to acquire datasets. In this section we describe how we chose and optimized these sampling schemes.

#### 8.1 Why make six measurements?

Our goal is to choose a set of illumination polarizations $\{\hat{\mathbf{p}}_{j\mathrm{v}}\} \in (\mathbb{S}^2)^N$—$N$ points on the sphere—that allow us to estimate valuable object parameters while keeping $N$ reasonably small so that our acquisition is fast and does not compromise our sample's health.

The central question becomes: what object parameters should we try to estimate? Qualitatively, our goal is to recover as much angular information about our sample as possible, and the transfer functions described in **Supplement 6** provide clear bounds on what we can hope to recover. Specifically, we found that our instrument (and any fluorescence instrument that uses non-saturating single-photon excitation) is band limited to the $\ell \in \{0, 2, 4\}$ spherical harmonics, which suggests a choice of $N = 1 + 5 + 9 = 15$ measurements to attempt to recover all 15 spherical harmonic coefficients in the zeroth-, second-, and fourth-order bands.

However, our instrument only gives us control over the polarizations we use to illuminate our sample, not the polarizations we detect. In **Supplements 6.3 and 6.4** we showed that the angular excitation and detection transfer functions individually transfer angular information from the $\ell \in \{0, 2\}$ bands, and their combination transfers information from the $\ell \in \{0, 2, 4\}$ bands. Since we only have polarization control over the illumination polarization, we can only meaningfully control what information we can collect from the $\ell \in \{0, 2\}$ bands. Since we do not have enough degrees of freedom to completely sample the information in the $\ell = 4$ band, we restrict our attention to the $\ell \in \{0, 2\}$ bands. This narrower goal suggests a choice of $N = 1+5 = 6$ measurements.

We note that polarization control on both the illumination and detection arms can allow measurement of all fifteen terms in the $\ell \in \{0, 2, 4\}$ bands, which is an angular analogue to structured illumination microscopy (SIM). For example, two-dimensional measurements with low-NA (widefield) illumination and high-NA detection results in a $2\mathrm{NA}/\lambda$ cutoff, illumination with sinusoidal patterns created with a high-NA objective and measured with a small-NA detection objective results in an effective $2\mathrm{NA}/\lambda$ cutoff, then combining illumination with sinusoidal patterns created with a high-NA objective and measured with a high-NA detection objective results in an effective $4\mathrm{NA}/\lambda$ cutoff.

#### 8.2 Why use tilting light sheets?

Our first iteration of the instrument described in this paper did not include the tilting degree of freedom, so our accessible illumination polarizations were restricted to a set of two great circles perpendicular to the illumination axes. Mathematically, this design restricted our illumination polarizations to choices that satisfied

$$\hat{\mathbf{p}}_{j\text{v}} \cdot \mathbf{R}_\text{v}\hat{\mathbf{d}}_B = 0. \tag{S90}$$

We attempted to choose six sample polarization illuminations subject to this constraint that would allow us to recover the $\ell \in \{0, 2\}$ spherical-harmonic coefficients, but we found this to be impossible. Our key finding was that the $\ell = 2$ band contains a *null function of our imaging system*, an object that, when added or subtracted from any object, generates identical data. Stated differently, we found that one of our six target parameters was invisible to our instrument.

The null function of the non-tilting design, depicted in **Figure S12**, is the angular distribution

$$f^{(\text{null})}(\hat{\mathbf{s}}_o) = \sin^2\theta\cos\phi\sin\phi, \tag{S91}$$

where $\theta$ is measured from the $\hat{\mathbf{y}}$ axis and $\phi$ is measured from the $\hat{\mathbf{d}}_B$ to the $\hat{\mathbf{d}}_A$ axis in the $\hat{\mathbf{d}}_A - \hat{\mathbf{d}}_B$ plane. We can see that this is a null function by inspection. Any illumination polarization perpendicular to the detection axes will equally excite the positive and negative lobes of the null function, resulting in zero irradiance. Equivalently, if we have an arbitrary ODF and add any multiple of the null function to that ODF, the resulting signal will be unchanged.

In practice, we found that this single null function caused our reconstructions to be difficult to use and interpret. We found that we could not recover all three-dimensional orientations, and our peak estimates would commonly show dramatically incorrect orientations from known samples.

To overcome this limitation we augmented the instrument with the tilting degree of freedom so that the propagation direction of the illumination light sheet was no longer constrained to a single axis. In principle we could tilt the illumination light sheet in any orientation, but most tilting orientations would move parts of light sheet out of the focal plane of the detection objective. Therefore, we tilted the illumination light sheet about the light sheet plane's normal axis. After implementing light-sheet tilting, the polarization samples need to satisfy the looser constraint

$$\hat{\mathbf{p}}_{j\text{v}} \cdot \mathbf{R}_\text{v}\mathbf{R}_{\hat{\mathbf{d}}_A}(\phi)\hat{\mathbf{d}}_B = 0, \tag{S92}$$

where $\mathbf{R}_{\hat{\mathbf{d}}_A}(\phi)$ denotes a rotation about the $\hat{\mathbf{d}}_A$ axis by angle $\phi$ with entries

$$\mathbf{R}_{\hat{\mathbf{d}}_A}(\phi) = \begin{bmatrix} \cos\phi & -\sin\phi & 0 \\ \sin\phi & \cos\phi & 0 \\ 0 & 0 & 1 \end{bmatrix}. \tag{S93}$$

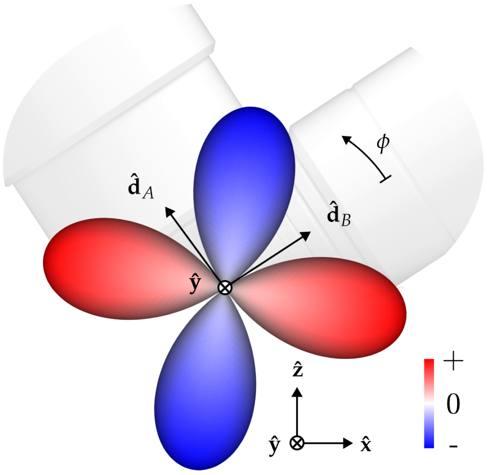


**Fig. S12 Non-tilting design null function.** Without light-sheet tilting, the imaging system has the angular null function depicted here. The null function is described mathematically as $f^{(\text{null})}(\hat{\mathbf{s}}_o) = \sin^2\theta\cos\phi\sin\phi$ where $\theta$ (not shown) is measured from the $\hat{\mathbf{y}}$ axis and $\phi$ is measured from the $\hat{\mathbf{d}}_B$ to the $\hat{\mathbf{d}}_A$ axis in the $\hat{\mathbf{d}}_A - \hat{\mathbf{d}}_B$ plane. The null function along each direction is drawn with radius proportional to $|f^{(\text{null})}(\hat{\mathbf{s}}_o)|$ with negative values colored blue and positive values colored red.

We implemented practical tilt angles that satisfied $|\phi| \lesssim 10°$, a limit set by aberrations, which allows us to eliminate the null function. Notice that our tilting scheme does not change the spatial illumination pattern when light-sheet broadening is negligible, so all of the models we developed in **Supplement 6** still apply to the augmented instrument.

Purely spatial transfer functions have null functions that correspond to zeros in the transfer function. This might suggest that we look for zeros in the spatio-angular transfer function to find null functions, but unfortunately the absence of zeros in the spatio-angular transfer function does not always indicate an absence of null functions. The coefficients of the spherical harmonics change when we choose different spherical coordinate systems (see **Supplement 5.6** to see how the Wigner-D matrices help us find these coefficients under rotations), so null functions will be linear combinations of the spherical harmonics in most coordinate systems. This is the case in this work where we found no zeros in the transfer function, but still found a null function.

We found the null function in **Equation S91** by examining the singular value decomposition of our spatio-angular transfer function $\mathsf{H}_{j\mathrm{v},\ell m}(\mathfrak{v})$, where $j\mathrm{v}$ indexes the rows and $\ell m$ indexes the columns. Every null function has a corresponding zero among the singular values, and we first identified the null function by noticing that we always had zero in our non-tilting singular spectra. In other words, we found that without tilting our spatio-angular transfer functions were at most rank 5, while with tilting we could find spatio-angular transfer functions that were rank 6.

#### 8.3 Optimizing polarization-tilt samples

Finally, we need to choose a set of six samples $\hat{\mathbf{p}}_{j\mathrm{v}} \in (\mathbb{S}^2)^6$ subject to the constraint in **Equation S92**. We chose samples that optimized our abilility to recover all six

$\ell = 0$ and $\ell = 2$ spherical harmonic coefficient for large spatial objects. We start by calculating the entries of the spatio-angular transfer function matrix at $\boldsymbol{\mathfrak{v}} = \mathbf{0}$ for a specific choice of polarization-tilt samples

$$\left(\mathcal{H}_{\hat{\mathbf{p}}_{j\mathtt{v}}}\right)_{j\mathtt{v},\ell\mathtt{m}} = \mathsf{H}_{j\mathtt{v},\ell m}(\mathbf{0}). \tag{S94}$$

This $6 \times 15$ matrix depends on our choice of illumination polarizations $\hat{\mathbf{p}}_{j\mathtt{v}}$ via the angular excitation transfer function, see **Equation S68**. We are only interested in recovering the $\ell = 0$ and $\ell = 2$ spherical harmonic coefficients, so we project this matrix onto that subspace by multiplying with $\mathcal{I}_{15\times 6}$, a $15 \times 6$ matrix of zeros with ones along the upper-left diagonal. Finally, we optimize the condition number of this $6 \times 6$ matrix by solving

$$\underset{\hat{\mathbf{p}}_{j\mathtt{v}}}{\operatorname{argmax}}\, \kappa\left(\mathcal{H}_{\hat{\mathbf{p}}_{j\mathtt{v}}}\mathcal{I}_{15\times 6}\right), \tag{S95}$$

where $\kappa(\mathcal{H})$ is the condition number of $\mathcal{H}$. Optimizing the condition number of this matrix leads to designs where changing each of the input parameters results in a large and independent change of the measured data, which makes the inverse problem maximally invertible and least susceptible to corruption by noise.

We found an approximate solution of **Equation S95** by discretizing the tilt angle into three choices $\mathtt{t} \in \{-1, 0, +1\}$, discretizing the polarizer angle into six choices $p \in \{0, 45°, 60°, 90°, 120°, 135°\}$, then performing a brute-force search across this space of possibilities. Each objective function evaluation required us to compute the condition number of a $6 \times 6$ matrix required $\sim 2$ ms on a single-core machine. With these choices, a brute-force search was feasible because $\binom{36}{6} \approx 2 \times 10^6$ objective function evaluations required about 2 hours.

Our optimized sample with and without light-sheet tilting are shown in **Figure S13**. Without tilting, the optimal samples use equally spaced polarization samples with three samples from each illumination direction, and we named this rank-5 scheme **Six no tilt**. With tilting, the optimal samples are asymmetric samples from each illumination direction, and we named this rank-6 scheme **Six with tilt**.

We also acquired datasets with all possible tilt and polarization settings, a rank-6 scheme we called **All**. See **Table S1** for a summary of our excitation sampling schemes.

### 9 Movies

**Movie M1. GUV fly around.** A spatio-angular reconstruction of a $\sim 6$ $\mu$m-diameter GUV labelled with FM1-43 with (a) ODFs and (b) peak cylinders separated by 390 nm, and (c) a 3D density MIP. As the movie progresses the camera's viewing axis rotates around the object.

**Movie M2. GUV peak slices.** A peak-cylinder reconstruction of a $\sim 6$ $\mu$m-diameter GUV labelled with FM1-43 shown (a) in overview with peak cylinders separated by 390 nm, and (b) a single-slice view where the slice is marked with a grey square in both (a) and (b) and peak cylinders separated by 260 nm. As the movies

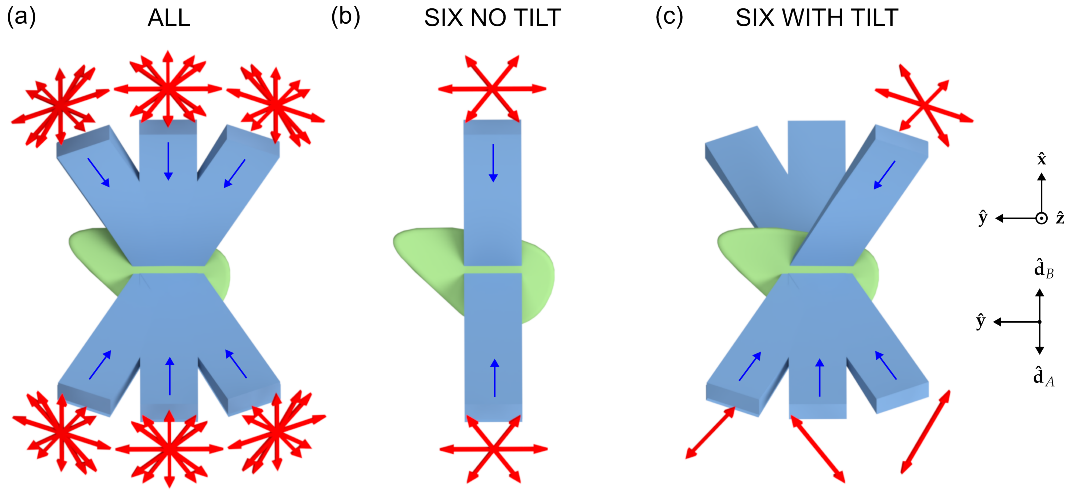


**Fig. S13 Excitation sampling schemes.** These top-down views of tilted illumination light sheets (light-blue rectangles propagating along the dark-blue arrows) and illumination polarization orientations (red arrows) summarize our excitation sampling schemes. Each red arrow has a transverse orientation, tilt, and view and corresponds to a single illumination sample. (a) We started with a complete set of 36 tilt illumination settings then searched from among the six-sample subsets that would optimize the condition number of the imaging system. When we restricted ourselves to samples without tilt, we found the **Six no tilt** scheme (b) three equally spaced polarization orientations for each illumination axis. When we allowed tilting, we found the **Six with tilt** scheme (c) which uses an view-asymmetric combination of polarization and tilt to maximize the condition number.

progresses the highlighted slice sweeps through the object in steps of 130 nm.

**Movie M3. Xylem fly around.** A spatio-angular reconstruction of a xylem cell with its cellulose labelled by fast scarlet with (a) ODFs and (b) peak cylinders separated by 1.3 $\mu$m, and (c) a 3D density MIP. As the movie progresses the camera's viewing axis rotates around the object.

**Movie M4. Xylem peak slices.** A peak-cylinder reconstruction of xylem cell with its cellulose labelled by fast scarlet shown (a) in overview with peak cylinders separated by 1.3 $\mu$m, and (b) a single-slice view where the slice is marked with a grey square in both (a) and (b) and peak cylinders separated by 520 nm. As the movies progresses the highlighted slice sweeps through the object in steps of 520 nm.

**Movie M5. U2OS actin fly around.** A spatio-angular reconstruction of a U2OS cell with its actin labelled by Alexa Fluor 488 phalloidin with (a) ODFs and (b) peak cylinders separated by 260 nm, and (c) a 3D density MIP. As the movie progresses the camera's viewing axis rotates around the object.

**Movie M6. U2OS actin peak slices.** A peak-cylinder reconstruction of a U2OS cell with its actin labelled by Alexa Fluor 488 phalloidin shown (a) in overview with peak cylinders separated by 260 nm, and (b) a single-slice view where the slice is marked with a grey square in both (a) and (b) and peak cylinders separated by 130 nm. As the movies progresses the highlighted slice sweeps through the object in steps of 130 nm.

| Scheme name | $N$ samples | $(p, \mathtt{t}, \mathtt{v})$ samples | Main-text figures |
|---|---|---|---|
| **All** | 42 | `for` $p$ `in` $\{0°, 45°, 60°, 90°, 120°, 135°, 0°\}$:<br> `for t in` $\{-1, 0, +1\}$:<br> `for v in` $\{A, B\}$:<br> $(p, \mathtt{t}, \mathtt{v})$ | 2 |
| **Six no tilt** | 6 | `for` $p$ `in` $[0°, 60°, 120°]$:<br> `for v in` $[A, B]$:<br> $(p, 0, \mathtt{v})$ | 3 |
| **Six with tilt** | 6 | $(0°, +1, A)$<br>$(60°, 0, B)$<br>$(60°, +1, A)$<br>$(120°, +1, B)$<br>$(120°, +1, A)$<br>$(120°, -1, B)$ | 3, 4, 5, 6 |

**Table S1** We acquired data under three different *excitation sampling schemes* named **All**, **Six no tilt**, and **Six with tilt**. Each sampling scheme consists of multiple samples, and each sample is described in the third column in $(p, \mathtt{t}, \mathtt{v})$ notation.

| Symbol | Member of | Description |
|---|---|---|
| $A(\boldsymbol{\tau}, \text{NA})$ | $\mathbb{R}$ | amplitude pupil function |
| $A, B$ | - | view labels |
| $\beta$ | $\mathbb{R}$ | thermodynamic beta $= 1/k_bT$ |
| $\beta_{in}$,$\text{b}_{nn'}$, $\gamma_{in}$ | $\mathbb{R}$ | intermediate electric-field representations |
| $\hat{\mathbf{d}}_A, \hat{\mathbf{y}}, \hat{\mathbf{d}}_B$ | $\mathbb{R}^3$ | unit vectors aligned with detection objectives |
| $\mathscr{D}_v$ | operator | Smoluchowski operator |
| $\nabla$ | operator | gradient |
| $\mathbf{D}$ | operator | generalized diffusion tensor |
| $\Delta^\ell_{mm'}$ | $\mathbb{R}$ | real Wigner D-matrix |
| $f^{(\text{gr})}(\mathbf{r}_o, \hat{\mathbf{s}}_o, t)$ | $\mathbb{L}_2(\mathbb{R}^3 \times \mathbb{S}^2 \times \mathbb{R})$ | ground state spatio-angular density |
| $f^{(\text{ex})}(\mathbf{r}_o, \hat{\mathbf{s}}_o, t)$ | $\mathbb{L}_2(\mathbb{R}^3 \times \mathbb{S}^2 \times \mathbb{R})$ | excited state spatio-angular density |
| $f^{(\text{em})}(\mathbf{r}_o, \hat{\mathbf{s}}_o)$ | $\mathbb{L}_2(\mathbb{R}^3 \times \mathbb{S}^2)$ | spatio-angular emission density |
| $f(\mathbf{r}_o, \hat{\mathbf{s}}_o)$ | $\mathbb{L}_2(\mathbb{R}^3 \times \mathbb{S}^2)$ | spatio-angular Boltzmann density |
| $\mathsf{F}_{\ell m}(\boldsymbol{\mathfrak{v}})$ | $\mathbb{L}_2(\mathbb{R}^3 \times \mathbb{S}^2)$ | spatio-angular Boltzmann spectrum |
| $g_{j\mathtt{v}}(\mathbf{r}_d)$ | $\mathbb{L}_2(\mathbb{R}^3)^N$ | irradiance measurements |
| $G_{j\mathtt{v}}(\boldsymbol{\mathfrak{v}})$ | $\mathbb{L}_2(\mathbb{R}^3)^N$ | irradiance spectrum |
| $\mathscr{G}^{mm'm''}_{\ell\ell'\ell''}$ | $\mathbb{R}$ | Gaunt coefficient |
| $h^{(\text{exc})}_{j\mathtt{v}}(\mathbf{r}_d, \mathbf{r}_o, \hat{\mathbf{s}}_o)$ | $\mathbb{L}_2(\mathbb{R}^3 \times \mathbb{R}^3 \times \mathbb{S}^2)^N$ | shift-variant spatio-angular excitation point-response function |
| $h^{(\text{exc, sp})}_{\mathtt{v}}(\mathbf{r}_d, \mathbf{r}_o)$ | $\mathbb{L}_2(\mathbb{R}^3 \times \mathbb{R}^3)^N$ | shift-variant spatial excitation point-response function |
| $h^{(\text{exc, ang})}_{j\mathtt{v}}(\hat{\mathbf{s}}_o)$ | $\mathbb{L}_2(\mathbb{S}^2)^N$ | angular excitation point-response function |
| $h^{(\text{det})}_{j\mathtt{v}}(\mathbf{r}_d, \mathbf{r}_o, \hat{\mathbf{s}}_o)$ | $\mathbb{L}_2(\mathbb{R}^3 \times \mathbb{R}^3 \times \mathbb{S}^2)^N$ | shift-variant spatio-angular detection point-response function |
| $h_{j\mathtt{v}}(\mathbf{r}_d, \mathbf{r}_o, \hat{\mathbf{s}}_o)$ | $\mathbb{L}_2(\mathbb{R}^3 \times \mathbb{R}^3 \times \mathbb{S}^2)^N$ | shift-variant spatio-angular point-response function |
| $h_{j\mathtt{v}}(\mathbf{r}, \hat{\mathbf{s}}_o)$ | $\mathbb{L}_2(\mathbb{R}^3 \times \mathbb{S}^2)^N$ | shift-invariant spatio-angular point-response function |
| $\mathsf{H}_{j\mathtt{v},\ell\mathtt{m}}(\boldsymbol{\mathfrak{v}})$ | $\text{L}_2(\mathbb{R}^3)^N$ | spatio-angular transfer function |
| $i$ | Z | imaginary unit, electric field component index |
| $j$ | $[1, 2, \ldots, N/2]$ | polarization index, combines $p$ and $\mathtt{t}$ |
| $k$ | $[1, 2, \ldots, R]$ | singular value index |
| $\kappa$ | $\mathbb{R}$ | transition rate |
| $\kappa^{(\text{d})}(\mathbf{r}_o, \hat{\mathbf{s}}_o)$ | $\mathbb{L}^2(\mathbb{R}^2 \times \mathbb{S}^2)$ | spatio-angular decay transition rate |
| $\lambda$ | $\mathbb{R} > 0$ | wavelength |
| $\ell$ | $[0, 1, 2, \ldots]$ | spherical-harmonic band index |
| $m$ | $\mathbb{Z}$ | spherical harmonic intra-band index $-\ell \leq m \leq \ell$ |
| $n_0$ | $\mathbb{R} > 0$ | index of refraction of the medium |
| $n$ | $[0, 1, 2]$ | dipole component index |
| $\mathtt{n}$ | $[1, ...\mathtt{N}]$ | index for spherical visualization directions |
| $\mathtt{N}$ | $\mathbb{Z}$ | number of spherical visualization directions |
| $\eta$ | $\mathbb{R}$ | regularization parameter |
| $p$ | $[0, \pi)$ | transverse-polarization index |
| $\hat{\mathbf{p}}_{j\mathtt{v}}$ | $(\mathbb{S}^2)^N$ | polarization axis vectors |
| $P_\ell(x)$ | $\text{L}_2([-1, 1])$ | Legendre polynomial |
| $\Phi(\boldsymbol{\tau}, r^\perp)$ | R | defocus-phase pupil function |
| $\rho(\mathbf{r}_o)$ | $\mathbb{L}_2(\mathbb{R}^3)$ | spatial labelling density |
| $\mathbf{r}_o$, $\mathbf{r}_d$ | $\mathbb{R}^3$ | 3D coordinate in object and data space |
| $r^\parallel_\mathtt{v}$ | R | view-specific axial coordinate |
| $R$ | $\mathbb{Z}^+$ | rank |
| $\mathbf{R}_\mathtt{v}$ | $\mathbb{SO}(3)$ | view-dependent rotation matrix |
| $\mathbb{R}^N$ | - | $N$D Euclidean space |
| $\hat{\mathbf{s}}_o$ | $\mathbb{S}^2$ | orientation in object space |
| $s_n$ | R | component of $\hat{\mathbf{s}}_o$ |
| $\mathbb{S}^2$ | - | 2D sphere |
| $t$ | $\mathbb{R}$ | time |
| $\mathtt{t}$ | $\{-1, 0, +1\}$ | tilt labels |
| $U_{k,\ell m}(\boldsymbol{\mathfrak{v}})$ | $\text{L}_2(\mathbb{R}^3 \times \mathbb{S}^2)$ | object-space singular vector |
| $V_{k,j\mathtt{v}}(\boldsymbol{v})$ | $\text{L}_2(\mathbb{R}^3)^N$ | data-space singular vector |
| $v(\mathbf{r}_o, \hat{\mathbf{s}}_o)$ | $\mathbb{L}_2(\mathbb{R}^3 \times \mathbb{S}^2)$ | spatio-angular potential |
| $\mathtt{v}$ | $\{A, B\}$ | view labels |
| $\boldsymbol{\mathfrak{v}}$ | $\mathbb{L}_2(\mathbb{R}^3)$ | spatial-frequency coordinate |
| $w_o$, $w_*$, $w$ | R ¿ 0 | beam widths |
| $\hat{\mathbf{x}}, \hat{\mathbf{y}}, \hat{\mathbf{z}}$ | $\mathbb{S}^2$ | unit vectors aligned with coverslip |
| $Y_{\ell m}(\hat{\mathbf{s}}_o)$ | $\mathbb{L}_2(\mathbb{S}^2)$ | real spherical harmonic function |
| $\mathbb{Z}$, $\mathbb{Z}^+$ | - | integers, positive integers |

**Table S2** Table of symbols.

| Pattern | Examples | Description |
|---|---|---|
| Blackboard bold | $\mathbb{R}^3$, $\mathbb{S}^2$, $\mathbb{L}_2$ | mathematical sets |
| Boldface Roman | $\mathbf{r}$, $\mathbf{s}$, $\boldsymbol{\tau}$, $\boldsymbol{\nu}$ | 2D vectors |
| Boldface Fraktur | $\mathfrak{r}$, $\boldsymbol{\nu}$ | 3D vectors |
| Hat on boldface | $\hat{\mathbf{s}}_o$, $\hat{\mathbf{d}}_A$, $\hat{\mathbf{d}}_B$, $\hat{\mathbf{x}}$, $\hat{\mathbf{y}}$, $\hat{\mathbf{z}}$ | unit vector |
| Hat on non-boldface | $\hat{\mathsf{F}}$ | estimate |
| Letter f | $f$, $F$, $\mathsf{F}$ | object properties to be estimated |
| Letter g | $g$, $G$ | measurements |
| Letter h | $h$, $H$, $\mathsf{H}$ | instrument-response functions |
| Subscript $d$ | $\mathfrak{r}_d$ | detector coordinate |
| Subscript $o$ | $\mathfrak{r}_o$, $\hat{\mathbf{s}}_o$ | object-related coordinate |
| Superscript $\perp$ | $\mathbf{r}^{\perp}$ | transverse coordinate perpendicular to the optical axis |
| Superscript $\parallel$ | $r^{\parallel}$ | axial coordinate parallel to the optical axis |

**Table S3** Notation patterns, subscripts, and superscripts.

| Steps | Symbols | `numpy` pseudocode |
|---|---|---|
| *Precalculations* | | |
| Calculate the model in a compact basis | $\mathsf{H}_{j,\ell m}(\mathfrak{v})$ | `H.shape -> (100, 100, 100, 15, 6)` |
| Calculate the SVD | | `u, s, v = np.linalg.svd(H, full_matrices=False)` |
| ... object space singular functions | $U_{k,\ell m}(\mathfrak{v})$ | `u.shape -> (100, 100, 100, 15, 6)` |
| ... singular values | $\sqrt{\mu_k(\mathfrak{v})}$ | `s.shape -> (100, 100, 100, 6)` |
| ... data space singular functions | $V_{k,j\mathrm{v}}(\mathfrak{v})$ | `v.shape -> (100, 100, 100, 6, 6)` |
| *Reconstruction* | | |
| Collect data | $g_{j\mathrm{v}}(\mathfrak{r}_d)$ | `g.shape -> (100, 100, 100, 6)` |
| DFT | $G_{j\mathrm{v}}(\mathfrak{v})$ | `G = np.fft.fftn(g, axes=(0,1,2))` |
| Choose $\eta$ and regularize singular values | $\sigma_k(\mathfrak{v}) = \frac{\sqrt{\mu_k(\mathfrak{v})}}{\mu_k(\mathfrak{v})+\eta}$ | `sr = s/(s**2 + eta)` |
| Estimate $\hat{\mathsf{F}}^{\eta}_{\ell m}(\mathfrak{v})$ | $\sum_k \sigma_k(\mathfrak{v}) U_{k,\ell m}(\mathfrak{v}) \ldots \sum_j V_{k,j\mathrm{v}}\mathfrak{v}) G_{j\mathrm{v}}(\mathfrak{v})$ | `F = np.einsum('xyk,xysk,xyjk,xyj->xys', sr, u, v, G)` |
| Inverse DFT then save | $\hat{F}^{\eta}_{\ell m}(\mathfrak{r}_o)$ | `Fr = np.fft.ifftn(F, axes=(0,1,2))` |
| *Visualization* | | |
| Choose spherical points | $\{\hat{\mathbf{s}}_{o,\mathrm{n}}\}$ | `sp.shape -> (500, 3)` |
| Calculate SH to ODF coefficients | $Y_{\ell m}(\hat{\mathbf{s}}_{o,\mathrm{n}})$ | `Y[n,s] = sphharm(s2l(s), s2m(s), sp[n,:])` |
| Choose mask (e.g. density > 0.5) | $\{\mathfrak{r}_{o,\mathrm{n}}\}$ | `mask = Fr[:,:,0] > 0.5` |
| Orientation distribution functions (ODFs) | $\hat{f}^{\eta}(\mathfrak{r}_{o,\mathrm{n}})$ | `ODF = np.einsum('ns,is->ni', Y, Fr[mask])` |
| Peak directions | $\hat{\mathbf{s}}^{\eta,(\mathrm{pk})}(\mathfrak{r}_{o,\mathrm{n}})$ | `np.amax(ODF, axis=0)` |
| Density | $\hat{\rho}^{\eta}(\mathfrak{r}_o)$ | `Fr[:,:,0]` |
| Generalized fractional anisotropy | $\widehat{\mathrm{GFA}}^{\eta}(\mathfrak{r}_o)$ | `np.sqrt(1 - (Fr[:,:,0]**2/ np.sum(Fr**2, axis=-1)))` |

**Table S4** Summary of reconstruction and visualization algorithms with pseudocode implementations.
